# Time-resolved, integrated analysis of clonally evolving genomes

**DOI:** 10.1101/2021.10.08.463633

**Authors:** Carine Legrand, Ranja Andriantsoa, Peter Lichter, Frank Lyko

**Affiliations:** Division of Epigenetics, DKFZ-ZMBH Alliance, German Cancer Research Center, 69120 Heidelberg, Germany; Université Paris Cité, Génomes, biologie cellulaire et thérapeutique U944, INSERM, CNRS, F-75010 Paris, France; Division of Molecular Genetics, German Cancer Research Consortium (DKTK), German Cancer Research Center (DKFZ), Im Neuenheimer Feld 280, 69120 Heidelberg, Germany; Molecular Precision Oncology, National Center for Tumor Diseases, Im Neuenheimer Feld 460, 69120 Heidelberg, Germany

## Abstract

Clonal genome evolution is a key aspect for parthenogenetic species and cancer. While many studies describe precise landscapes of clonal evolution in cancer, few determine the underlying evolutionary parameters from molecular data, and even fewer integrate theory with data. We derived theoretical results linking mutation rate, time, expansion dynamics, and clinical parameters. We then inferred time-resolved estimates of evolutionary parameters from mutation accumulation, mutational signatures and selection. Using this framework, we traced the speciation of the clonally evolving marbled crayfish population to a time window between 1986 and 1990, which is consistent with biological records. We also used our framework to analyze a published dataset of glioblastoma samples, which identified tumor expansion patterns, cell survival at resection, and selective forces as important factors for tumor development. In conclusion, our framework allowed a time-resolved, integrated analysis of key parameters in clonally evolving genomes, and provided novel insights into the evolutionary age of marbled crayfish and the progression of glioblastoma.

## Introduction

The evolution of genomes is shaped by many factors, among which the random accumulation of mutations over time plays a fundamental role [1,2]. Far from being homogeneous, the probability of a mutation depends on many factors such as the genomic location [3], mutator alleles, local nucleotide context or mutagenic exposures [4]. Other genomic modifications include recombination in sexual reproduction, copy number variants and genomic rearrangements, gene transfers and hybridization. The capacity of any genomic modification to be inherited is partly stochastic, for instance through genetic drift [5], but can be favored or disfavored by positive or negative selection. Genome evolution was historically observed through the analysis of phenotypes [6], and can now be determined more precisely using high-throughput sequencing in parallel with experimental or cohort settings, such as mutation accumulation experiments, or the analysis of genetic trios [7,8].

Clonal genome evolution is shaped by a more limited set of mechanisms. Mutation rate, growth and variant frequencies are key parameters, which determine the speed of evolution, and which function under the influence of selection pressure. Due to its particular mode of asexual reproduction, the marbled crayfish (*Procambarus virginalis*) represents an ideal animal model to study clonal genome evolution [9]. The animals are currently colonizing diverse habitats in a process that is associated with emerging genetic differentiation [10]. Interestingly, marbled crayfish appears to be an evolutionary young species, as their first emergence can be traced back to a specific event in 1995 [11]. If confirmed, this exceptionally young evolutionary age would represent a highly distinctive feature of the model system.

Clonal genome evolution also plays an important role in cancer formation. Cancer is a disease of clonal evolution, characterized by the accumulation of somatic mutations into a pathogenic tumoral genome. Several authors have described the critical role of mutational patterns and selection in cancer [1,3,12], while neutral evolution is still debated [13,14]. In glioblastoma, the analysis of tumor trajectories revealed a tumor initiation years before diagnosis [15]. Consequently, it would be of great interest to infer evolutionary parameters over the course of tumor progression. In this study, we aimed to develop an integrated analysis of clonal genome evolution. To this end, we reformulated the dependence of mutation accumulation on variant allele frequency, and used this formulation to determine the links between the mutation rate, growth and survival rates. We further integrated these parameters with selection estimates, obtained from the non-synonymous to synonymous ratio. Finally, we integrated time estimates in our model, based on clock-like mutational signatures. We applied our approach to the clonally evolving marbled crayfish. We provided a detailed view of mutation accumulation, selection, and time, and estimated the time of speciation. We further applied our framework to clonal genome evolution in cancer, using recently published samples of primary and recurrent glioblastoma [15].

## Results

The genetic near-monoclonality of the marbled crayfish population [9,10] establishes this species as an excellent model system for studying clonal genome evolution. In order to assess the mutation rate of the *P. virginalis* genome, we used whole-genome sequencing of a line of direct descendants from our laboratory colony of *P. virginalis*, that were sampled over a period of seven years (Fig 1A). The mutation rate was calculated as the average number of de novo mutations in animals 34 and 35 as compared to animal 1, per nucleotide and per year. From these samples, we obtained a mutation rate equal to *μ*=3.51 • 10^*−*8^ /*n t* / *y* (95% confidence interval (CI): [1.67 •10^*−*8^ ; 5.35• 10^*−* 8^]/ *nt* / *y*). This mutation rate of *P. virginalis* is comparable to known mutation rates from other arthropods (Fig 1B).

**Fig 1.**
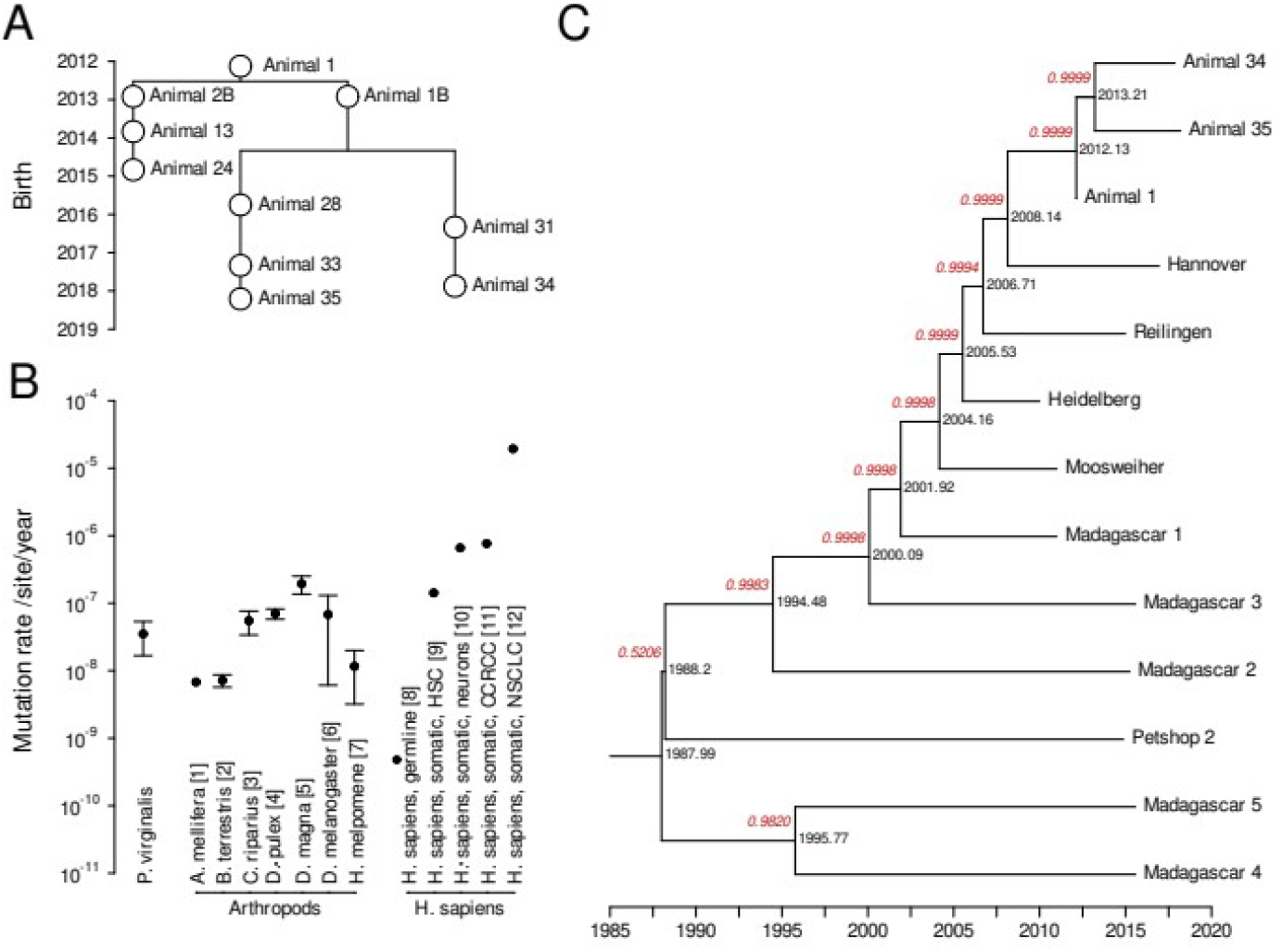
Mutation rate of *P. virginalis* and coalescent. (A) Genealogy of laboratory animals, with sequenced animals marked in grey. (B) Mutation rate in *P. virginalis*, in other arthropods and in *Homo sapiens* [16–27]. HSC: Hematopoietic Stem Cells, CCRCC: Clear Cell Renal Cell Carcinoma, NSCLC: Non-Small Cell Lung Carcinoma. Error bars correspond to reported 95% confidence intervals. (C) Coalescent tree based on a constant mutation rate and sequences of sampled animals. The posterior probability of each branch is indicated in red.

Based on this mutation rate, we made an evaluation of the evolutionary age of *P. virginalis*, using a Markov Chain Monte Carlo with Bayesian evolutionary analysis [28] on whole-genome sequencing datasets from 13 animals (Fig 1C). We generated 10 million states, which allowed convergence of the sampled states, and led to adequately large effective sample sizes (see Methods for details). The resulting coalescent tree showed that animals 1, 34 and 35 correctly clustered together, as well as animals from German wild populations (Hannover, Reilingen, Moosweiher) and from the likely foundational laboratory lineage of the German wild populations (Heidelberg). Furthermore, samples from Madagascar formed a separate branch. Interestingly, Petshop 2 [9] was nested in the branch of animals from Madagascar. This is consistent with the notion that the Malagasy population was founded by an animal that was originally obtained from a German pet shop. Posterior probabilities (Fig 1C, red annotations) indicate highly probable branching for all but the top coalescent event, which has 0.5206 probability. From this tree, the most recent common ancestor of the 13 animals occured in 1988.0 (95% CI: [1986.1 ; 1989.8]). This is anterior, and therefore broadly consistent, with the first documented appearance of *P. virginalis* in 1995 [11].

We next modeled mutation accumulation under a fast growth scenario. This included the patterns of mutation accumulation as a function of time, the dependence of allele frequency on time, and the number of mutations as a function of allelic frequency. We found that mutation accumulation gave information on the mutation rate, but not on selection (S1 File, Eq. 12). This provided the rationale for examining the dynamics of the mutation rate in *P. virginalis* using the *M(1/f)* curve, where *M* is the number of mutations and *f* is the allelic frequency (Fig 2A). The resulting curve suggested that the mutation rate changed over time, with 4 phases delineated by a segmented regression (Fig 2A; *p*=0.06). The mutation rate was reduced in phase 3, as compared to phases 1 and 2, and increased in phase 4 (Fig 2A). Under our model, selection *s* is not observable using *M(1/f)* (S1 File, Eq. 12). We therefore used the ratio of non-synonymous to synonymous mutations to estimate *s* (Fig 2B). The resulting values were close to unity, for phases 1 and 2, suggesting the absence of selection. During phases 3 and 4, we detected *s* values >1, and <1, suggesting phases of positive and negative selection respectively, in the more recent past of *P. virginalis*.

**Fig 2.**
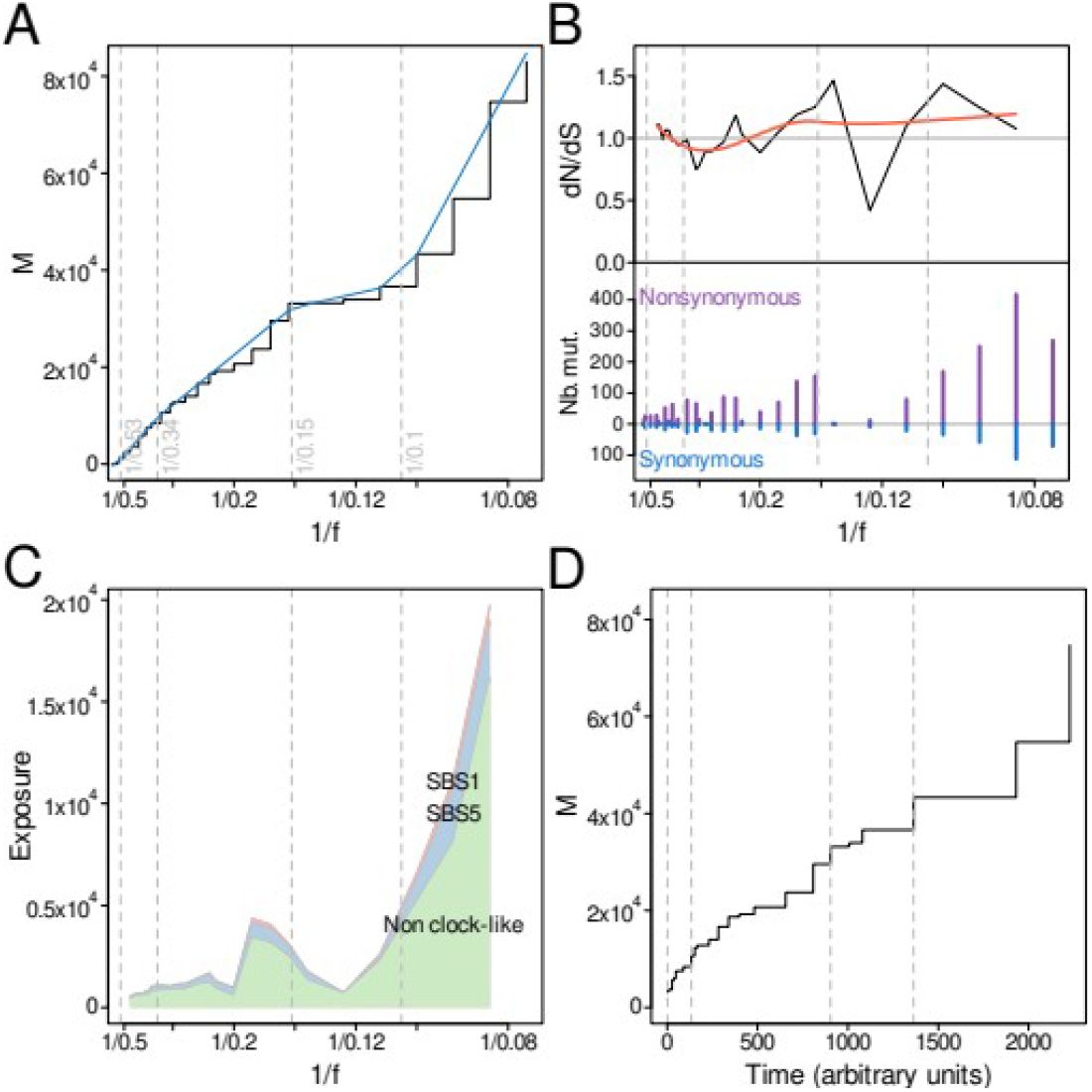
Mutation accumulation, selection and time course of *P. virginalis* genome evolution. (A) Mutation accumulation as a function of the inverse allele frequency *1/f* (black) and phases from automated segmentation (breakpoints in grey, segments in blue). (B) Non-synonymous to synonymous ratio (*dNdS*). The smoothed ratio is shown in red. (C) Comparison of clock-like and non-clock-like mutational signatures. (D) Mutation accumulation as a function of time. Smoothened mutation accumulation is shown in red.

In order to obtain time-resolved estimates, we then used previously established clock-like mutational single-base signatures (SBS1 and SBS5) [29–31] as a proxy for the time course of mutation accumulation (Fig 2C). We further assumed that the arrow of time from past to present corresponds to the arrow of increasing *1/f*. To obtain a time course in arbitrary units, we calculated the integral of the clock-like components of mutation accumulation (Fig 2D, Methods). According to the mathematical model, the slope of this curve is proportional to the mutation rate as a function of time (S1 File, Eq. 12). The results (Fig 2D) showed that this mutation rate exhibited less variation than the mutation rate per division (Fig 2A). Because the temporal and per-division mutation rates differ essentially by the growth rate, this might indicate fluctuations in the growth rate of the marbled crayfish population. As a whole, our analyses suggested distinct phases, detected significant variations of evolutionary parameters in the recent past of *P. virginalis*, and allowed to trace its speciation to a time point that is consistent with biological records.

In *P. virginalis*, we developed a framework to analyse the evolution of a clonal genome, which is driven by germline mutations. This framework can in principle also be applied to analyse the clonal evolution of a tumor genome, which is driven by somatic mutations. Since glioblastoma is a high grade tumor with systematic recurrence and poor patient survival, a better understanding of evolutionary parameters is important. We therefore applied our framework to a published set of whole-genome sequencing data of primary and recurrent glioblastoma tumors [15].

This study also estimated the age of primary tumors, allowing further data integration. Based on the curve *M(1/f)*, we generated mutation rate profiles (Fig. 4A, see S1 Fig A for individual samples), which we further segmented into phases (Fig. 4A, p < 2.2×10^−16^). The results indicated distinct variations in the mutation rate in primary and recurrent samples (Fig 3; S1 Table). In the exemplary sample 1 in Fig 3A, the segmentation separates 5 phases significantly. After we excluded the outermost phases 1 and 5, where changes in mutation frequencies are more likely to be influenced by artifacts, the mutation rate per division decreased steadily in phases 2-4.

**Fig 3.**
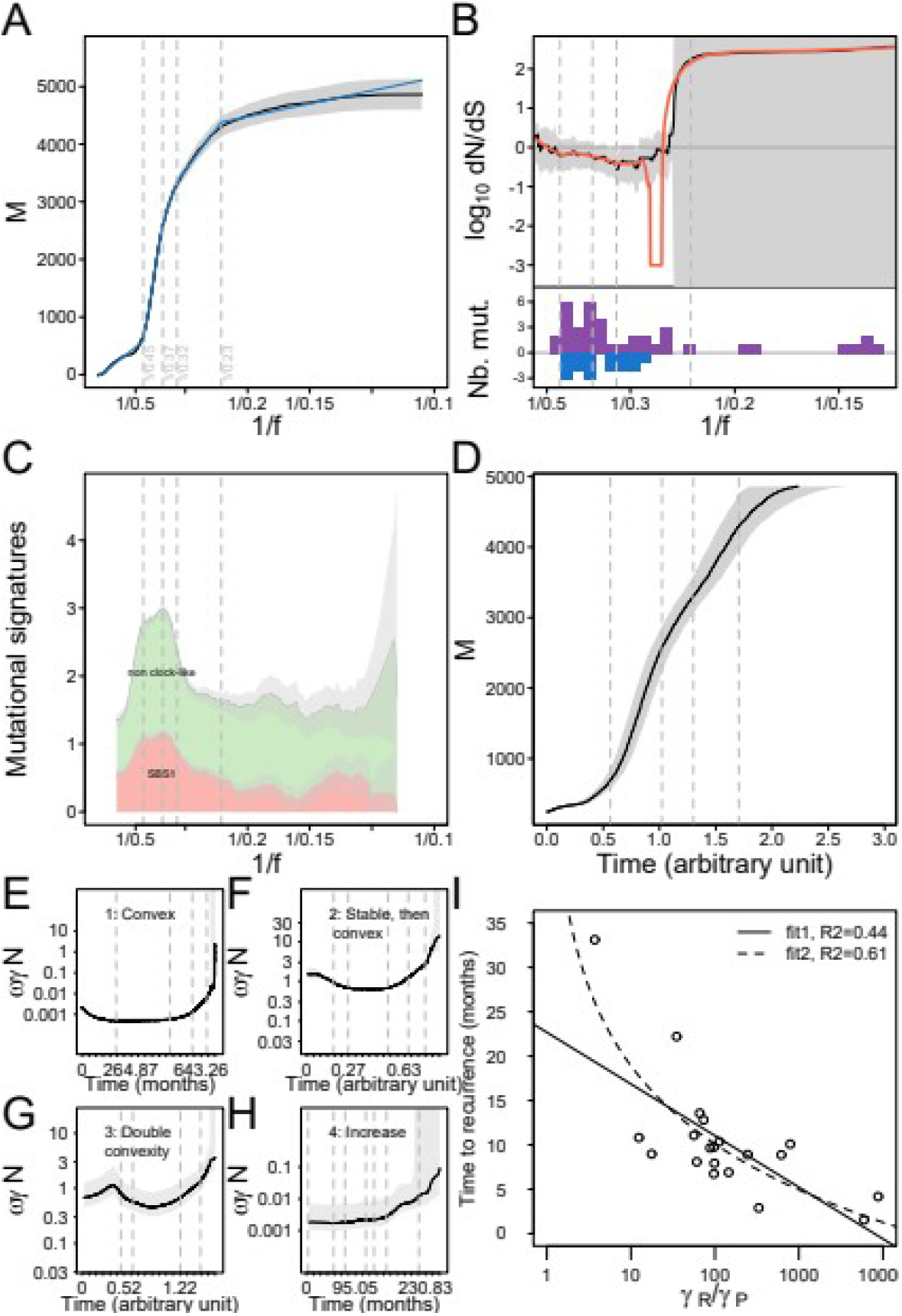
Mutation accumulation, selection and time dynamics of a representative glioblastoma tumor (patient 1, primary tumor), expansion patterns and survival ratio. (A) Mutation accumulation as a function of the inverse allele frequency *1/f* (black) and phases from automated segmentation (breakpoints are indicated as dashed vertical lines, segments are indicated in blue). The confidence band at 95% level is shown in grey. (B) Non-synonymous to synonymous ratio *dNdS*. In the lower inest, purple and blue bars show non-synonymous and synonymous mutations, respectively. The smoothened ratio is shown in red. (C) Clock-like and non-clock-like mutational signatures. (D) Mutation accumulation as a function of time. (E-H) Stratification of expansion curves *ωγN* into 4 subtypes: 1: convex, 2: horizontal then convex, 3: complex, 4: increase (showing exemplary primary tumor samples GBM30 (GlioBlastoma Multiforme, sample number 30), GBM26, GBM17, GBM14, respectively). (I) Dependence of time to recurrence on the *γ*_*R*_*/γ*_*P*_ ratio. Fit1 corresponds to a linear regression of time versus log_10_(*γ*_*R*_*/γ*_*P*_), with intercept=19.511 (standard error SE=2.544) and slope=-5.819 (SE=1.455), fit2 corresponds to a linear regression of time versus log_10_(log_10_(*γ*_*R*_*/γ*_*P*_)), with intercept=18.922 (SE=1.806) and slope=-29.321 (SE=5.285).

We next looked at selection using the *dN/dS* ratio. Taking confidence bounds into account, the results were compatible with neutral selection for most tumors (Fig. 4B, S1 Fig B per sample). However, 11 primary tumor samples showed evidence of negative selection during intervals, for instance sample 35 (S1 Fig B for sample 35). We also observed evidence for positive selection in two primary tumor samples (Samples 2 and 7, S1 Fig B). Interestingly, 7 out of 9 recurrent tumor samples underwent prolonged phases of negative selection (for example, sample 4, 1/f in [1/0.5; 1/0.1], S1 Fig B), while 2 samples exhibited short phases of negative selection. No recurrent tumor sample showed any significant phase of positive selection.

We next determined the timeline of tumor evolution, by comparing the prevalence of the stable, clock-like SBS1 signature to the other, non-clock like SBS signatures (Fig 3C, see S1 Fig for individual samples). This includes SBS5, which is non clock-like in glioblastoma according to [29]. This was confirmed by low correlation coefficients between SBS1 and SBS5 in the samples under study (ρ_P_=-0.060 +/- 0.143 in primary tumors; ρ_P_=0.067 +/- 0.136 in recurrences). Using the information on the clock-like signature SBS1, and using equation (9) (Methods), we reconstructed *M* as a function of time (Fig 3D, S1 Fig D per sample), in arbitrary units. The slope of this curve is proportional to the mutation rate per time unit. Similar to the mutation rate per division in Fig 3A, the mutation rate per time unit decreased from phase 2 to phase 4, though less markedly. Furthermore, similar to the situation for *P. virginalis*, the difference observed in sample 1 might indicate fluctuating growth during phases 2-4. More specifically, differences between per-division and temporal mutation rates corresponded in our model to the growth rate, the survival rate, and the number of cells (S1 File, Eq. 8).

We next asked if these terms could be used to characterize the set of 42 primary and recurrent tumor pairs. Using temporal and per division mutation rates, we could reconstruct these terms, which correspond to the product *ωγN* (growth rate *ω*, tumor cell survival rate *γ* and number of cells *N*), and which we termed expansion parameters. We examined the corresponding curves and found that they allowed us to classify the tumors into four subtypes: (1) convex, (2) stable then convex, (3) double convexity and (4) increase (Fig 3E).We then looked at a possible association between the patterns of the *ωγN* curve in the primary tumors, and the time to the recurrence, but the results were inconclusive (*p*=0.0367, *p*_adj_=0.0734, n=20). However, the time difference between the resection of the primary tumor and the resection of the recurrence is known for a subset of samples, and the age of tumors was estimated previously [15]. This allowed us to transform the time course from arbitrary units into real units (Methods, Eq. 2, S2 Fig). Furthermore, we extended our modelling to be able to express the transition from the primary to the recurrent tumor (Methods, Eq. 4). With this, we could determine the tumor survival ratio from time estimates. Using the previously established values of 2 years and 7 years [15] as the lowest and highest limits for the time course of the primary tumor, we could determine a range for the value of the tumor survival ratio *γ*_*R*_*/γ*_*P*_ for each individual sample (Methods, Eq. 8, S2 Table). As a result, the lowest value of the ratio *γ*_*R*_*/γ*_*P*_, corresponding to a tumor emergence about 2 years before diagnosis, was always higher than 1 (median =27.8 [17.4; 54.0] for the lower bound, median=97,5 [60.9; 189.0] for the upper bound, n=20 samples). These results indicated that tumor cell survival was higher at the start of the recurrence than at the end of the primary tumor growth. Not surprisingly, *γ*_*R*_*/γ*_*P*_ ratios were associated with the time to recurrence (Fig 3F, *p*_adj_=1.258×10^−3^ and *p*_adj_=8.649×10^−4^), with higher *γ*_*R*_*/γ*_*P*_ ratios corresponding to shorter time to recurrence. Collectively, these results uncover substantial variations of evolutionary parameters among glioblastoma samples, and provide an improved understanding of growth and survival in tumor subtypes.

## Discussion

In this study, we presented an integrated framework to analyse the evolution of clonally evolving genomes. We first determined the mutation rate of *P. virginalis* to be 3.51×10^−8^/nt/y, which is comparable to the mutation rates of other arthropods. We detected separated evolutionary phases, during which the mutation rate varied significantly, and which suggested selection events in the recent past of the species. The variations of the mutation rate, observed as well in the tumor samples, support the argument that the mutation rate should not be considered constant [1,32,33]. Known mechanisms can explain these variations, including a temporarily more pronounced effect of error-prone mechanisms, hypoxia-induced mutagenesis, or transcription-associated mutagenesis [1,34]. We further noted that the temporal mutation rate seemed more stable than the per-division mutation rate, and that the difference, according to our modelling approach, could be explained by variations of growth. Finally, we traced the speciation of *P. virginalis* to 1988 (95% confidence limits: [1986 ; 1990]), in agreement with first reports of this animal in 1995 [11], and consistent with a very young evolutionary age of the species.

In tumor samples, our approach allowed a single patient-level analysis of evolutionary parameters, and similarly revealed the presence of different phases, variations of the mutation rate, and a few significant events of selection. Utilizing the difference between temporal and per-division mutation rate, we could stratify the samples into 4 subtypes. Building on previously estimated tumor age, we could also derive the survival ratio for tumor cells in the recurrence, relatively to the primary tumor. We found that tumor cells survive better at the start of the recurrence, though with important variations. This supports the notion that tumor regrowth can be more aggressive after surgical resection of primary glioblastoma tumors [35,36], possibly because resection-induced astrocyte injury can support faster growth [36], or because the tumor microenvironment can promote tumor regrowth [37–39]. Conversely, a stronger immune response might prevent tumor regrowth.

In conclusion, this integrated analysis of mutation accumulation, dNdS ratio and mutational signatures provided a detailed landscape of evolutionary parameters, and delivered important insights for two paradigms of clonal genome evolution, demonstrating its potential for further applications.

## Materials and Methods

### *Procambarus virginalis* samples

Freshwater crayfish samples from [9] were used. Additionally, samples from Madagascar 1 sample and Moosweiher sample were resequenced (S3 Table). Animal 1 corresponds to the lab strain, acquired from a pet shop. New genomic DNA samples were taken from animal 34 and animal 35, which, as animal 1, also correspond to lab strains animals, and which are direct offsprings of animal 1. These new samples were prepared and submitted for whole genome sequencing following the protocol already described. The genealogy and birth date of animals were retrieved from laboratory records and field records (S3 Table).

Sequence data was trimmed using Trimmomatic v0.32 (settings: LEADING:3 TRAILING:3 SLIDINGWINDOW:4:20 MINLEN:40, adapter sequence: TruSeq3-PE). Next, trimmed data was mapped to Pvir genome assembly v04 (https://www.ncbi.nlm.nih.gov, Bioproject Accession: PRJNA356499), using Bowtie2 (v2.2.6, setting: --sensitive). Aligned reads were sorted, cleared from duplicates, sorted and indexed using samtools. Subsequently, variant calling was performed using Freebayes v0.9.21-g7dd41db (parameters: --report-all-haplotype-alleles -P 0.7 -p 3 --min-mapping-quality 30 --min-base-quality 20 --min-coverage 6 --report-genotype-likelihood-max).

### Glioblastoma Multiforme samples

The glioblastoma primary and recurrent tumor samples correspond to the WGS cohort already described in [15]. In particular, summary information can be found in supplementary table 1 of [15]. After approval of the research project, access to the SNP data of primary and recurrent tumor samples, as well as time to recurrence when available, was granted.

### Mutation rate of *P. virginalis*

Mutation accumulation between animals 1 and descendant animals 34 and 35 was used to calculate the mutation rate. SNP variants were examined in terms of quality and coverage. Variants with quality≥35, coverage≥50 and ≤200 were retained for the main estimate of the mutation rate.

Coverages 200 and higher exhibited altered SNP distribution and were thus excluded because possibly corresponding to a distinct part of *P. virginalis* genome (possibly highly repetitive and variable domains). Subsequently, the mutation rate per nucleotide per year was calculated as the count of biallelic mutated nucleotides in animal 34 (respectively, animal 35) as compared to animal 1, divided by the count of nucleotides in the triploid genome of *P. virginalis* (N=10.5×10^9^), and divided by the time (5.75 and 6.08 years), between the birth dates of animal 1 and 34 (respectively, birth date of animal 35). We assumed that counts of new mutations follow a Poisson distribution, and with this we determined the standard deviation on the count of mutations observed (genotyping uncertainty). Second, we assumed that the standard deviation for the date of animal birth equated to a third of the total uncertainty on time of birth. Third, we calculated the standard deviation between mutation rates for animal 34 and animal 35 (biological variability). Finally, we took the total standard deviation as the quadratic sum of these three components (assuming that the different sources of variability follow a normal distribution)

### Coalescent time

Time to most recent common ancestor for *P. virginalis* samples was determined using Bayesian evolutionary analysis by sampling trees (BEAST v1.10.4 [28]). Mutation data with quality >35 and coverage depth >15 was used in this analysis (a coverage cutoff of 25 was not justified here because samples other than animals 1, 34 and 35 possessed a notably lower average sequencing depth). Samples birth dates were used as tip dates. Further BEAST parameters used were: simple substitution model with estimated base frequencies, strict clock, skyride coalescent prior. The length of chain for the Markov chain Monte Carlo was 10M. These parameters were built into the BEAST input file, using the utility BEAUTi. The outputs were analyzed using the utility TRACER. In particular, convergence was read from the sampled states curves of the different parameters, and effective sample sizes were adequately >100 (2984 or more), indicating sufficiently decorrelated sampled states.

### Study of mutation accumulation

An infinitesimal increment of mutations was defined from the mutation rate, ploidy, cell survival, growth and number of cells (see S1 File for a detailed description). We stratified this expression for each subclone. We have then determined the relationship between the observable allele frequency of a mutation, which is the one obtained after sequencing and SNP calling, and the features of the subclone where this mutation appeared. Next, we have determined a formula for the increment of the number of cells *dN* and for the increment of inverse allele frequency *d(1/f)*. For this latter increment, we have made the assumption that ploidy was constant. We could then deduce the mutation accumulation *dM* as a function of inverse allele frequency *1/f*, in each subclone *i*. Finally, the equation for mutation accumulation over all subclones *dM* was obtained by summing the individual contributions *dM* of each subclone. Mutation accumulation *M(1/f)*, which slope corresponds to *dM/d(1/f)*, was plotted from SNP data (filtered by quality phred score QUAL ≥30, depth ≥10, depth of alternate allele ≥3), and from the corresponding allele frequencies. For uncertainties, we used bins of +/-0.25 over 1/f. In each bin, the mutation count was subjected to bootstrap resampling, which yielded a bootstrap distribution. The confidence interval at 95% was taken as the interval bounded by the 2.5% and 97.5% quantiles of the bootstrap distribution.

### Mutation annotation and dNdS ratio

Mutations were annotated as synonymous or non-synonymous (including splice or stopgain mutations) using SNPdat v1.0.5. The *dNdS* ratio was calculated as the quotient of non-synonymous mutations by synonymous mutations in a sample, divided by the average quotient in the full genome. The average quotient of non-synonymous to synonymous in humans was taken equal to 3.34951759 (hg19). Uncertainties were determined by bootstrap resampling of mutations, and calculations of the dNdS ratio for each bootstrap sample.

### Mutational signatures

Mutational signatures for human subjects were downloaded from the COSMIC database (https://cancer.sanger.ac.uk/signatures/ ; version 3.1 as of 11.08.2020). Mutation data was binned using a bin half-width of 0.5 on the inverse allele frequency. Exposure of binned data was determined using R 3.5.2 with package YAPSA (version 1.8.0). Uncertainty on mutational signatures was determined by bootstrap resampling of mutations and generation of the binned data and YAPSA exposures on the resampled data. We have used 1000 bootstrap replicates as a compromise between an ideally larger (1M) number of replicates, and reasonable computing time. Large mutation sets (>100,000 mutations) were subsampled to 50,000-60,000 mutations for the bootstrap analysis. Mean, median, percentiles and 95% confidence bounds were determined using the resulting bootstrap distribution.

### Time course

We made the assumption that clock-like mutational signatures SBS1 and SBS5 were a surrogate indicator for time (SBS1 only for glioblastoma, in agreement with Alexandrov et al. 2015). We further assumed that the arrow of time, could be identified with the arrow of inverse allele frequency *1/f*. Under these assumptions, an increment of time can be deteremined in arbitrary units, by integrating *θ*, which denotes the exposure to clock-like mutations SBS1 or SBS5, over an increment of inverse allele frequency *1/f*. Since the exposure *θ* is also proportional to the number of cells in the tumor, it is necessary to normalize *θ*, by dividing its value by the number of cells *N*. Since *N* is proportional to *1/f* under some assumptions (S1 File), *N* can be replaced, up to an unknown constant, by *1/f*. This yielded the formula for determining time *t* in arbitrary units, over an interval of time which is unknown, but identifiable with an interval over inverse allele frequency *1/f*:

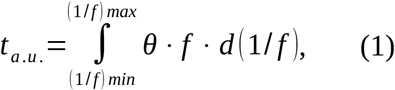

where *t*_*a*.*u*._is the time in arbitrary units (a.u.), and *θ* corresponds to the exposure to clock-like mutational signatures. By computing *t*_*a*.*u*._ at all values of *1/f*, we obtained a vector of values *T*_*a*.*u*._ for the time in arbitrary units, from its minimum value, min(*T*_*a*.*u*._) (0 by definition), to its maximum value, max(*T*_*a*.*u*._) (which corresponds to integration from *(1/f)*_*min*_ to *(1/f)*_*max*_. Confidence limits at 95% for time were calculated using the confidence bounds for mutational signature SBS1, taken as exposure *θ*.

### Time calibration in recurrent glioblastomas

In a subset of glioblastoma samples, we additionally know the time-to-relapse, denoted T, in months. We thereby identify the time course of the relapse [0 ; T] in months, with the time course in arbitrary units [0 ; max(T_a.u._)] (see the preceding paragraph “Time course” for the calculation of T_a.u._). To obtain *t*, the time in real units at any instant *t* in the interval [0 ; T], *t*_*a*.*u*._ is multiplied by the scaling factor T/max(T_*a*.*u*._):

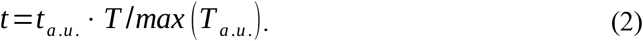

### Time propagation from primary to recurrent glioblastomas

To obtain a link between the time course in the primary tumor, and the time course in the recurrent tumor, we studied the ratio of mutation accumulation between the end of primary tumor (subscripted ‘P’, taken as the last 5% time points) and start of recurrence (subscript ‘R’, first 5% time points), over the entire tumor. This is justified by the observation that the passage from the primary tumor to the recurrence corresponds to the instant of primary tumor resection. Using Eq. 1 (S1 File), this ratio could be written as follows:

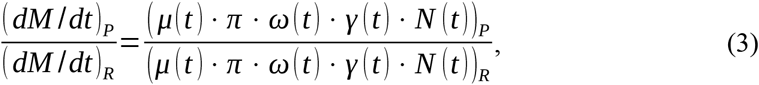

where subscript *i* per subclone is not used, because we considered here that evolution parameters are taken over the whole tumor. The constant term *π* can be normalized out of this ratio. Further, we have assumed that the mutation rate *μ* and division rate *ω* stay constant over this short period, because they are intrinsic features of the tumor cells. However, the count of tumor cells *N(t)*, and the tumor cell survival rate *γ(t)* could not be considered constant. Regarding *N(t)*, we expressed it as the ratio of inverse allele frequency, since it is proportional to *N* [13]:

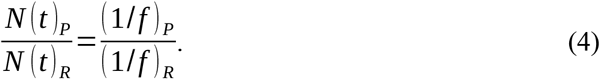

Of note, expression (11) is biased in practice by mutations which are not de novo in the recurrence, but inherited from the primary tumor. Ideally, only de novo mutations should be included to perform this calculation. Finally, the survival rate of tumor cells, *γ*, also could not be considered constant, and we had no indicator or surrogate for this value. For this reason, we have set an arbitrary value for the survival at end of primary tumor, relatively to the start of recurrence, γ_P_/γ_R_ = 1/300.

Using the above, we could obtain expression (5) for *dM/dt* at end of primary tumor:

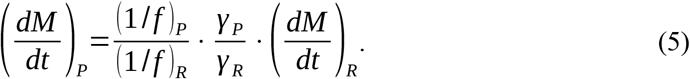

From this, and since the number of mutations at the end of the primary tumor, as well as the rest of parameters, was known, the time in real units at the end of primary tumor could be determined as follows:

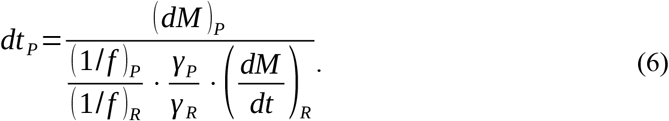

Similarly to the time course in the recurrence, the fact that the last instant of the recurrence, *dt*_*P*_, was known, allowed to calibrate the time course *t* in the primary at each instant, by multiplying by the scaling factor *dt* _*P*_ / *dt*_*a*. *u*., *P*_:

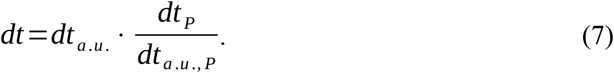

### Tumor cell survival ratio

For time calibration to real units, we have made an assumption on tumor cell survival ratio γ_R_/γ_P_ to determine real time in the primary tumor. To quantify the survival ratio, we have proceeded the other way around, using an assumption on the time course in the primary tumor, in order to determine the ratio γ_R_/γ_P_. We have taken the assumption that the time between the most recent common ancestor (TMRCA) lies either 2 years or 7 years before primary tumor resection.

These durations correspond to the shortest and longest time spans from [15]. The calculation of the tumor cell survival ratio was done by reformulating expression (6) into the following expression:

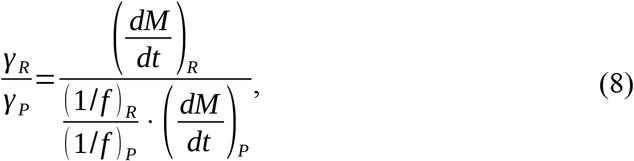

where all terms on the right hand side are known, either from the data (*dM, f, dt*_*R*_) or from assumptions (*dt*_*P*_).

### Tumor expansion profile

From equation (8) in S1 File, time and *1/f* are proportional, with, as modulators, the growth rate *ω*, the tumor cell survival rate *γ*, and the number of tumor cells *N*:

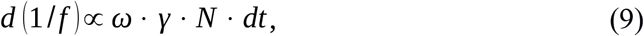

where the increment *d(1/f)* is known from the data, and increment *dt* could be determined above, in section “Time course”. As a consequence, one obtains the product ωγ*N*, by dividing the increment *d(1/f)* by the increment *dt*. The expansion parameters ωγ*N* are known within the constants of expression (8) from S1 File, which are *π* / *K*_*i,r*_. The curves giving *ωγN* as a function of time are denoted expansion curves or expansion profiles.

### Curve segmentation

Segments for curves M(1/f) were determined using R package segmented (v1.1-0), using an objective adjusted R² set to 0.995 (P. virginalis) and 0.9995 (GBM), and using the lowest number of segments which attained this objective, limited to a maximum of 20 breakpoints. P-values of significant changes between segments were evaluated using Davies’ test (implementation from R package segmented). Because the regions before the first breakpoint, and after the last breakpoint, display marked and consistent differences with the general profile of the curve, we hypothesized that the automated segmentation revealed clonal mutations, or mutations which could originate from contamination by normal tissue (mutations with allele frequency lower than the first breakpoint), as well as mutations likely affected by the limit of detection of SNVs on sequencing data (mutations with allele frequency higher than the last breakpoint), We exclude those mutations, and restrict the accepted range to the interval between the first and last breakpoints.

### Classification of expansion profiles

Expansion profiles *ωγN(t)* were visually inspected. This led to the identification of four curve categories (Fig 3). The curves were then manually curated to one of these four categories.

### Statistical analyses

R [40] and Python [41] were used for all statistical analyses. Confidence intervals at 95% probability for the tree root in *P. virginalis* are taken as the 95% highest posterior density (HPD) interval. All statistical tests were unpaired and two-sided, with a level of significance set at 5%. Segmentation p-values correspond to the test of significant difference between segments (Davies’ test, R package segmented). Correlation coefficients between SBS1 and SBS5 were determined using Pearson method, and summarized by their median and IQR over GBM samples. A comparison between groups was made using an unpaired Wilcoxon rank-sum test. Differential time to recurrence between subgroups in the manually sorted *ωγN* curves was assessed using a wilcoxon rank-sum test against curve type 3. Differences on the time to recurrence in the systematic analysis was explored by analysis of variance against the number of maxima in the primary tumors and in the recurrence, with Bonferroni adjustment of *p*-values. The difference of time to recurrence between subgroups based on the number of maxima in the primary tumors was further explored using a non-parametric wilcoxon test and using a F-test of comparison of variances. The association of the *γ*_*R*_*/γ*_*P*_ ratio (n=20) with the pattern of *ωγN* curves was investigated using a F-test of comparison of variances. The association of the *γ*_*R*_*/γ*_*P*_ ratio with the time to recurrence was assessed with a linear regression, using a simple or double log_10_ scale on the *γ*_*R*_*/γ*_*P*_ ratio, with Bonferroni adjustment.

## Acknowledgements

We would like to thank Verena Körber and Thomas Höfer for helpful discussions and for providing data, and Julian Gutekunst for discussions about the methods. We would also like to thank Katharina Hanna for data and for crayfish culture, and Sina Tönges for sample processing. We further acknowledge the German Cancer Research Center Genomics and Proteomics Core Facility for whole-genome sequencing.

## Data availability

Sequence data for marbled crayfish data have been deposited as a National Center for Biotechnology Information BioProject (accession number: PRJNA356499). Glioblastoma data were accessed from the European Genome-phenome Archive (EGA) database, with accession number: EGAS00001003184 (glioblastoma).

## Supporting Information

**S1 Fig. Mutation accumulation, selection and time dynamics of GBM tumors**. Panels A1-D1 show the primary tumor T1, panels A2-D2 show the recurrent tumor T2. (A) Mutation accumulation as a function of the inverse of allele frequency *1/f* (black) and phases from automated segmentation (breakpoints (grey) and segments (blue)). The confidence band at 95% level is indicated in grey. (B) Nonsynonymous to synonymous ratio. Purple and blue stars show nonsynonymous and synonymous mutations, respectively. The smoothened ratio is shown in red. (C) Clock-like and non-clock-like mutational signatures. (D) Mutation accumulation as a function of time.

**S2 Fig. Transition of primary tumor to recurrent tumor**. (A) Dynamics of growth rate *ω* times tumor cell survival rate *γ* times number of cells *N*, for (P) the primary tumor and (R) the recurrence. (B) Time-resolved mutation accumulation for primary tumor and recurrence.

**S1 File. Supplementary Methods**.

**S2 File. Supplementary Tables**.

