## Supplemental Figure 1 for "Time-resolved, integrated analysis of clonally evolving genomes"

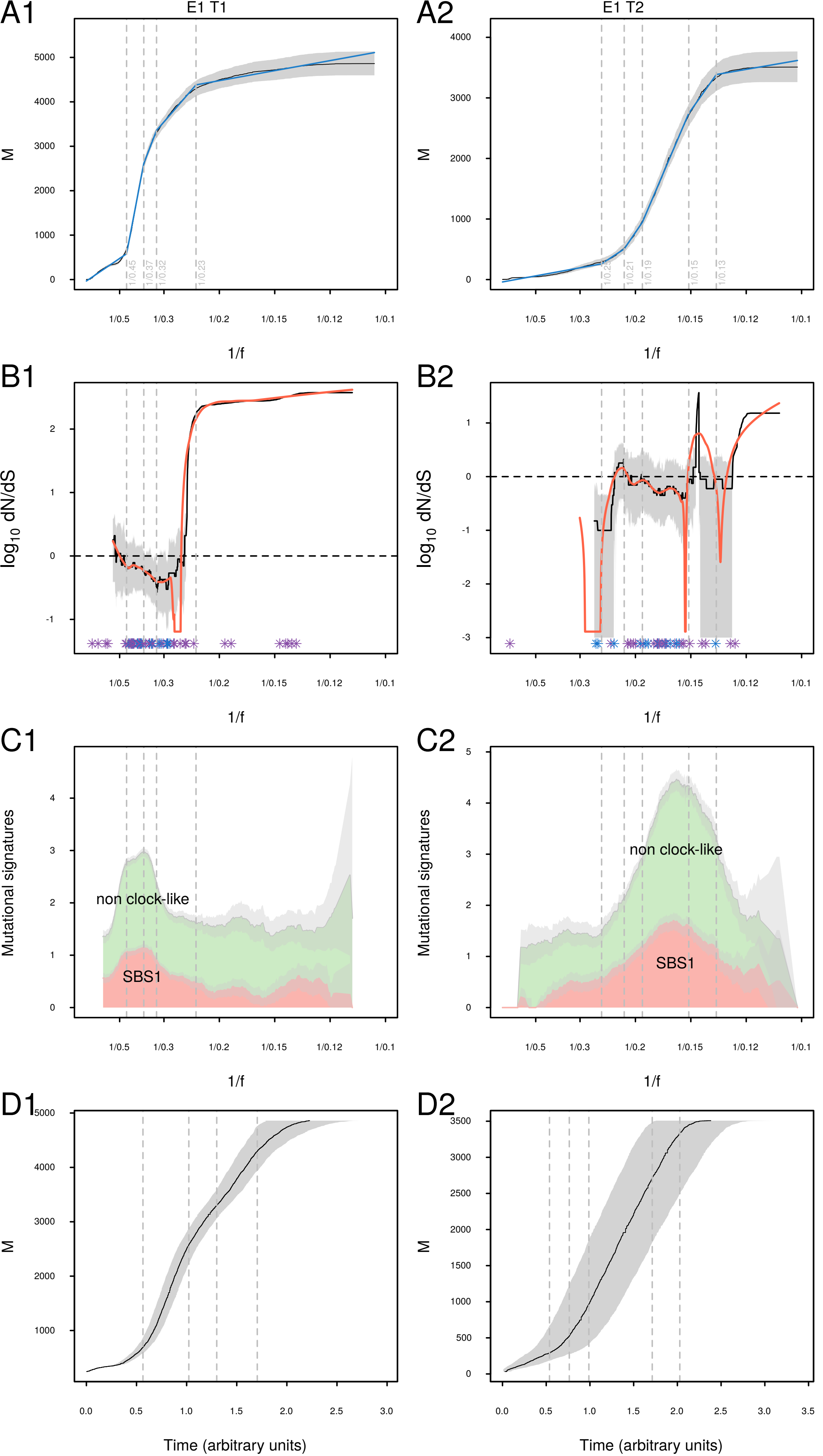
A1
E1 T1
A2
E1 T2
M
1/f
B1
B2
log10 dN/dS
C1
C2
Mutational signatures
non clock-like
SBS1
D1
D2
Time (arbitrary units)

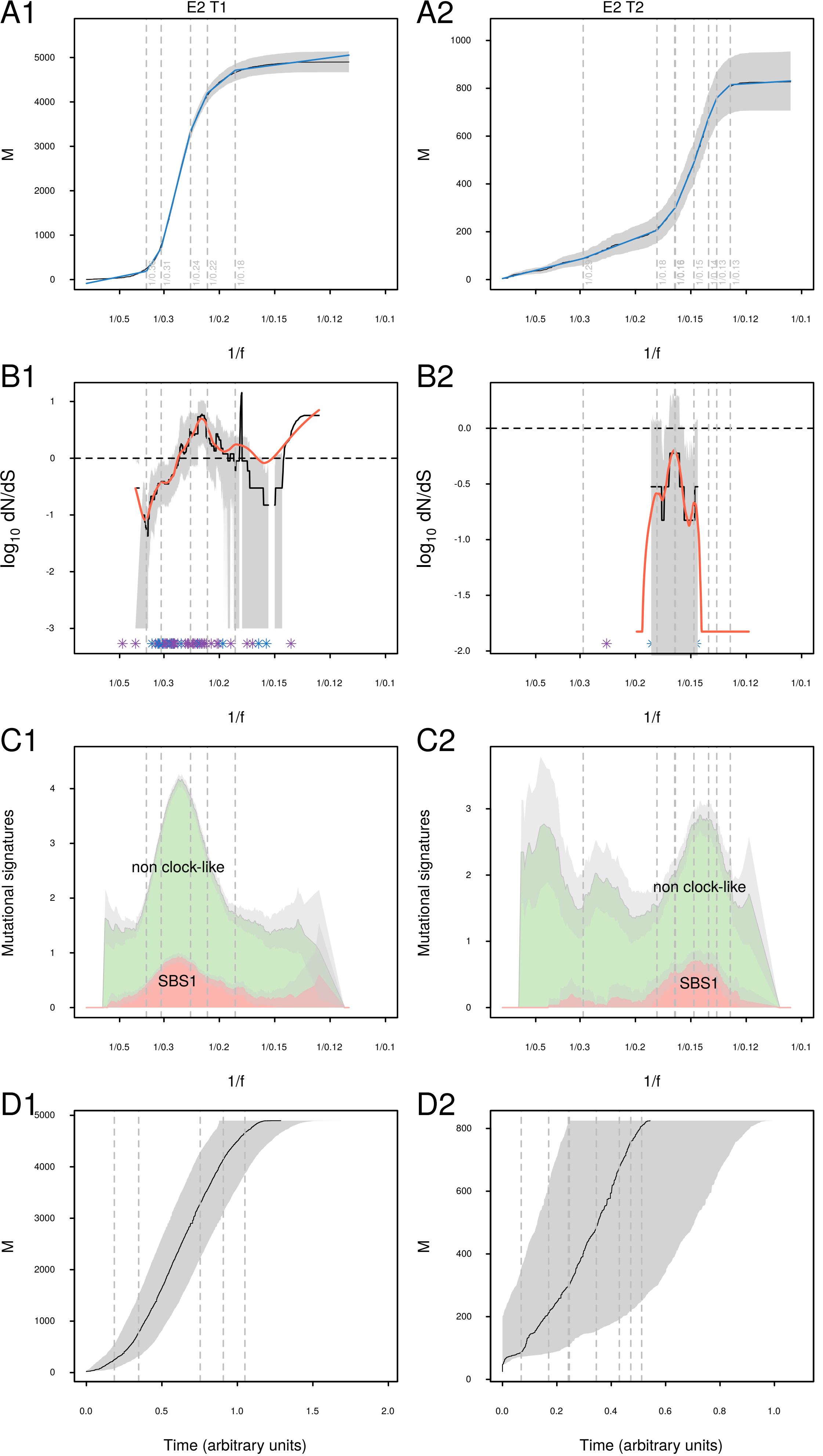

A1
E2 T1
A2
E2 T2
M
1/f
B1
B2
log10 dN/dS
C1
C2
Mutational signatures
non clock-like
SBS1
D1
D2
Time (arbitrary units)

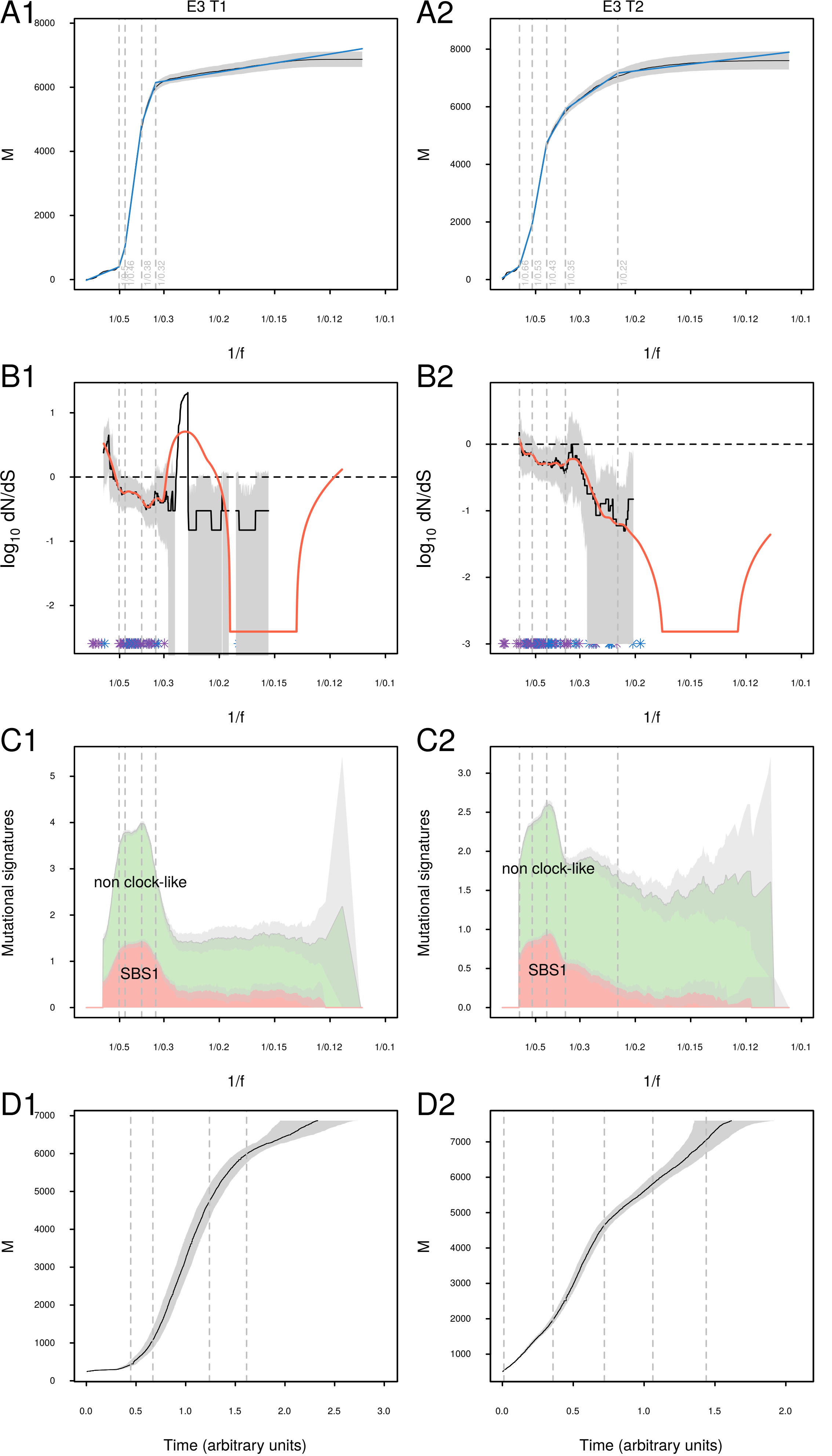

A1
E3 T1
A2
E3 T2
M
1/f
1/0.5
1/0.46
1/0.38
1/0.32
1/0.66
1/0.53
1/0.43
1/0.35
1/0.22
B1
B2
log10 dN/dS
C1
C2
Mutational signatures
non clock-like
SBS1
D1
D2
Time (arbitrary units)

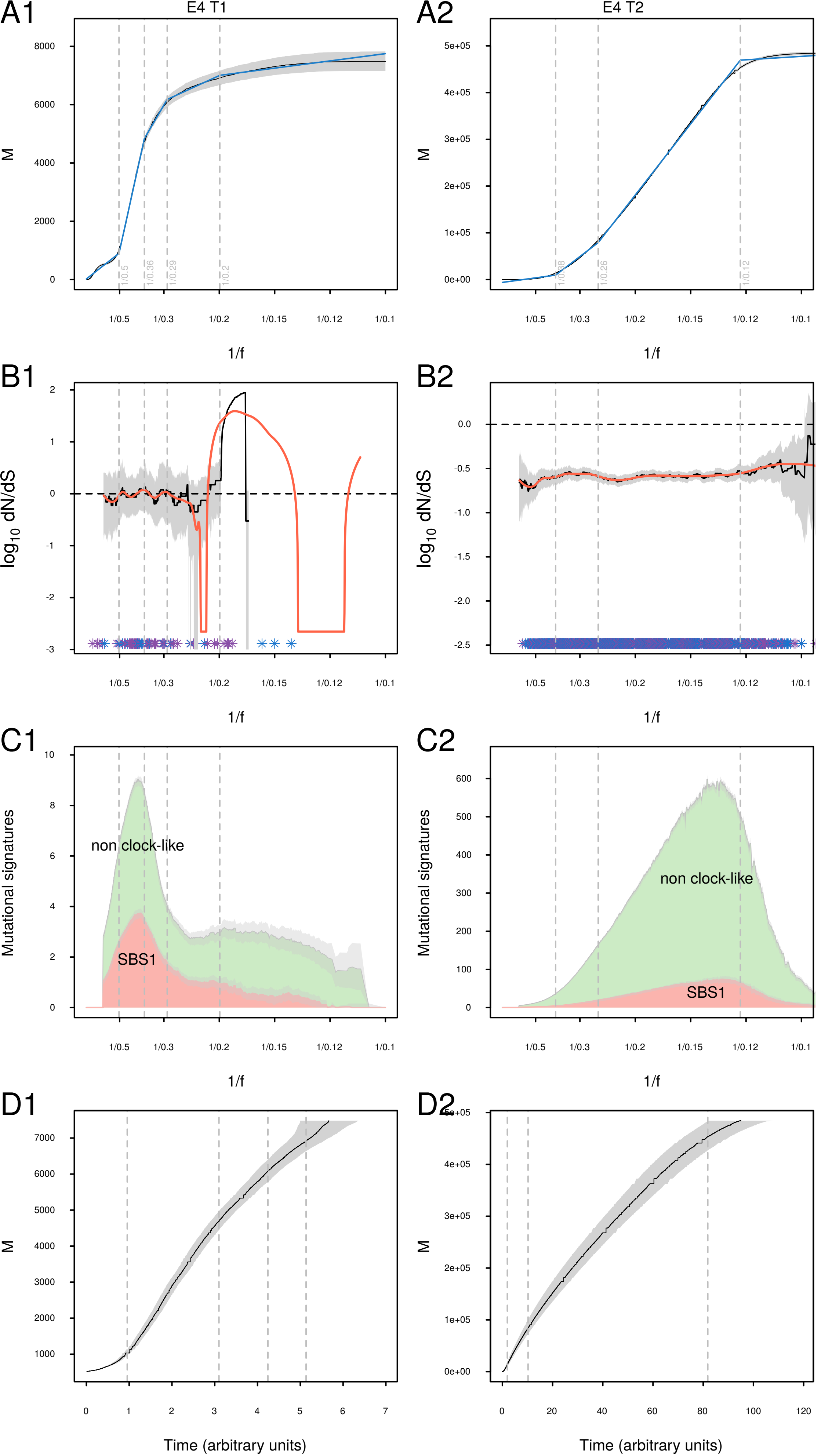

A1
E4 T1
A2
E4 T2
M
1/f
B1
B2
log10 dN/dS
C1
C2
Mutational signatures
non clock-like
SBS1
D1
D2
Time (arbitrary units)

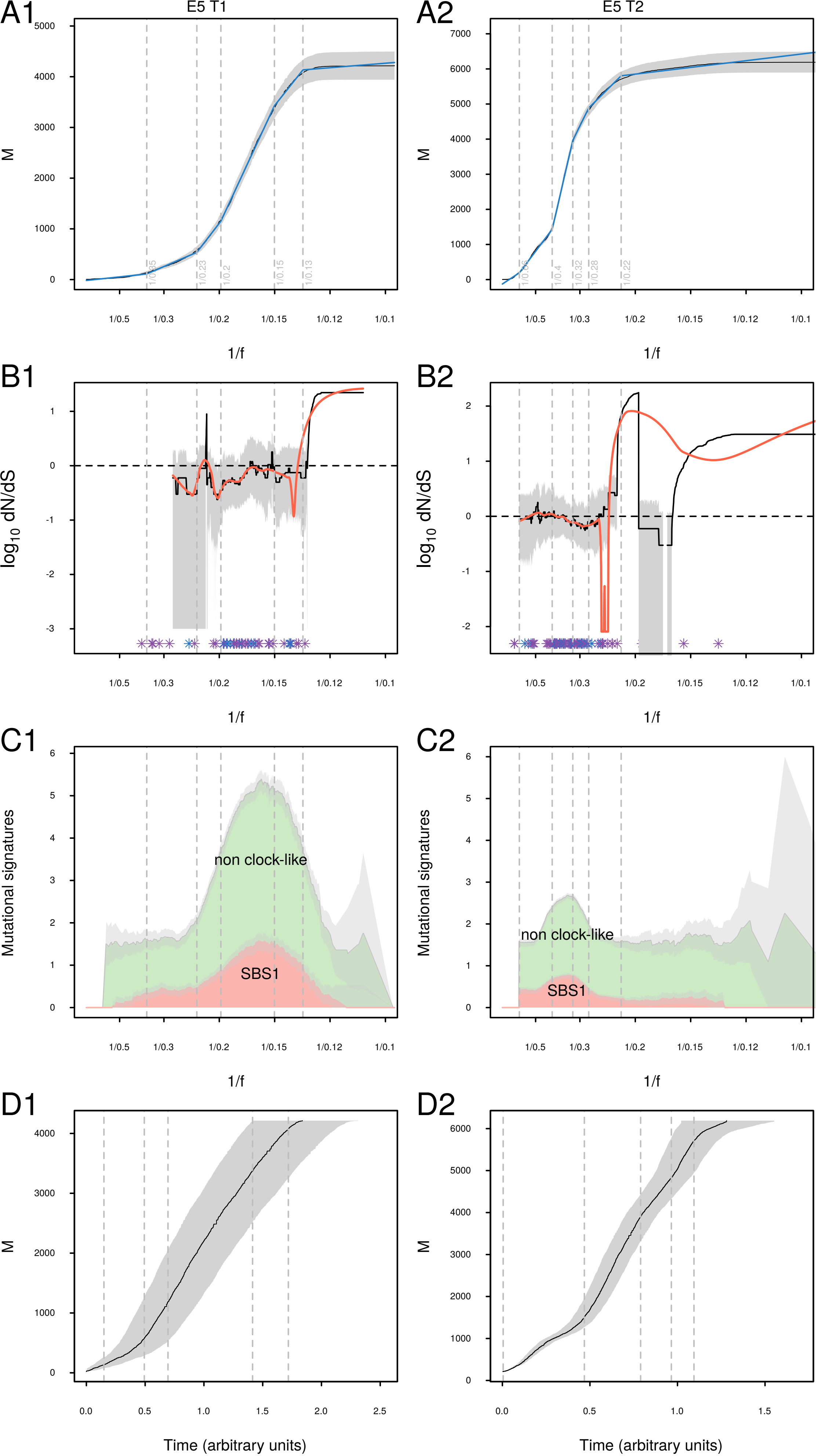

A1
E5 T1
M
1/f
A2
E5 T2
B1
log10 dN/dS
B2
C1
Mutational signatures
non clock-like
SBS1
C2
D1
Time (arbitrary units)
D2

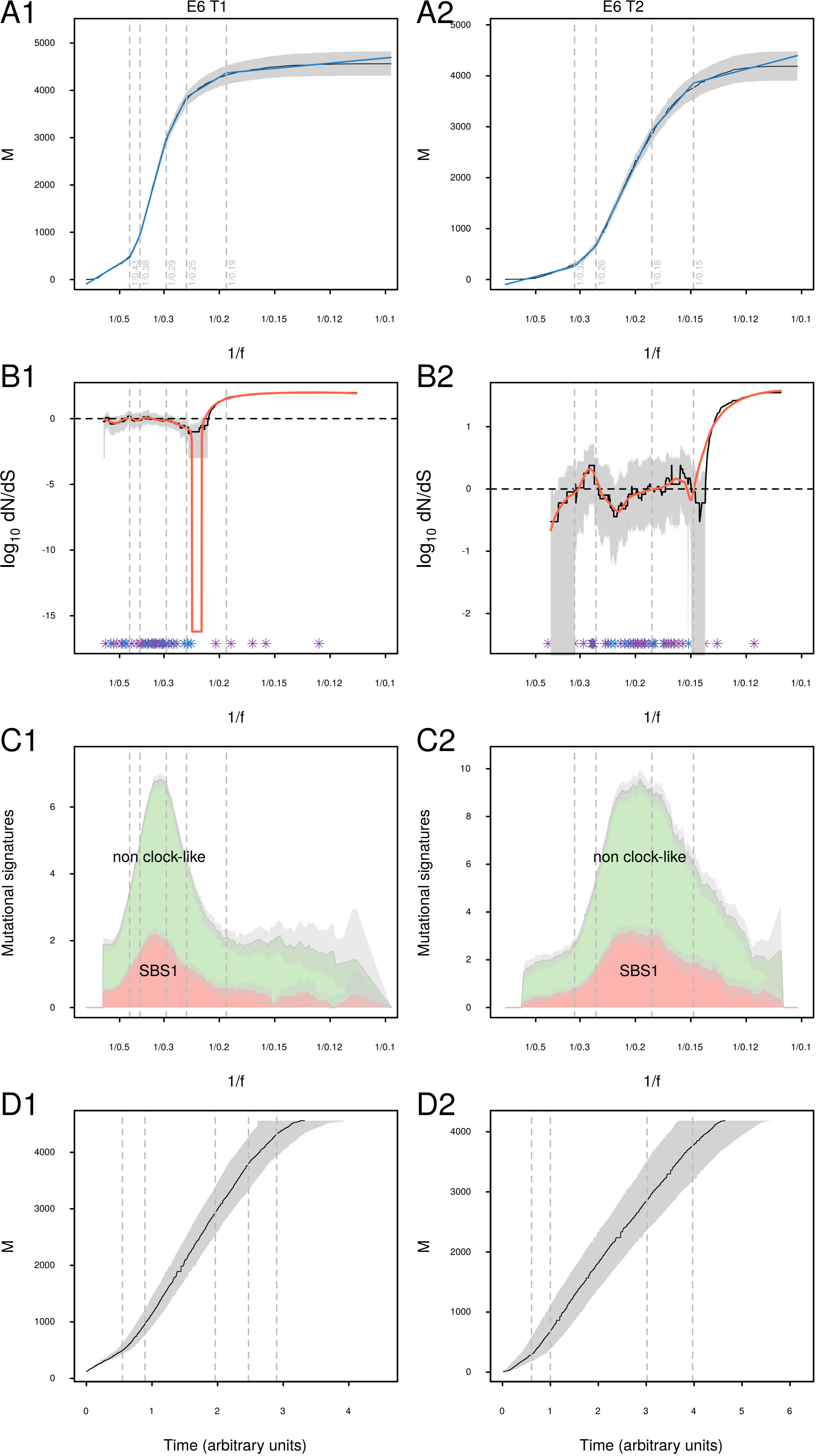
A1
E6 T1
A2
E6 T2
M
1/0.43
1/0.38
1/0.29
1/0.25
1/0.19
1/0.26
1/0.18
1/0.15
1/f
B1
B2
log10 dN/dS
C1
C2
Mutational signatures
non clock-like
SBS1
D1
D2
Time (arbitrary units)

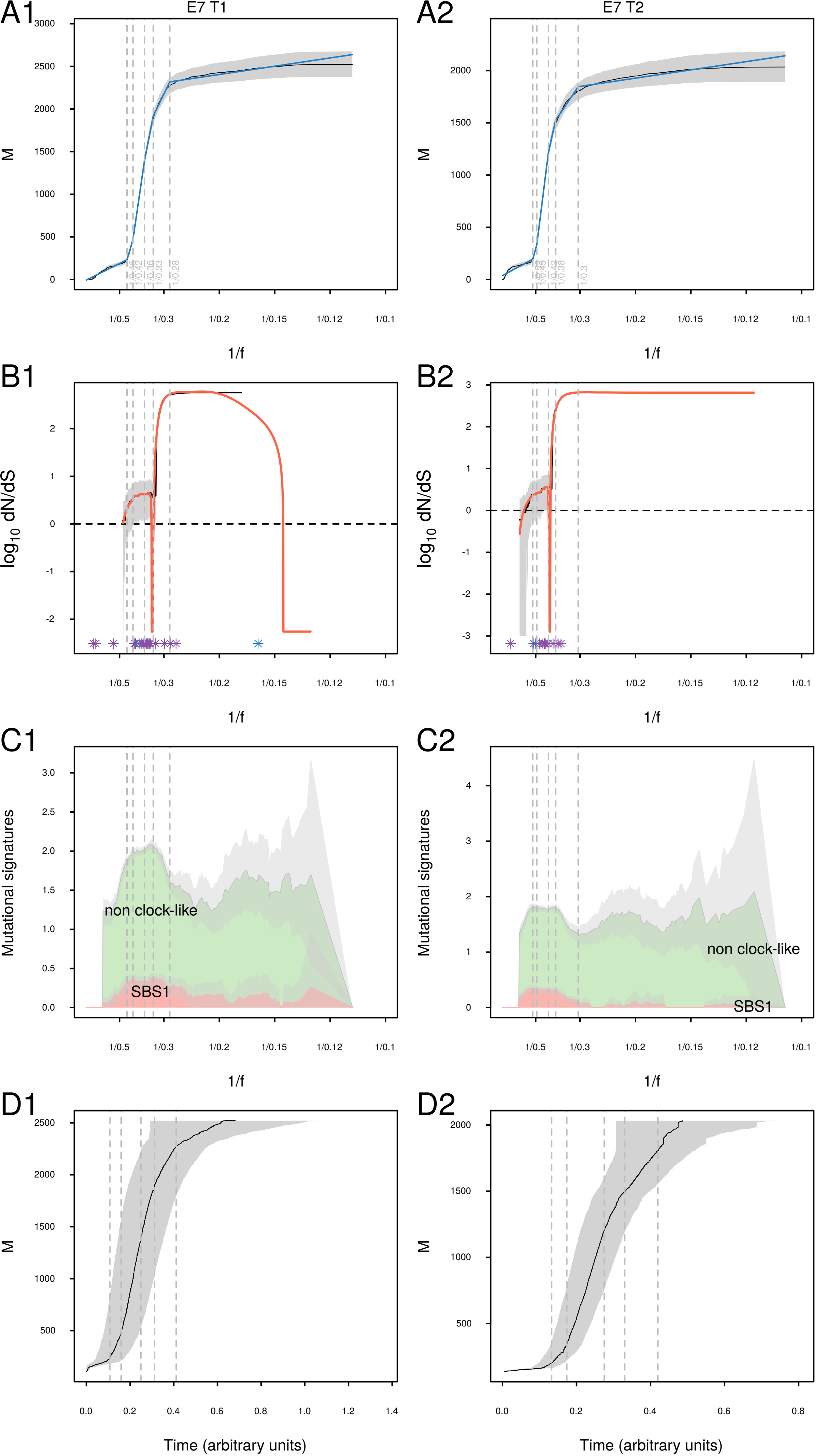

A1
E7 T1
A2
E7 T2
M
1/f
B1
B2
log10 dN/dS
C1
C2
Mutational signatures
non clock-like
SBS1
D1
D2
Time (arbitrary units)

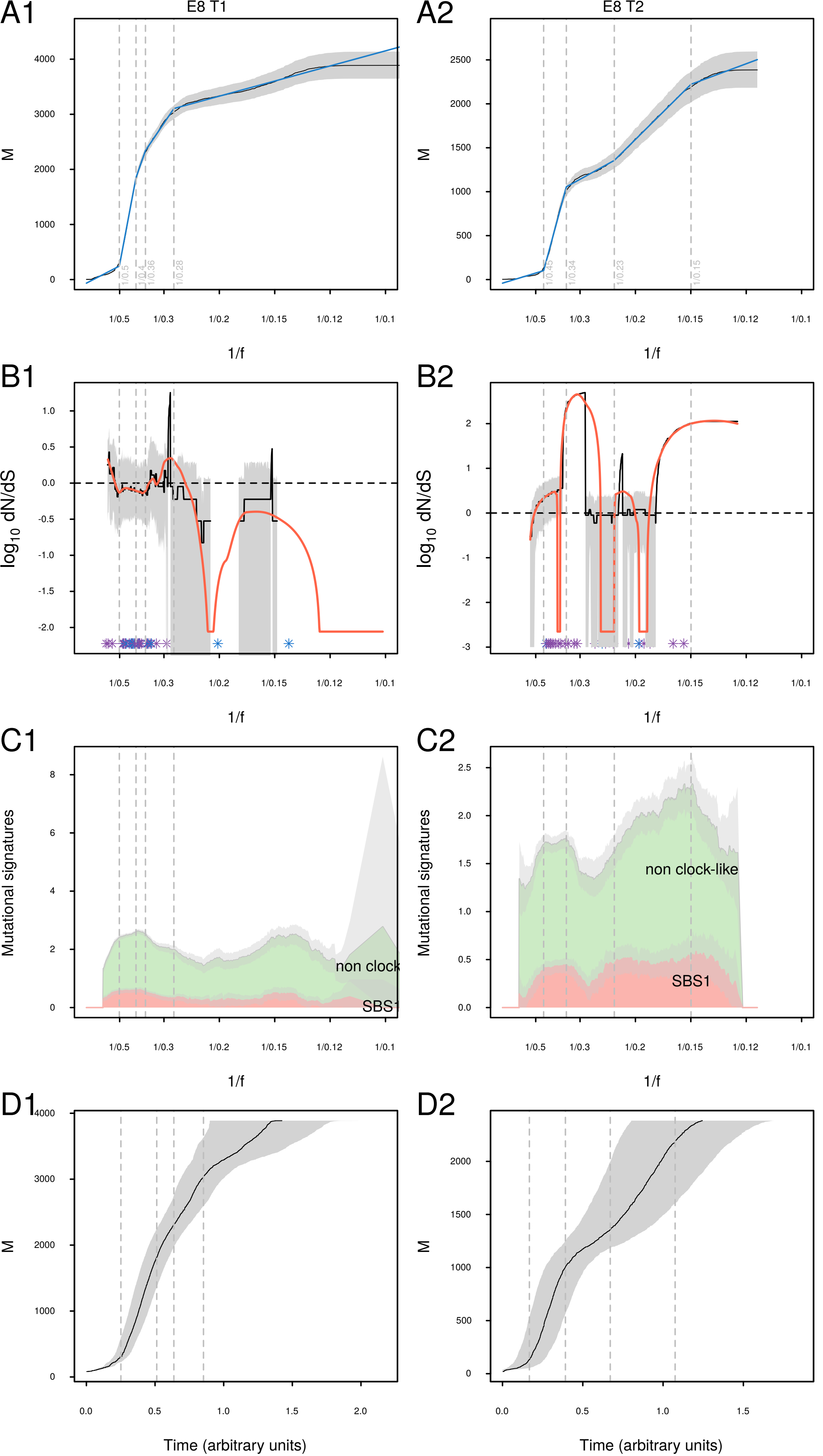
A1
E8 T1
A2
E8 T2
M
1/0.5
1/0.4
1/0.36
1/0.28
1/0.45
1/0.34
1/0.23
1/0.15
1/0.5
1/0.3
1/0.2
1/0.15
1/0.12
1/0.1
1/f
B1
B2
log10 dN/dS
C1
C2
Mutational signatures
non clock
non clock-like
SBS1
D1
D2
Time (arbitrary units)

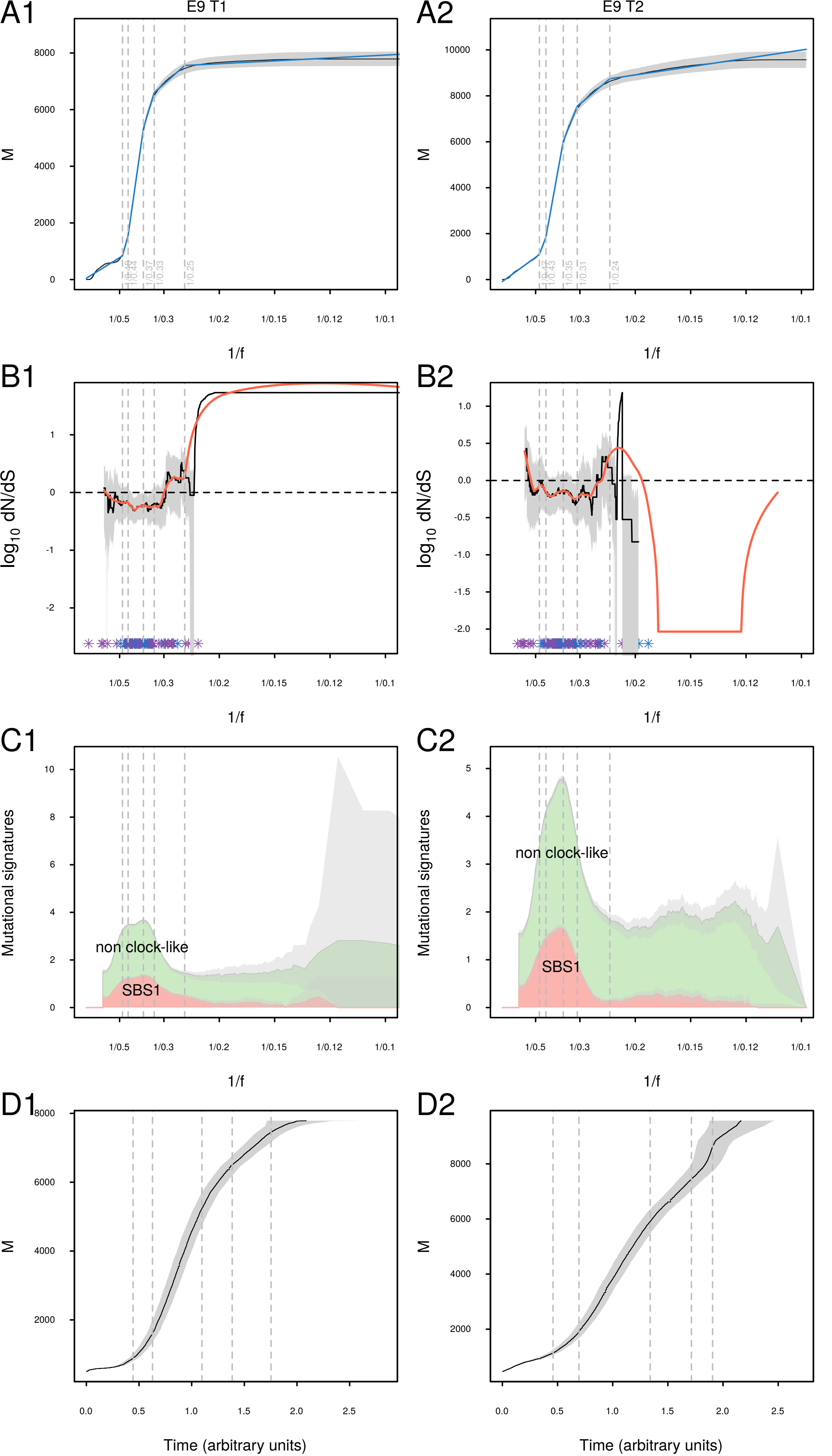

A1
E9 T1
A2
E9 T2
M
1/f
B1
B2
log10 dN/dS
C1
C2
Mutational signatures
non clock-like
SBS1
D1
D2
Time (arbitrary units)

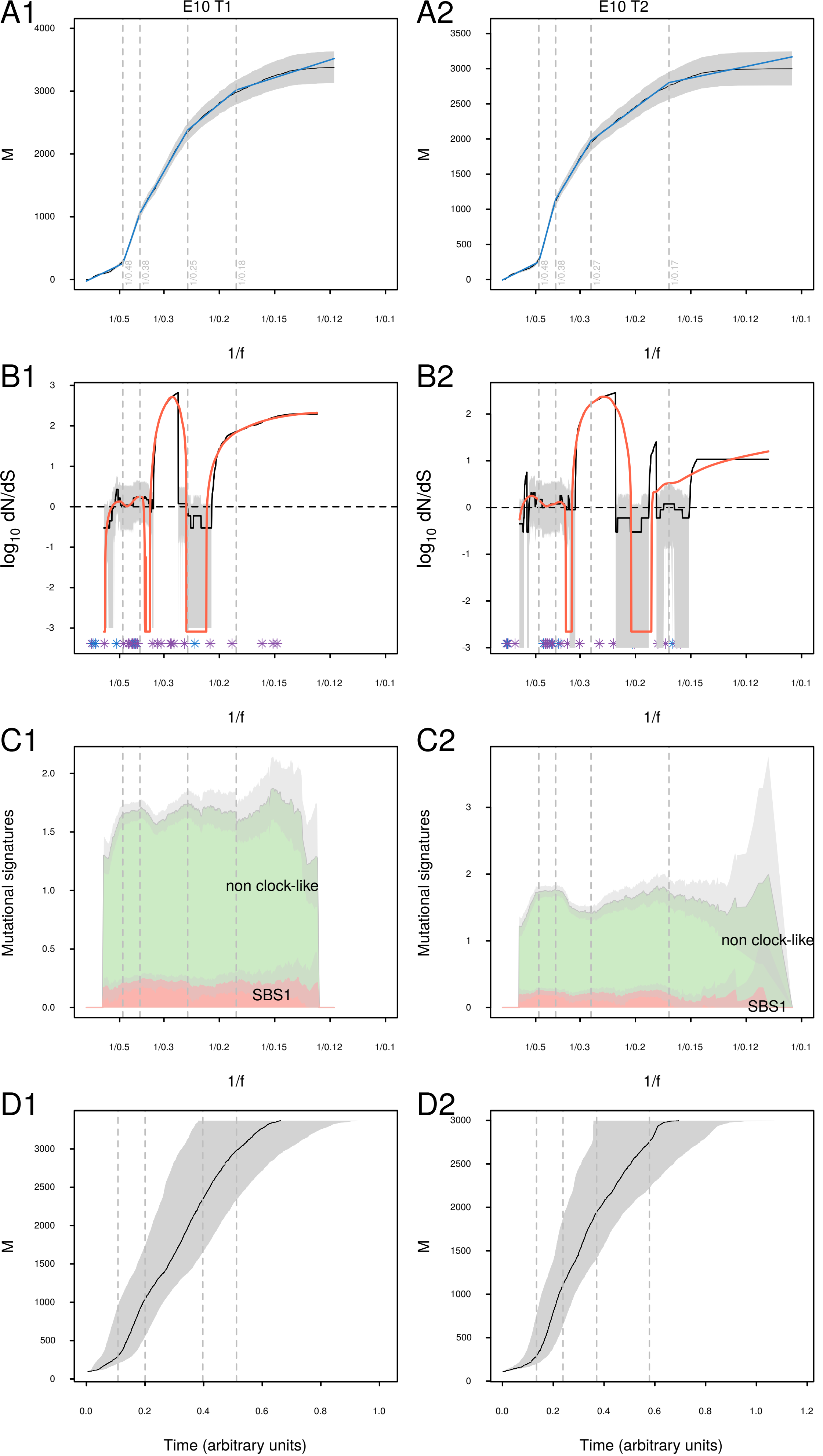

A1
E10 T1
A2
E10 T2
M
1/f
B1
B2
log10 dN/dS
C1
C2
Mutational signatures
non clock-like
SBS1
D1
D2
Time (arbitrary units)

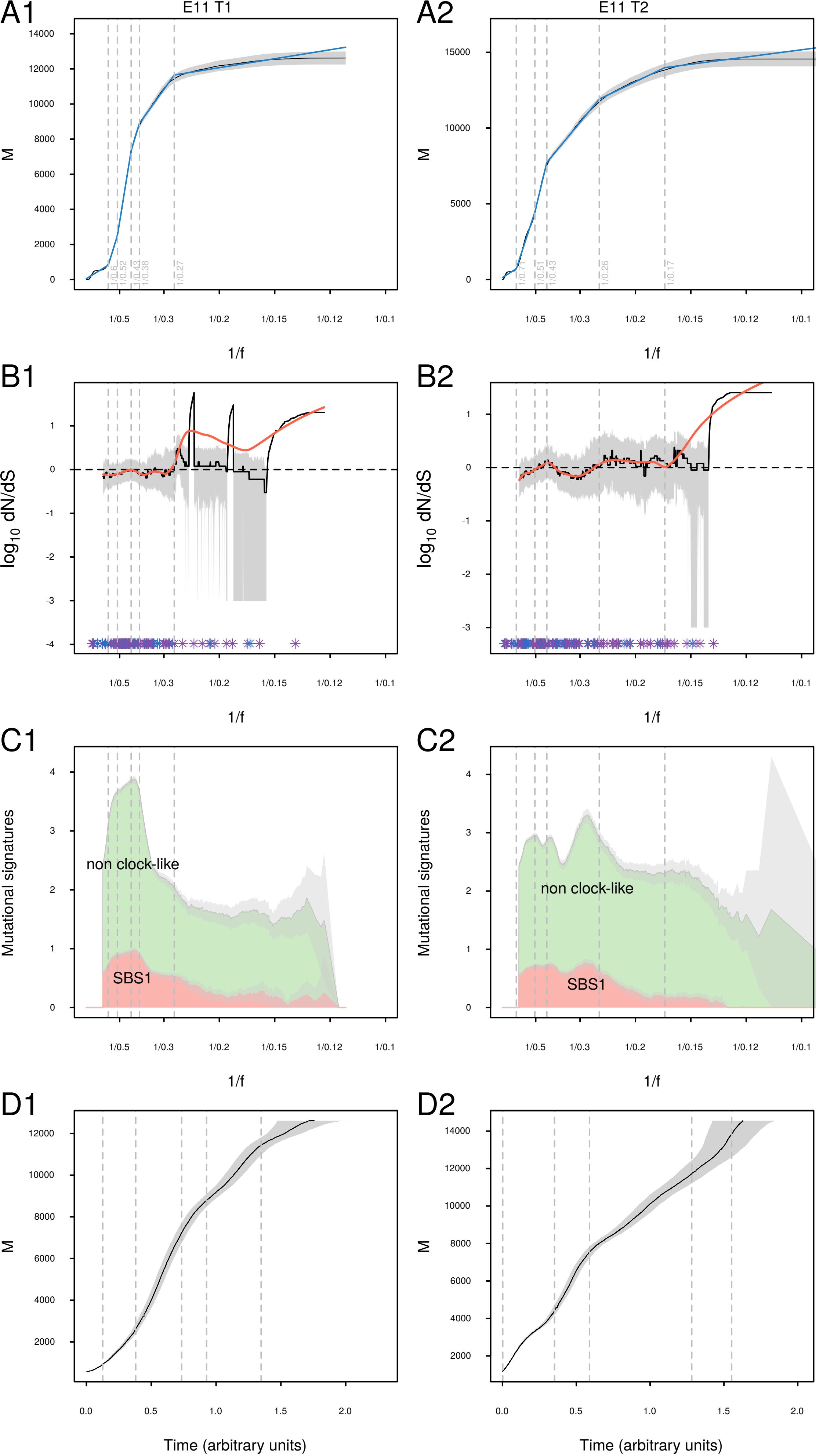
A1
E11 T1
A2
E11 T2
M
1/f
B1
B2
log10 dN/dS
C1
C2
Mutational signatures
non clock-like
SBS1
D1
D2
Time (arbitrary units)

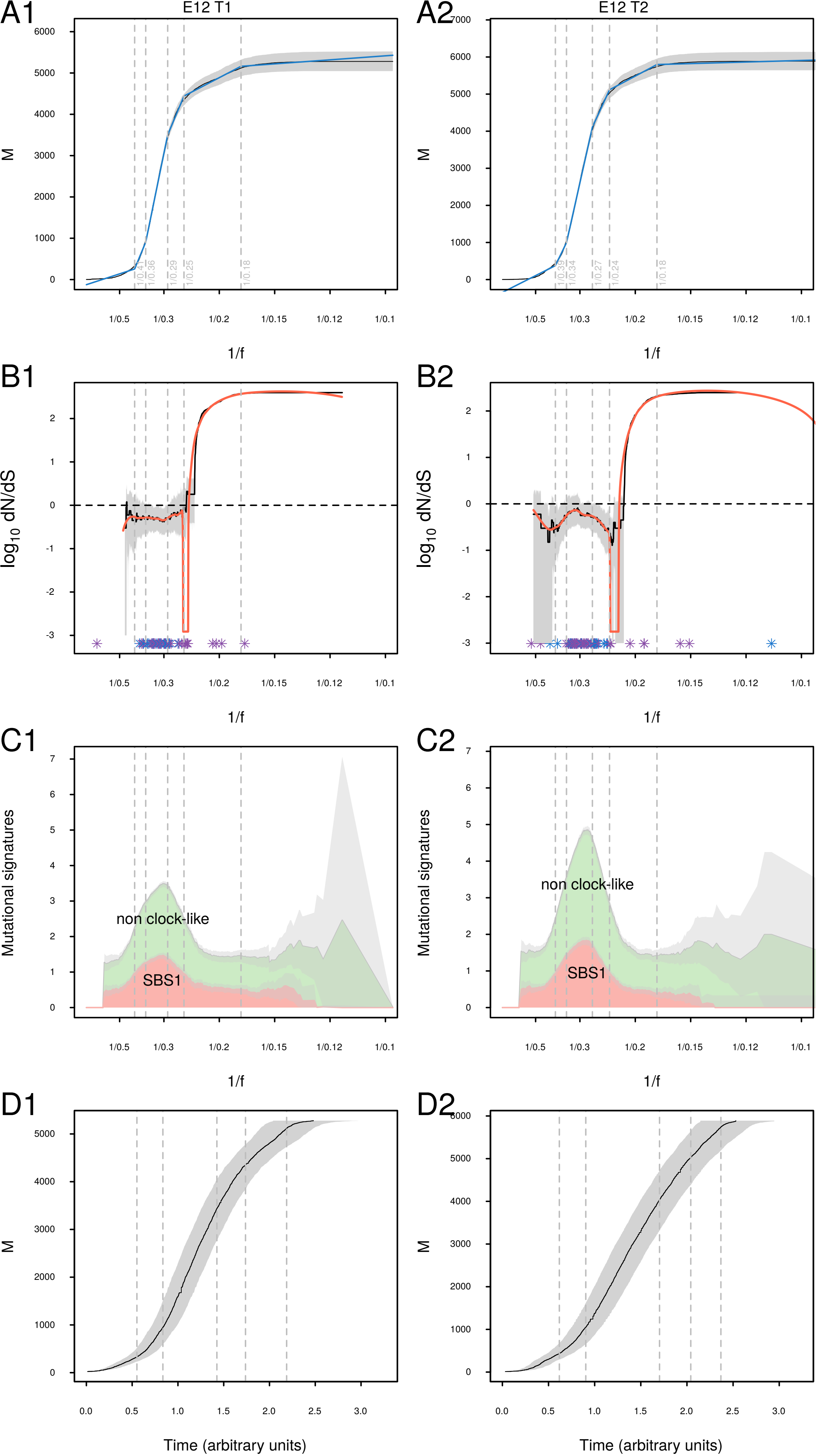
A1
E12 T1
A2
E12 T2
M
1/f
1/0.41
1/0.36
1/0.29
1/0.25
1/0.18
1/0.39
1/0.34
1/0.27
1/0.24
B1
B2
log10 dN/dS
C1
C2
Mutational signatures
non clock-like
SBS1
D1
D2
Time (arbitrary units)

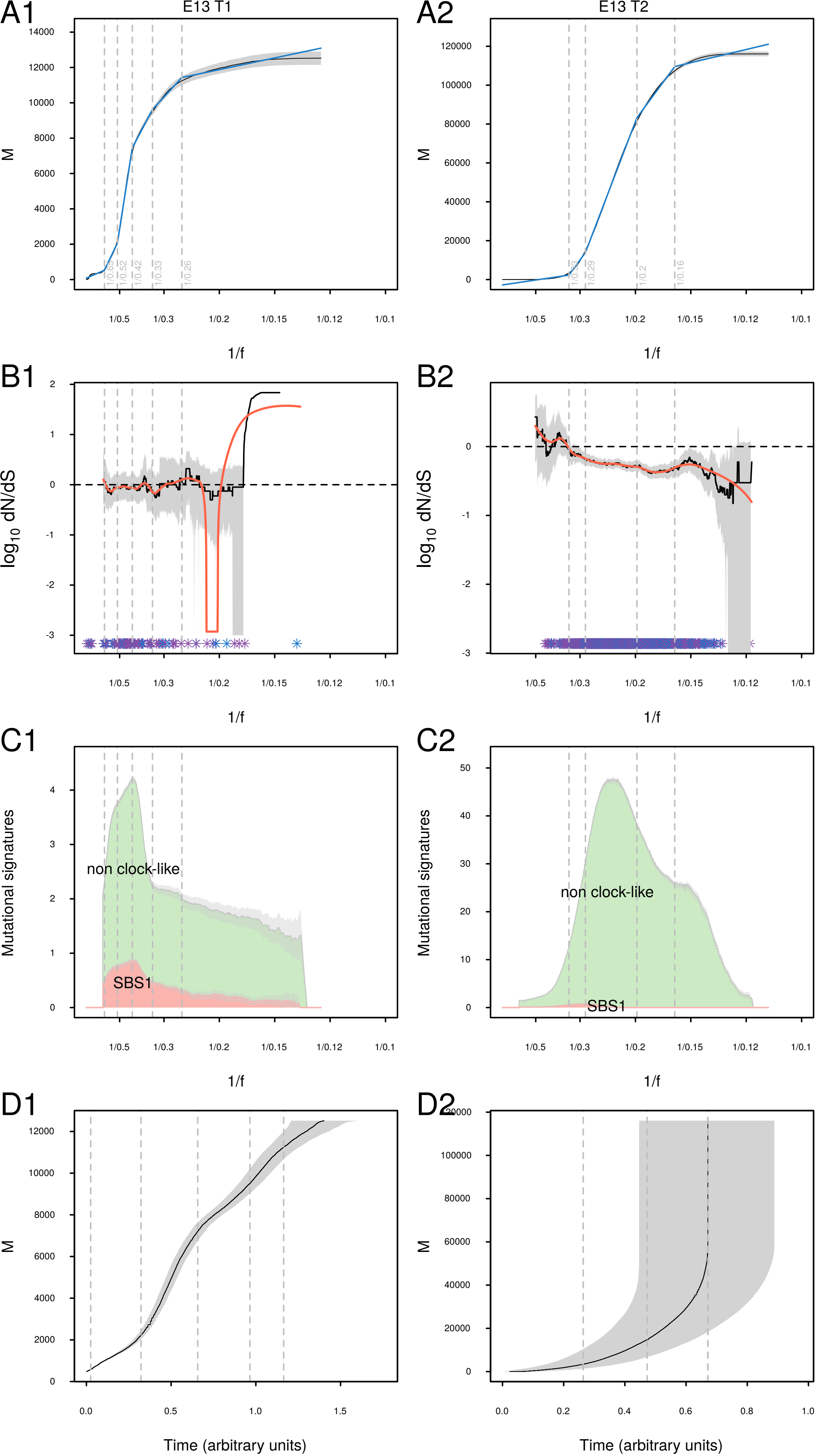

A1
E13 T1
A2
E13 T2
M
1/f
1/0.5
1/0.3
1/0.2
1/0.15
1/0.12
1/0.1
B1
B2
log10 dN/dS
C1
C2
Mutational signatures
non clock-like
SBS1
D1
D2
Time (arbitrary units)

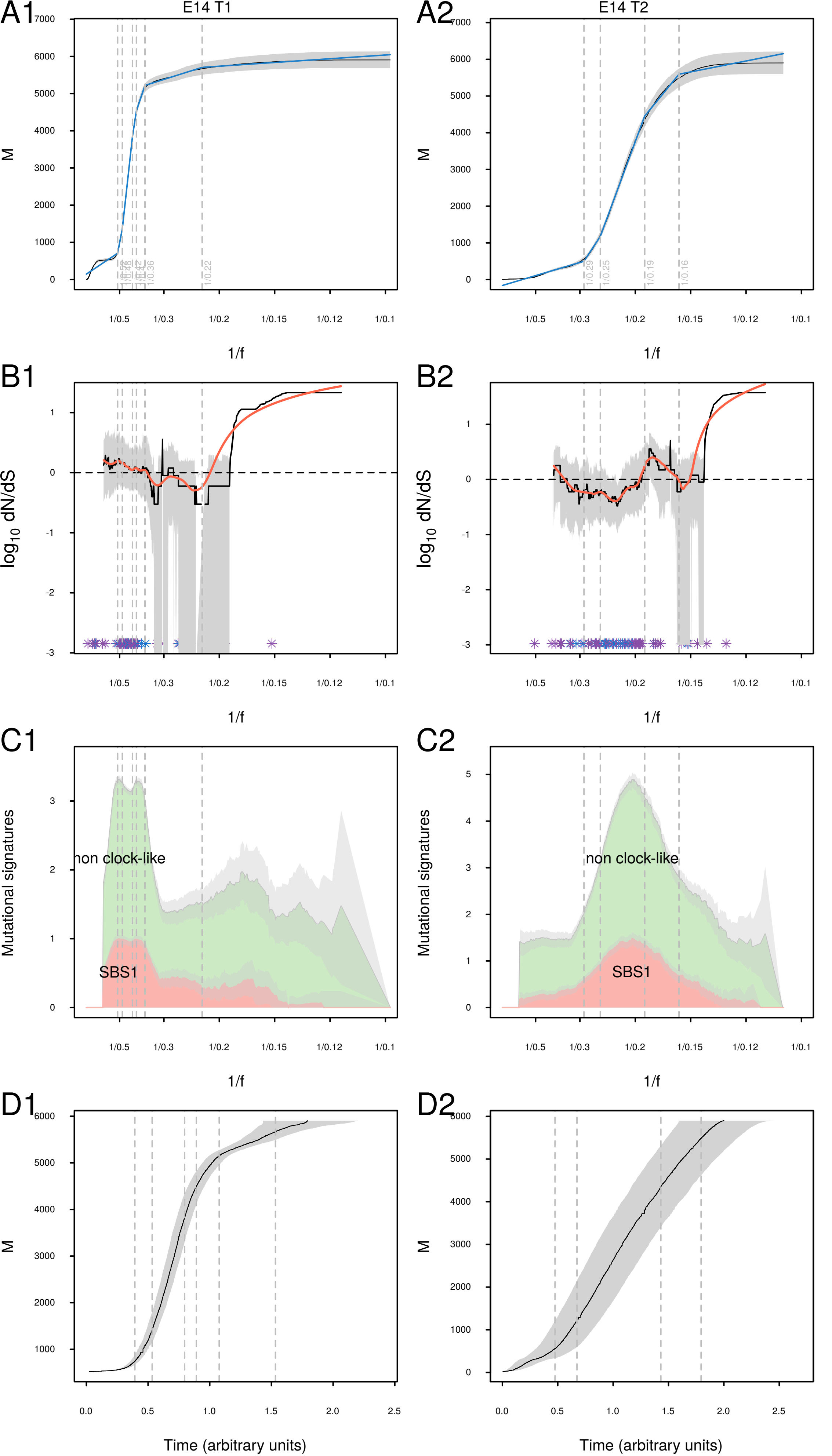
A1
E14 T1
A2
E14 T2
M
1/f
B1
B2
log10 dN/dS
C1
C2
Mutational signatures
non clock-like
SBS1
D1
D2
Time (arbitrary units)

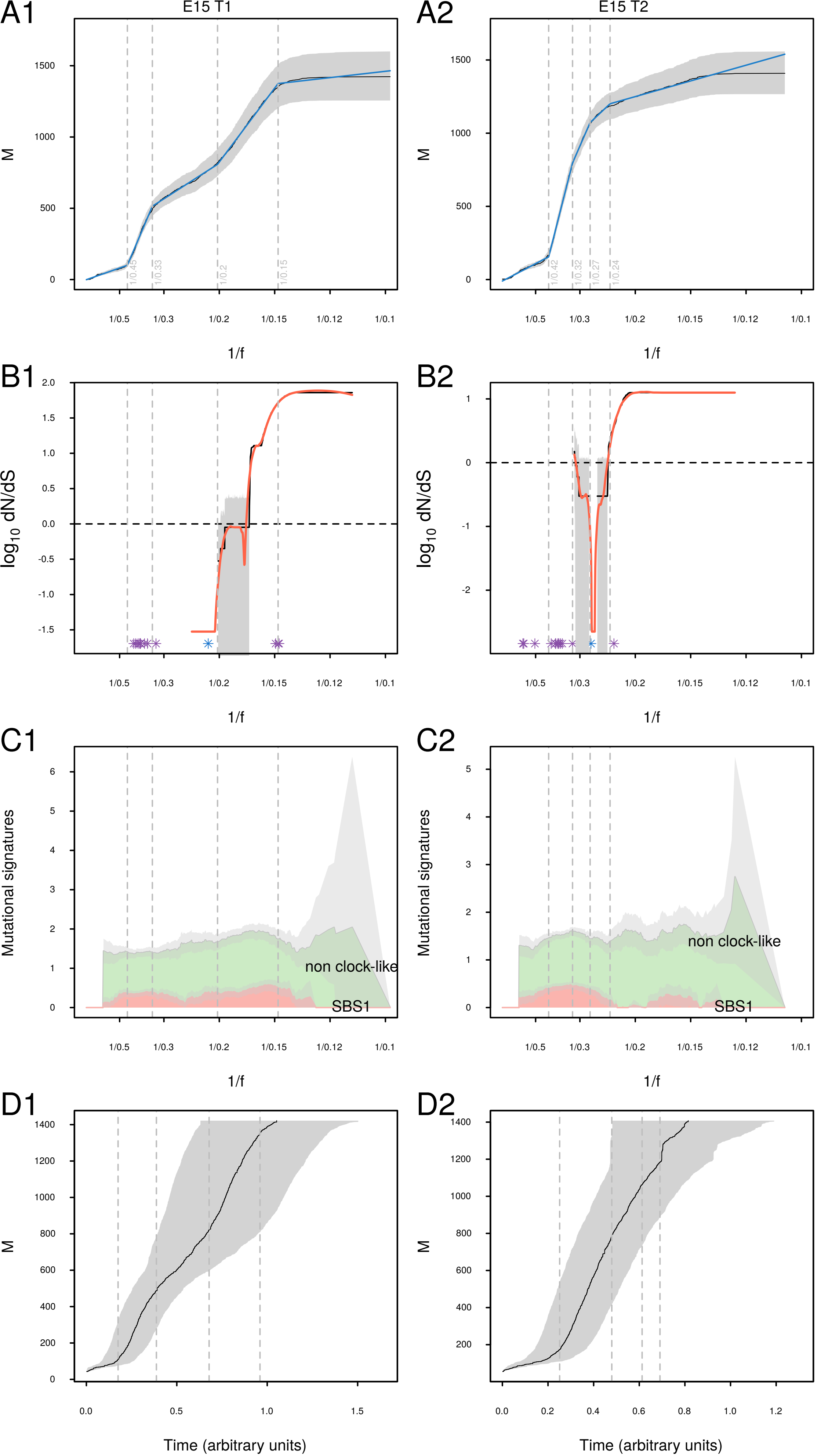
A1
E15 T1
A2
E15 T2
M
1/f
B1
B2
log10 dN/dS
C1
C2
Mutational signatures
non clock-like
SBS1
D1
D2
Time (arbitrary units)

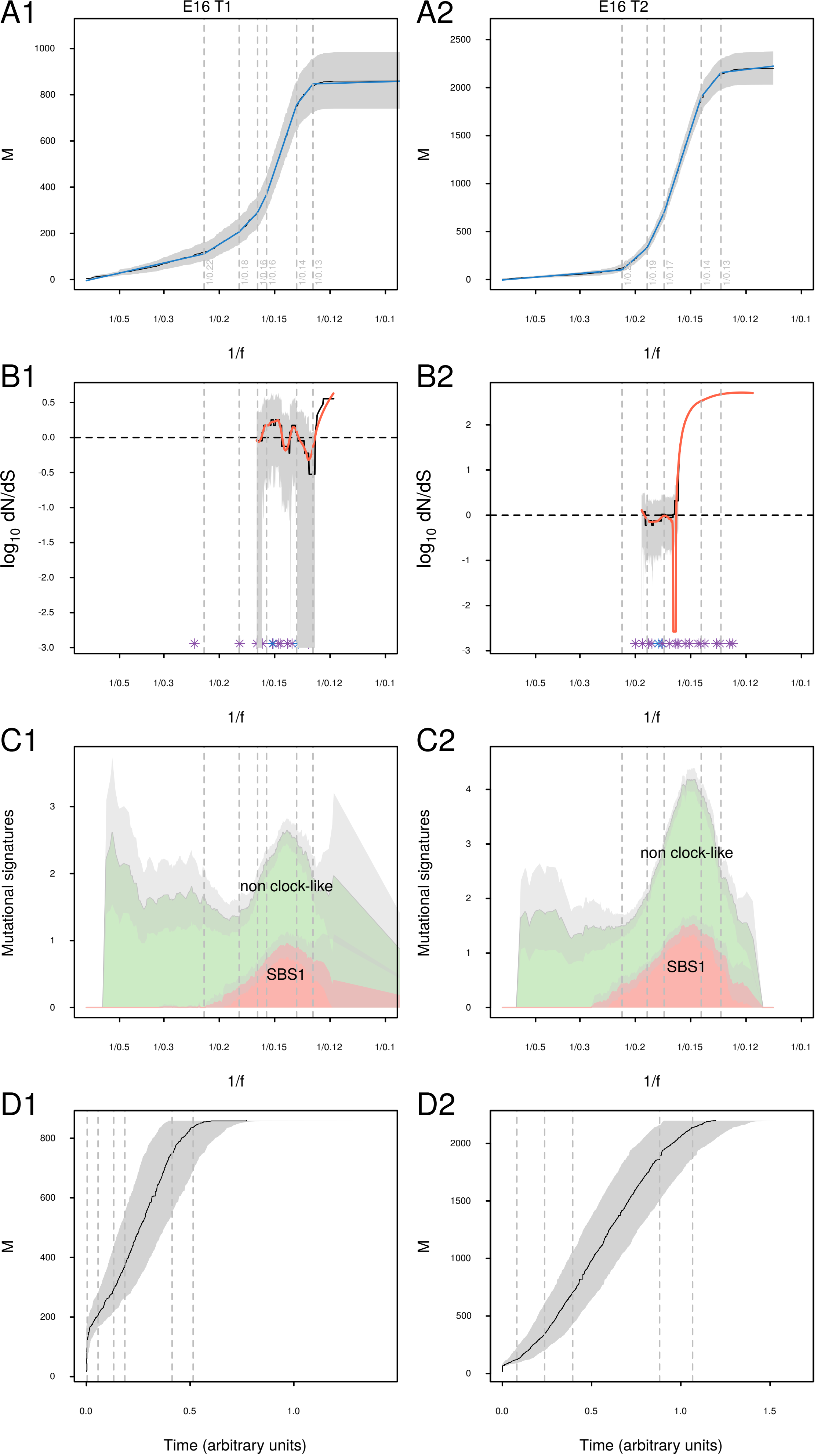

A1
E16 T1
A2
E16 T2
M
1/f
B1
B2
log10 dN/dS
C1
C2
Mutational signatures
non clock-like
SBS1
D1
D2
Time (arbitrary units)

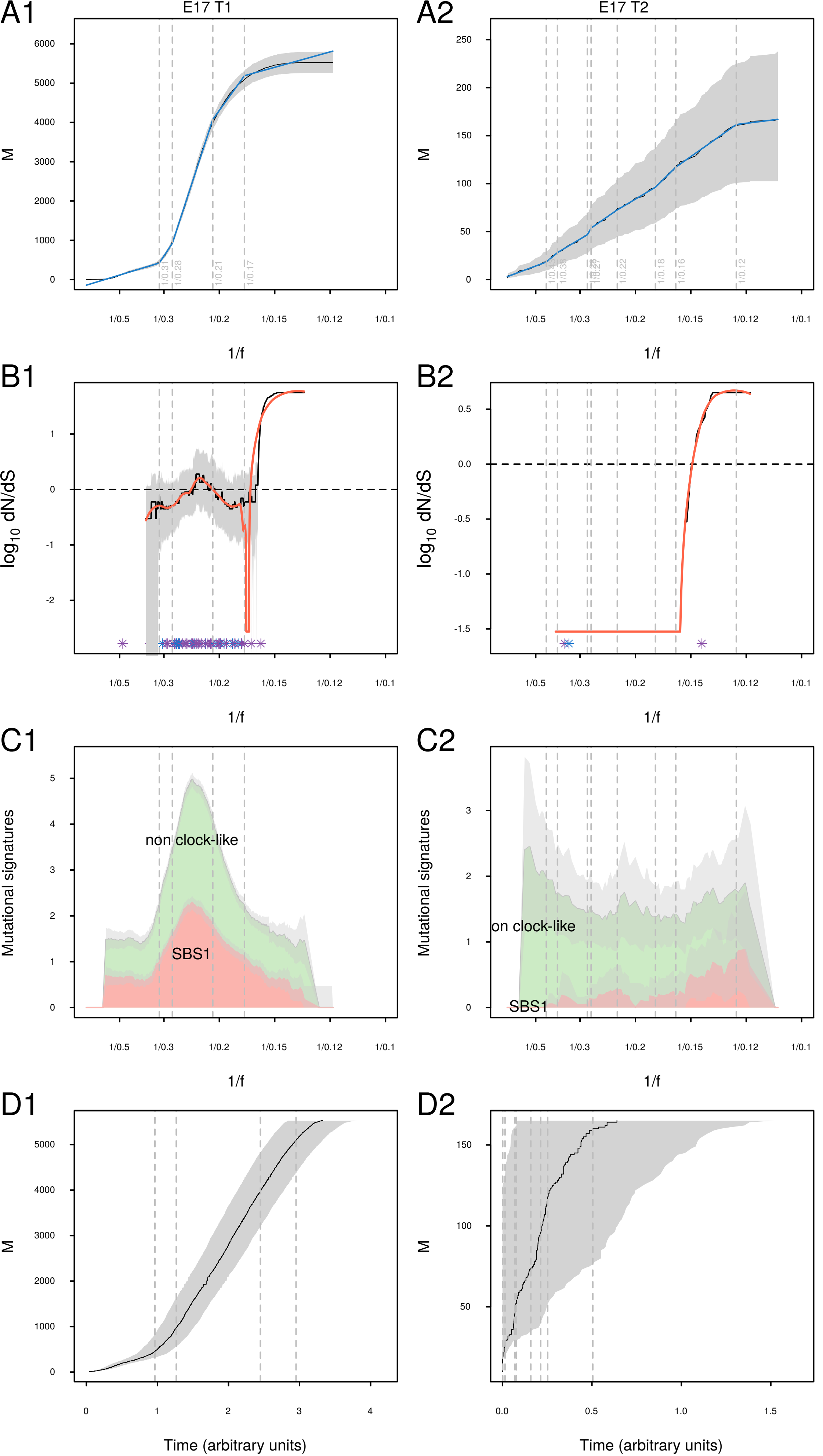

A1
E17 T1
A2
E17 T2
M
1/f
B1
B2
log10 dN/dS
C1
C2
Mutational signatures
non clock-like
SBS1
D1
D2
Time (arbitrary units)

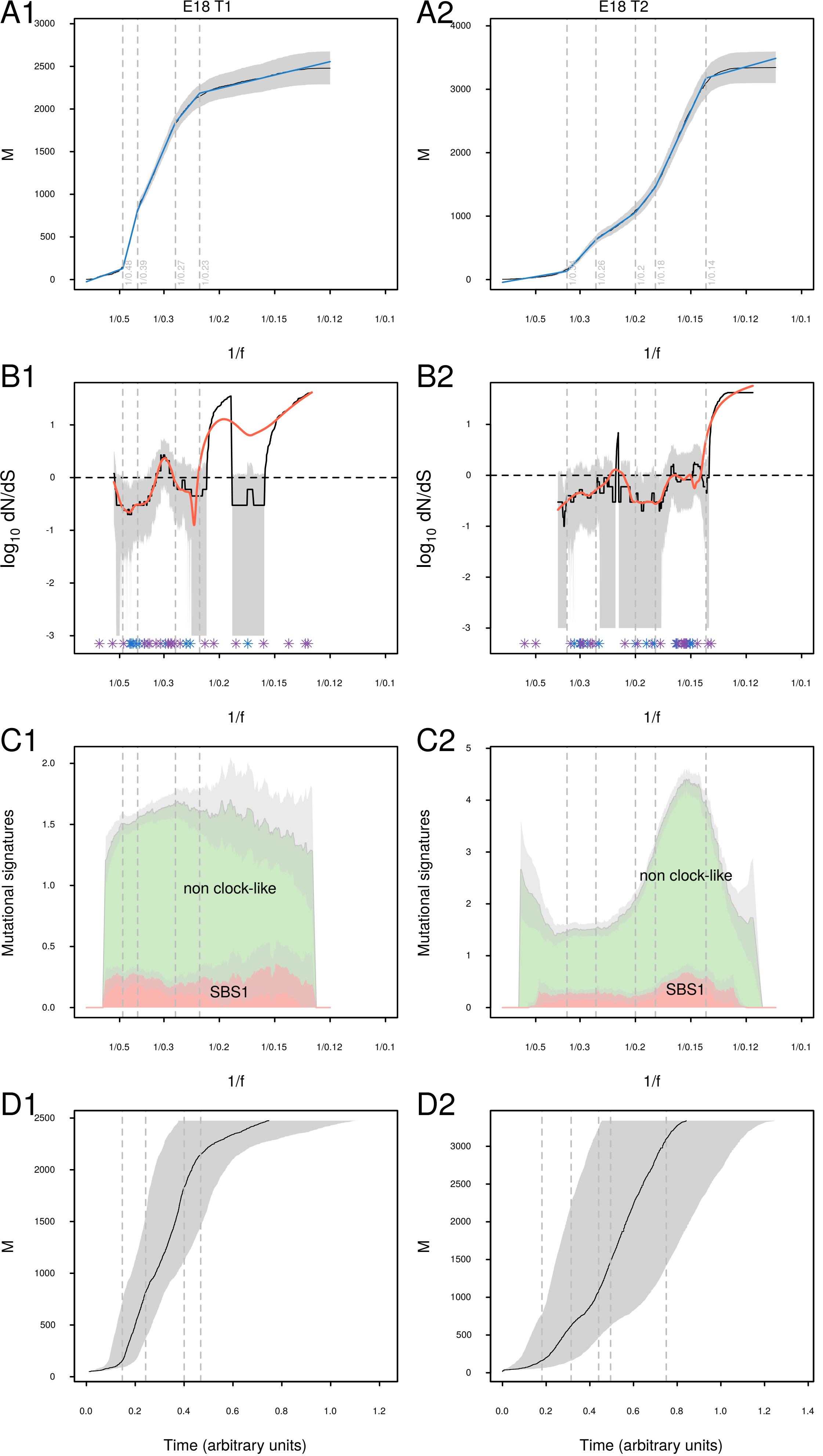

A1
E18 T1
A2
E18 T2
M
1/f
B1
B2
log10 dN/dS
C1
C2
Mutational signatures
non clock-like
SBS1
D1
D2
Time (arbitrary units)

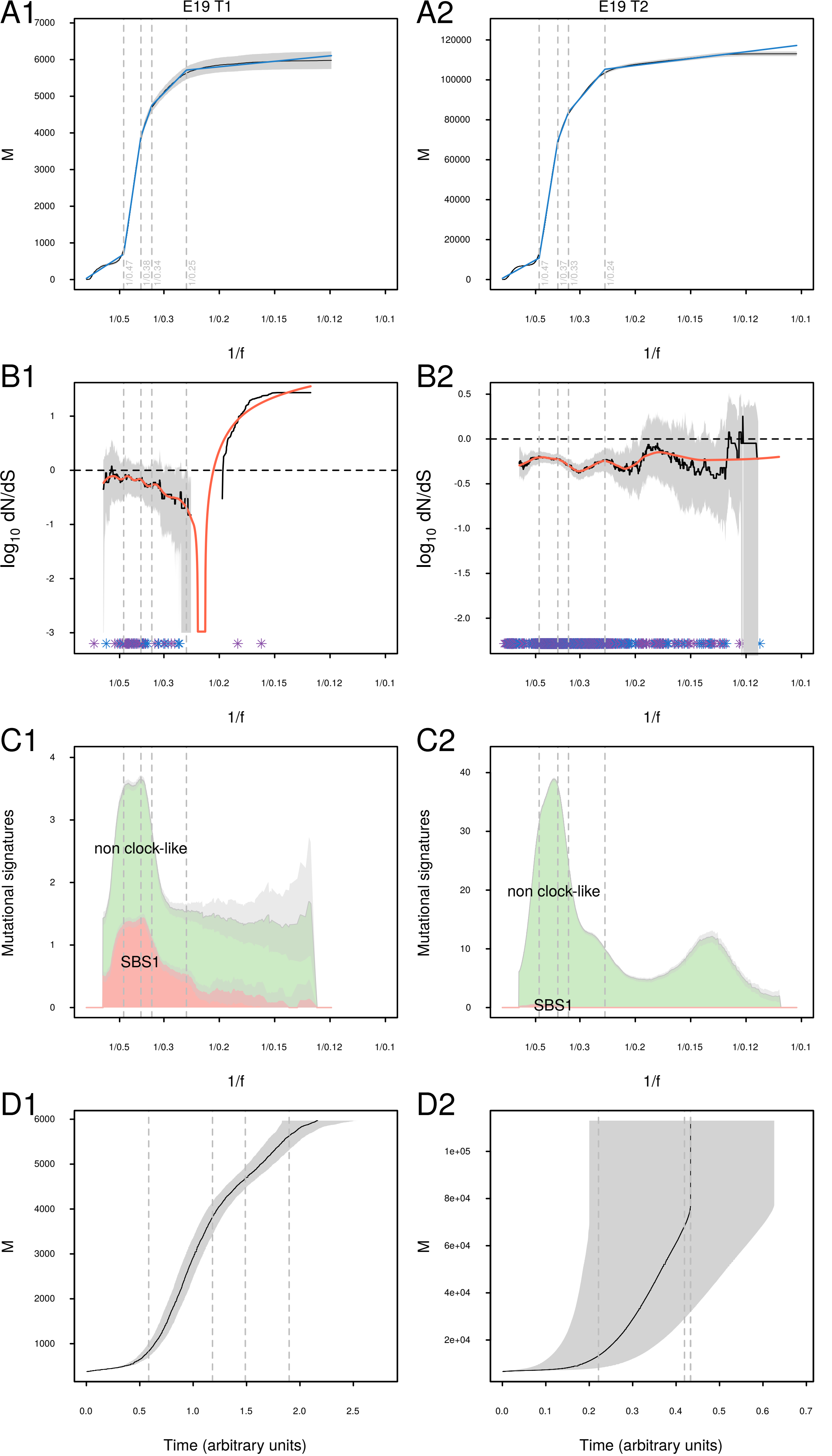

A1
E19 T1
A2
E19 T2
M
1/f
1/0.47
1/0.38
1/0.34
1/0.25
1/0.37
1/0.33
1/0.24
1/0.5
1/0.3
1/0.2
1/0.15
1/0.12
1/0.1
B1
B2
$log_{10}$ dN/dS
C1
C2
Mutational signatures
non clock-like
SBS1
D1
D2
Time (arbitrary units)

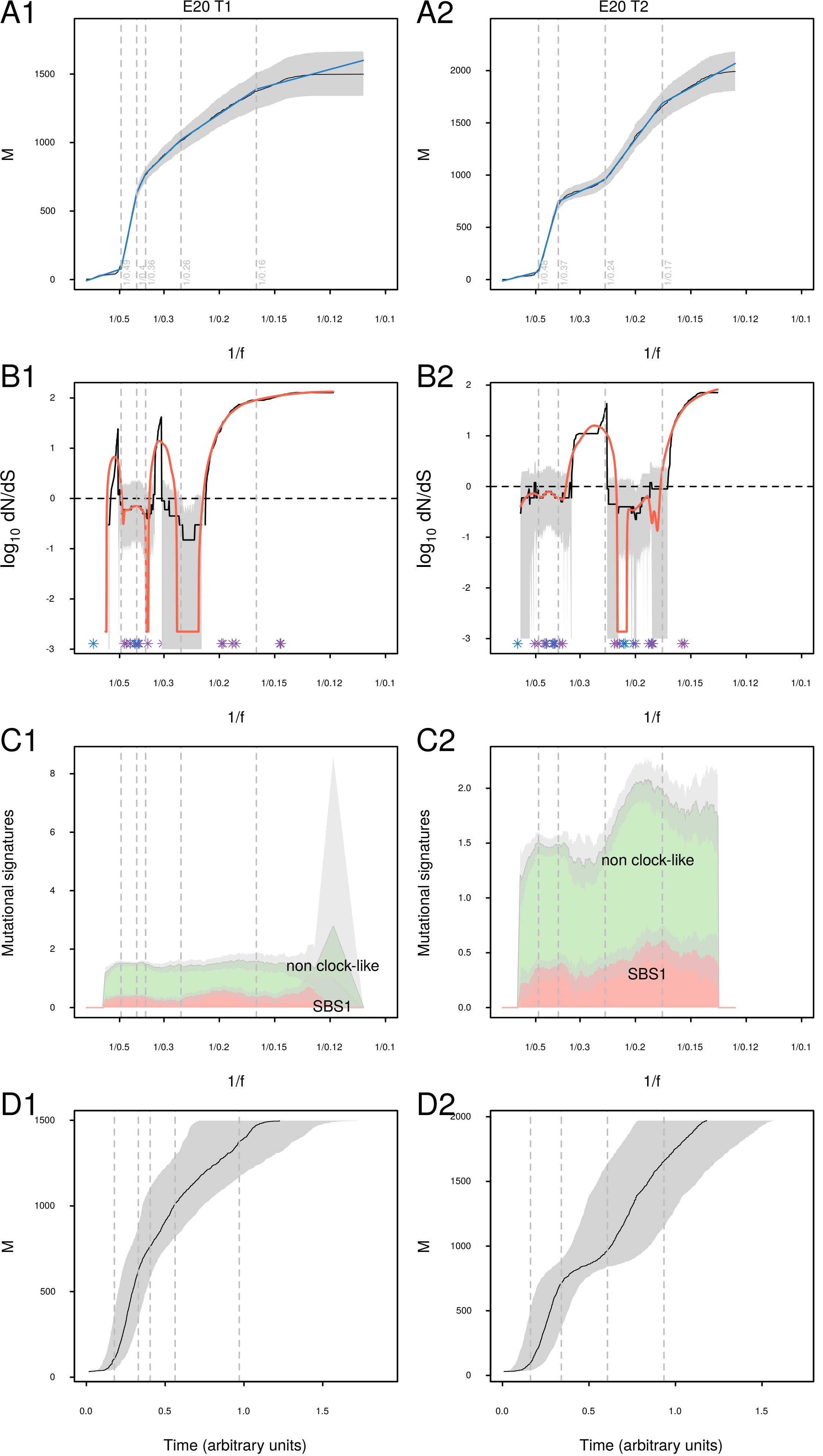

A1
E20 T1
A2
E20 T2
M
1/f
B1
B2
$\log_{10}$ dN/dS
C1
C2
Mutational signatures
non clock-like
SBS1
D1
D2
Time (arbitrary units)

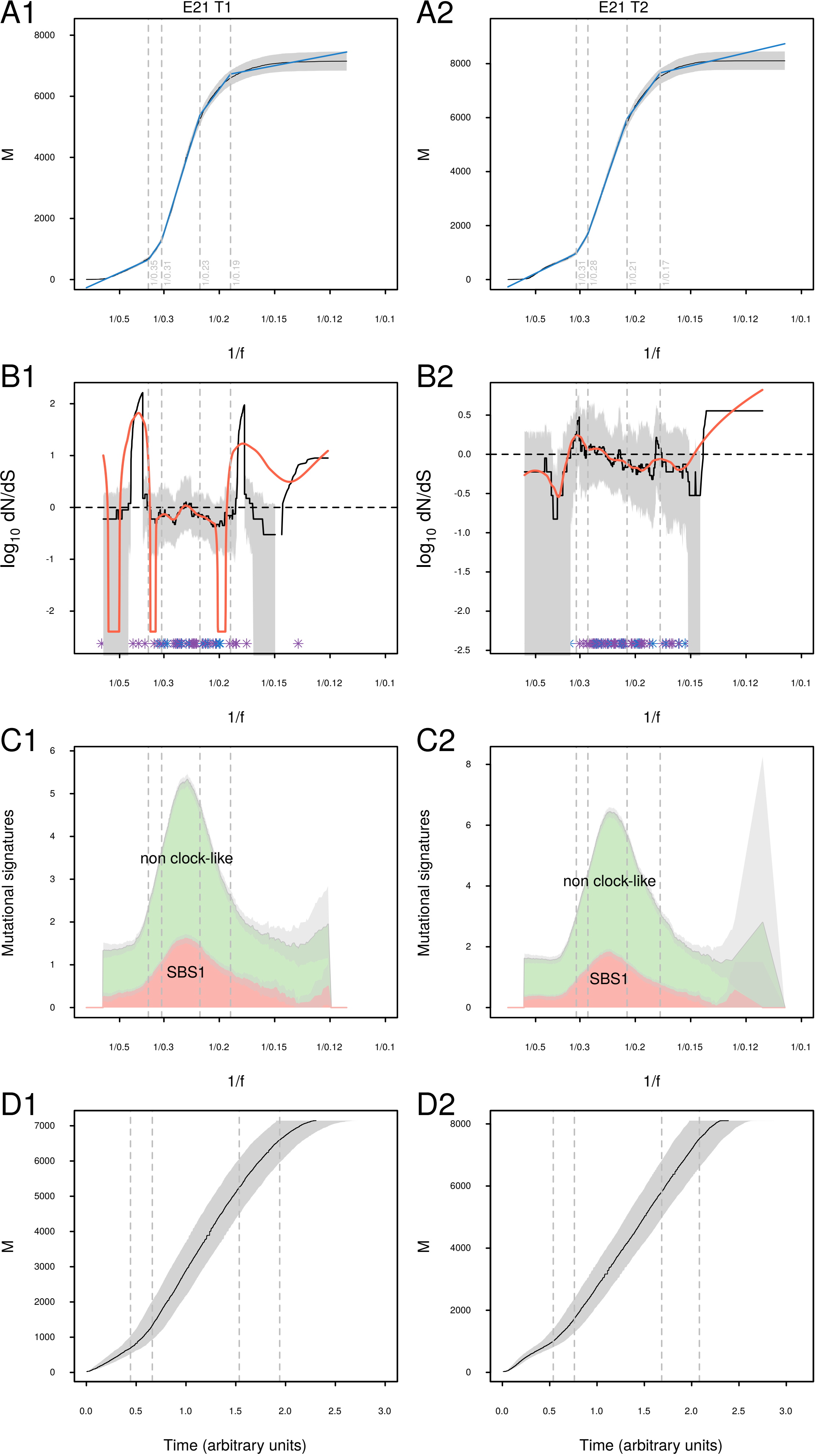

A1
E21 T1
A2
E21 T2
M
1/f
1/0.35
1/0.31
1/0.23
1/0.19
1/0.31
1/0.28
1/0.21
1/0.17
1/0.5
1/0.3
1/0.2
1/0.15
1/0.12
1/0.1
B1
B2
$\log_{10}$ dN/dS
C1
C2
Mutational signatures
non clock-like
SBS1
D1
D2
Time (arbitrary units)

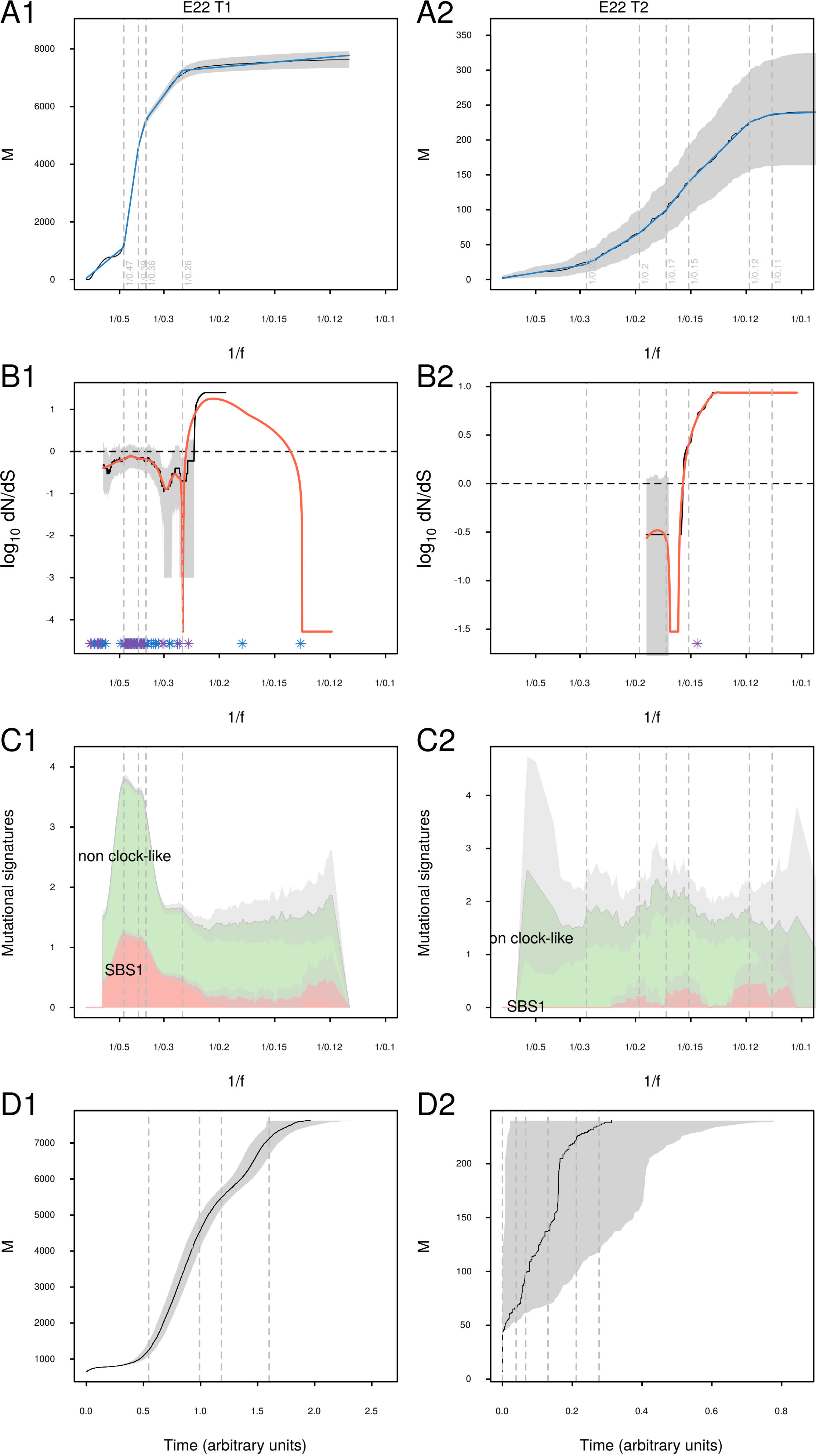

A1
E22 T1
M
1/0.47
1/0.39
1/0.36
1/0.26
1/f
A2
E22 T2
1/0.2
1/0.17
1/0.15
1/0.12
1/0.11
B1
log10 dN/dS
B2
C1
Mutational signatures
non clock-like
SBS1
C2
on clock-like
D1
D2
Time (arbitrary units)

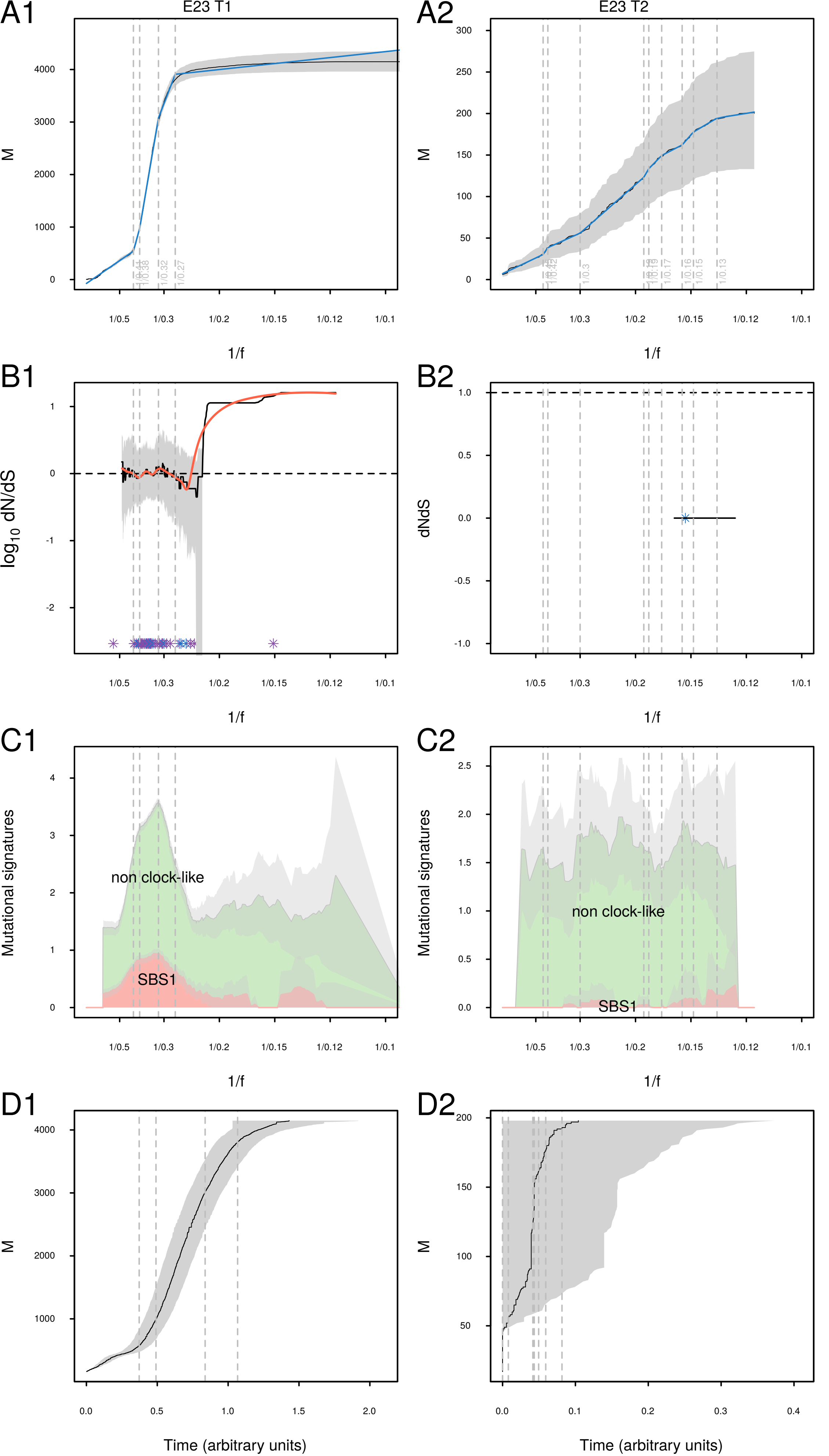
A1
E23 T1
M
1/f
A2
E23 T2
B1
log10 dN/dS
B2
dNdS
C1
Mutational signatures
non clock-like
SBS1
C2
D1
Time (arbitrary units)
D2

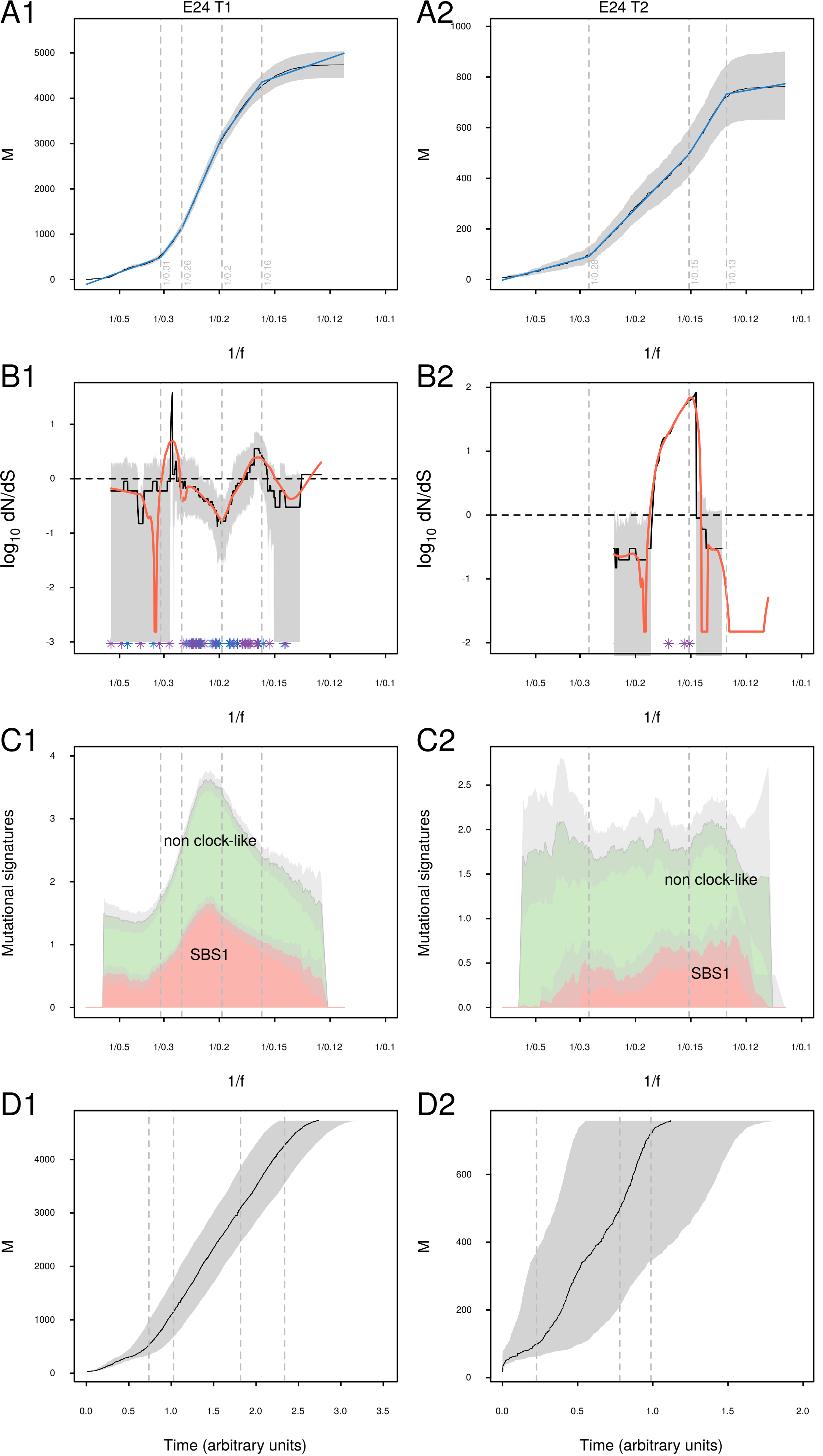

A1
E24 T1
A2
E24 T2
M
1/f
B1
B2
$\log_{10}$ dN/dS
C1
C2
Mutational signatures
non clock-like
SBS1
D1
D2
Time (arbitrary units)

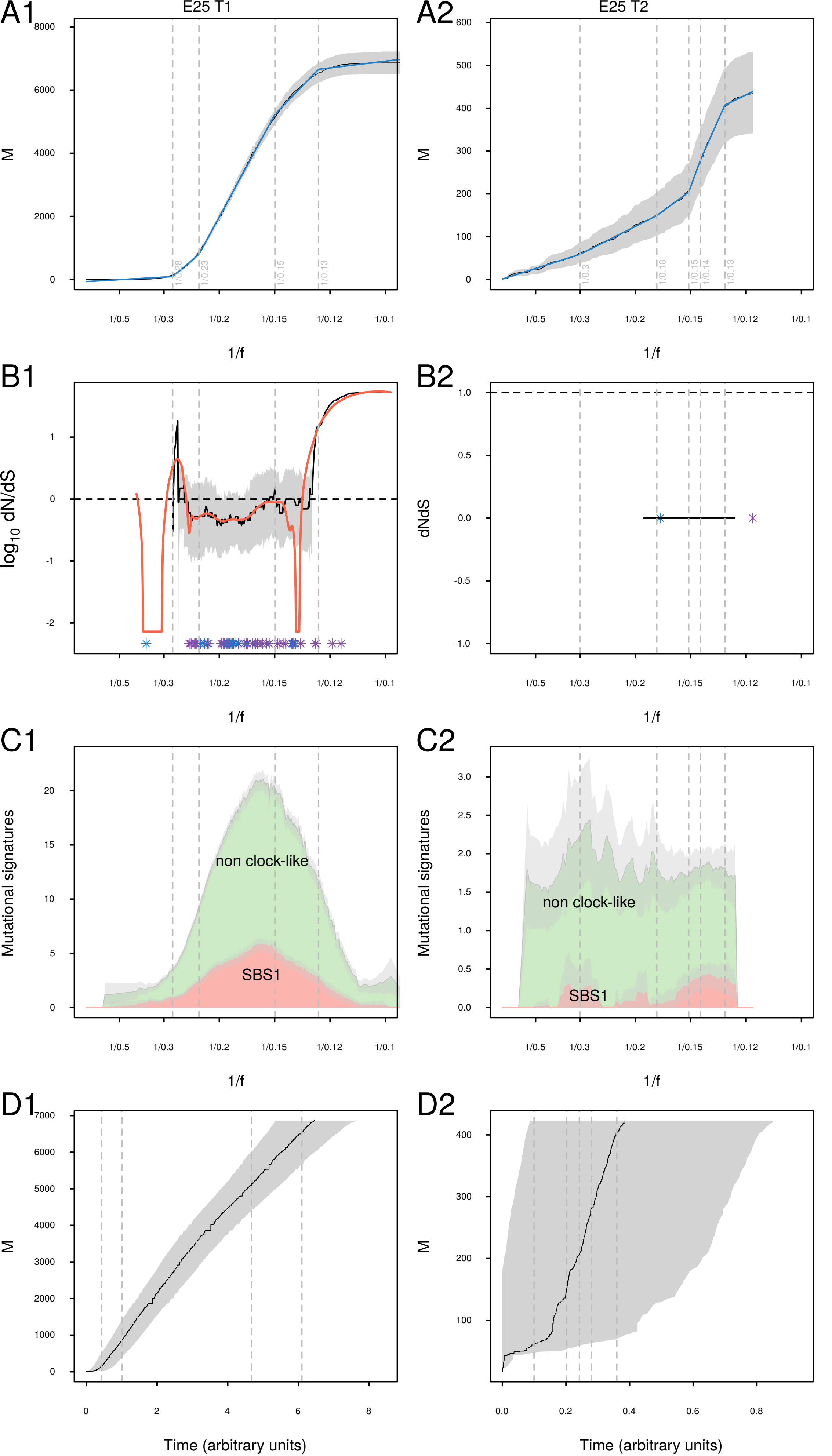

A1
E25 T1
A2
E25 T2
M
1/f
B1
B2
log10 dN/dS
dNdS
C1
C2
Mutational signatures
non clock-like
SBS1
D1
D2
Time (arbitrary units)

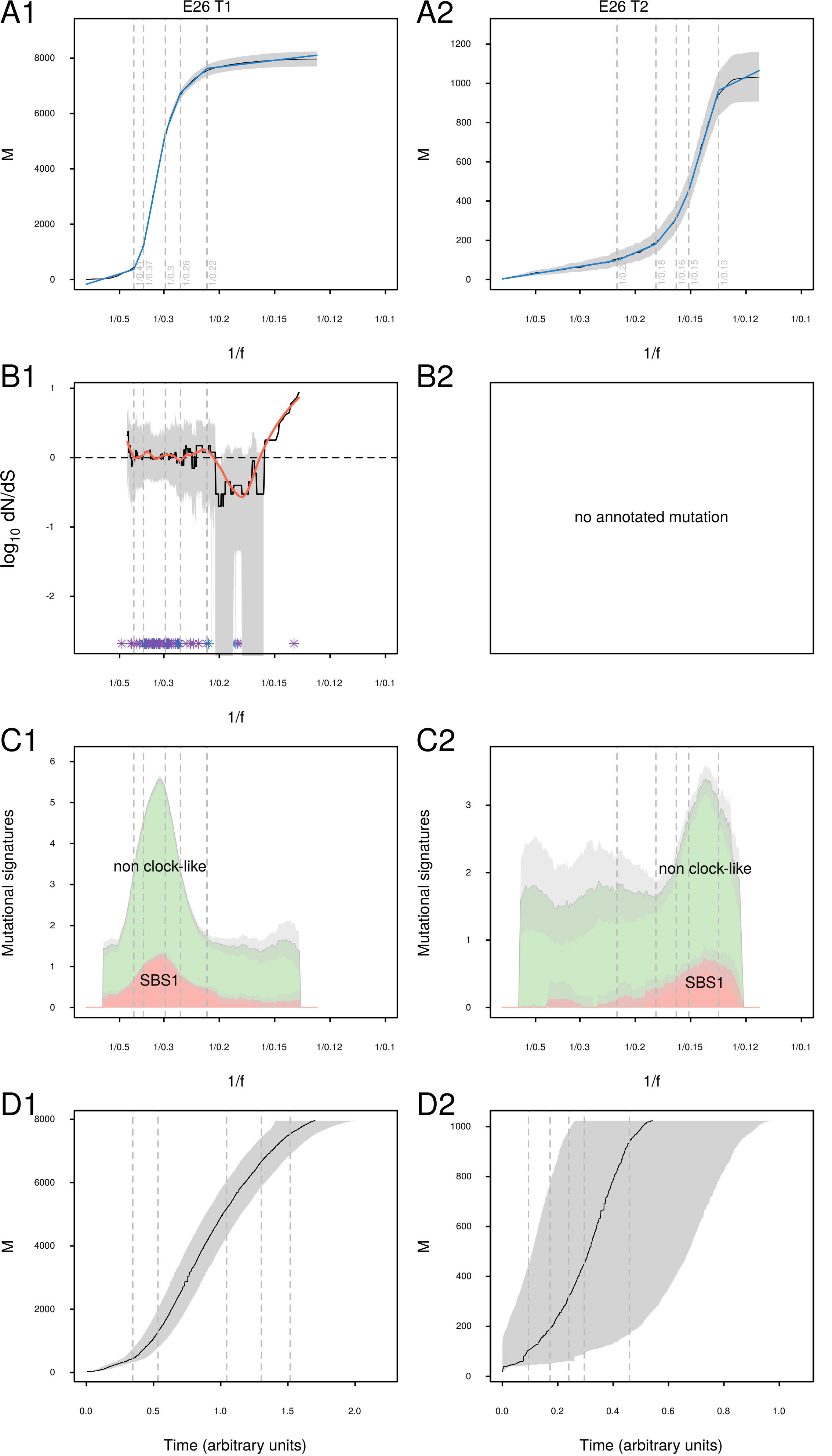

A1
E26 T1
A2
E26 T2
M
1/f
B1
B2
log10 dN/dS
no annotated mutation
C1
C2
Mutational signatures
non clock-like
SBS1
D1
D2
Time (arbitrary units)

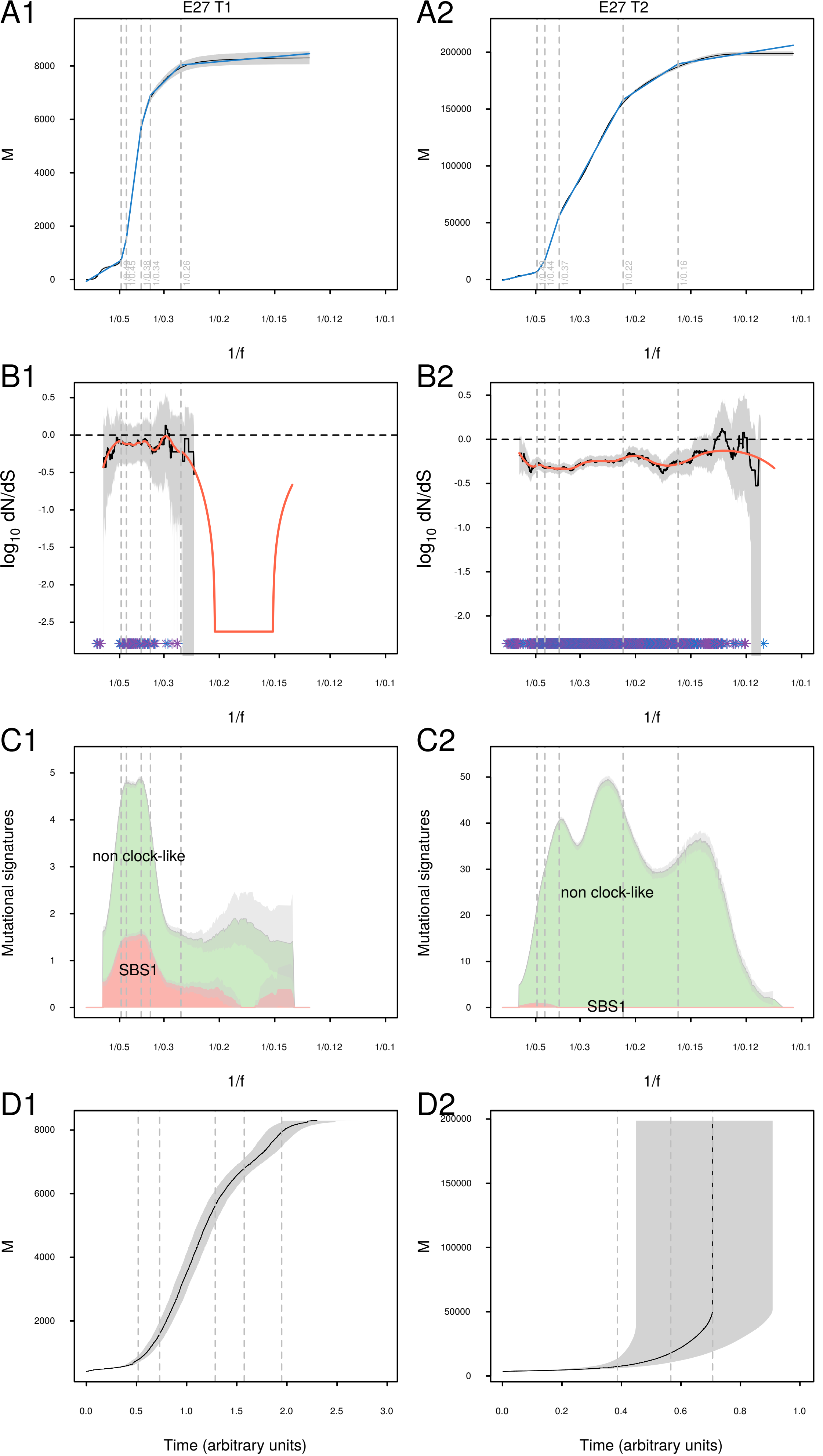

A1
E27 T1
M
1/f
A2
E27 T2
B1
log10 dN/dS
B2
C1
Mutational signatures
non clock-like
SBS1
C2
D1
Time (arbitrary units)
D2

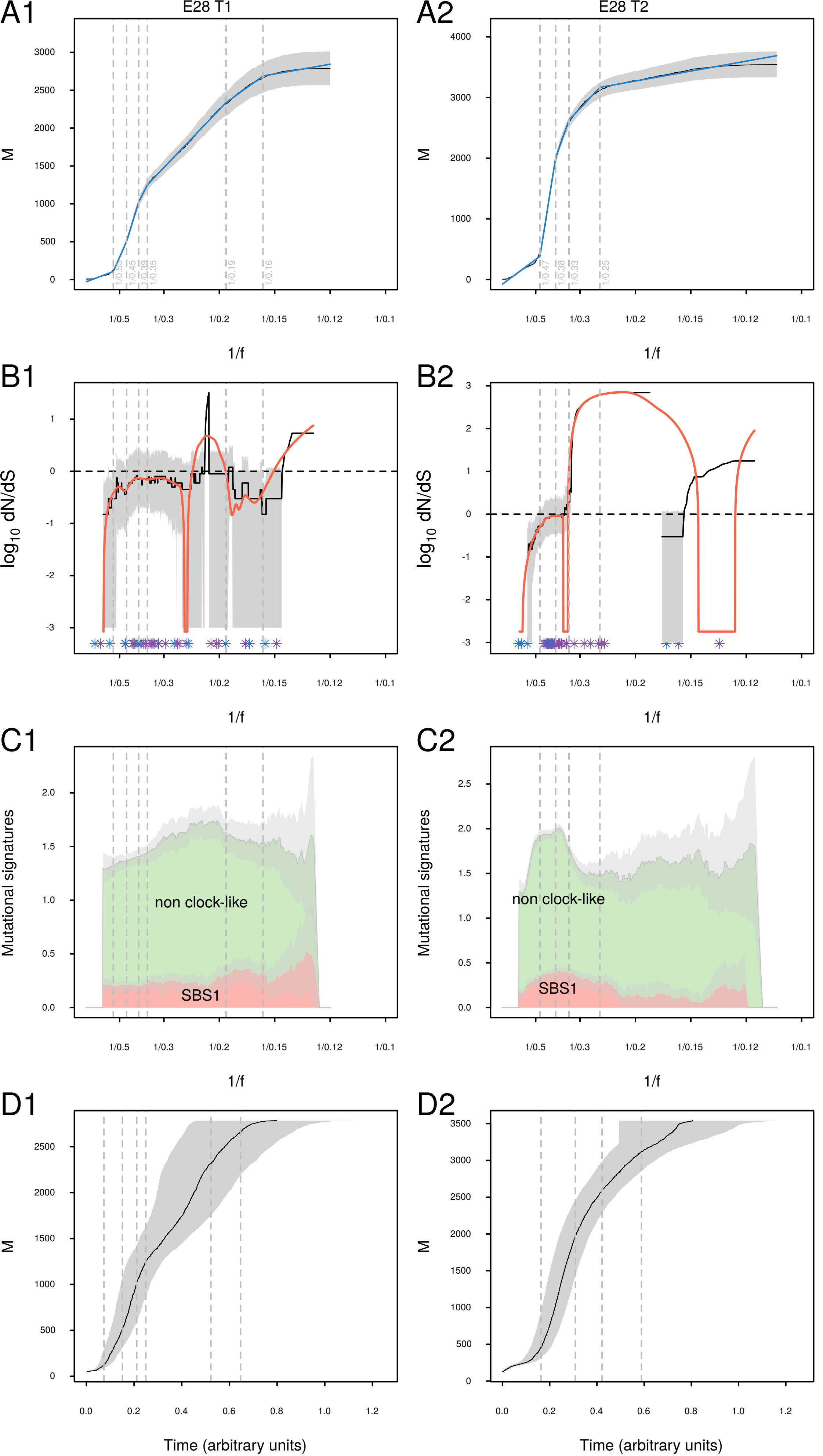

A1
E28 T1
A2
E28 T2
M
1/f
B1
B2
log10 dN/dS
C1
C2
Mutational signatures
non clock-like
SBS1
D1
D2
Time (arbitrary units)

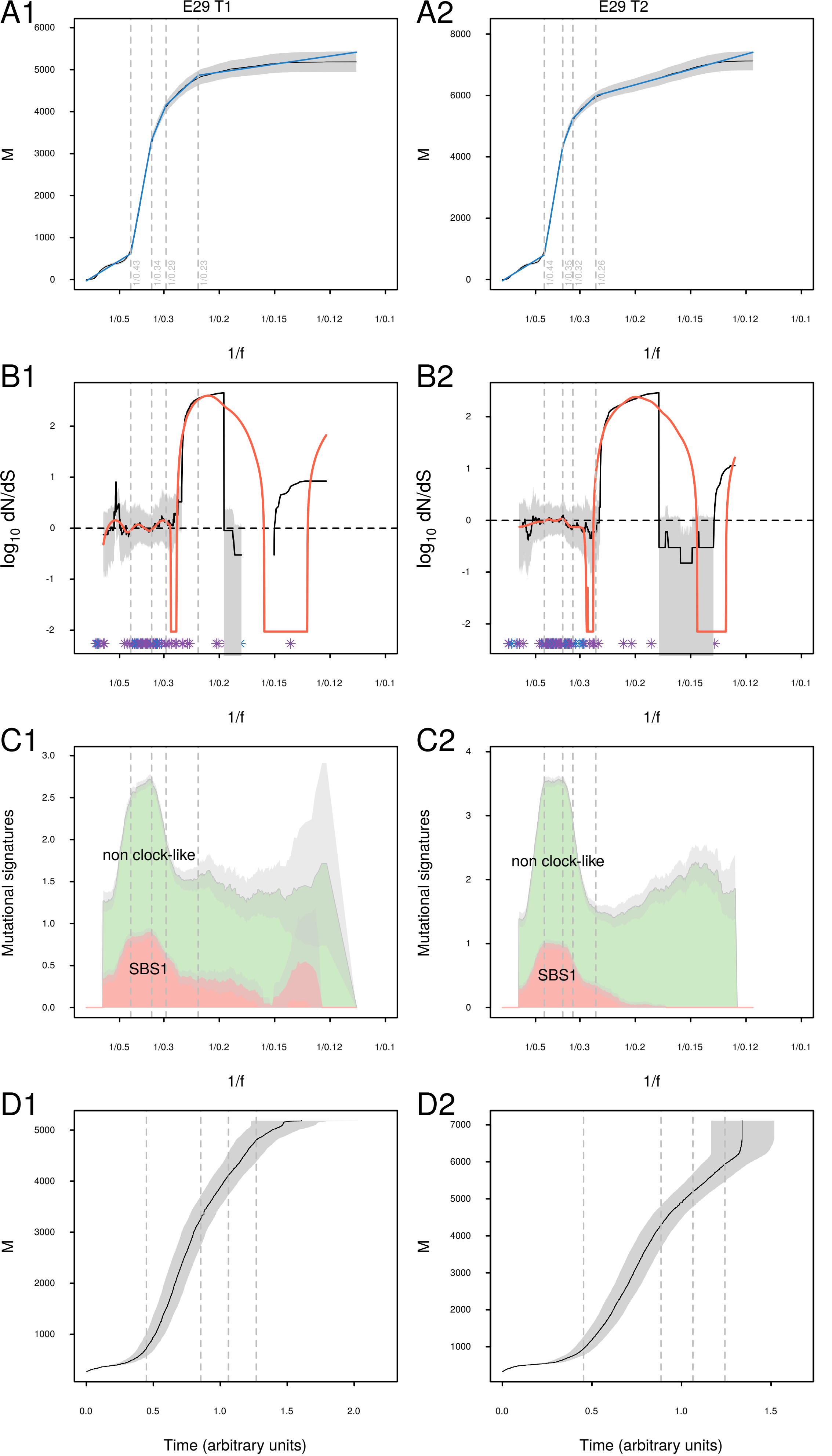
A1
E29 T1
M
1/f
A2
E29 T2
B1
log10 dN/dS
B2
C1
Mutational signatures
non clock-like
SBS1
C2
D1
Time (arbitrary units)
D2

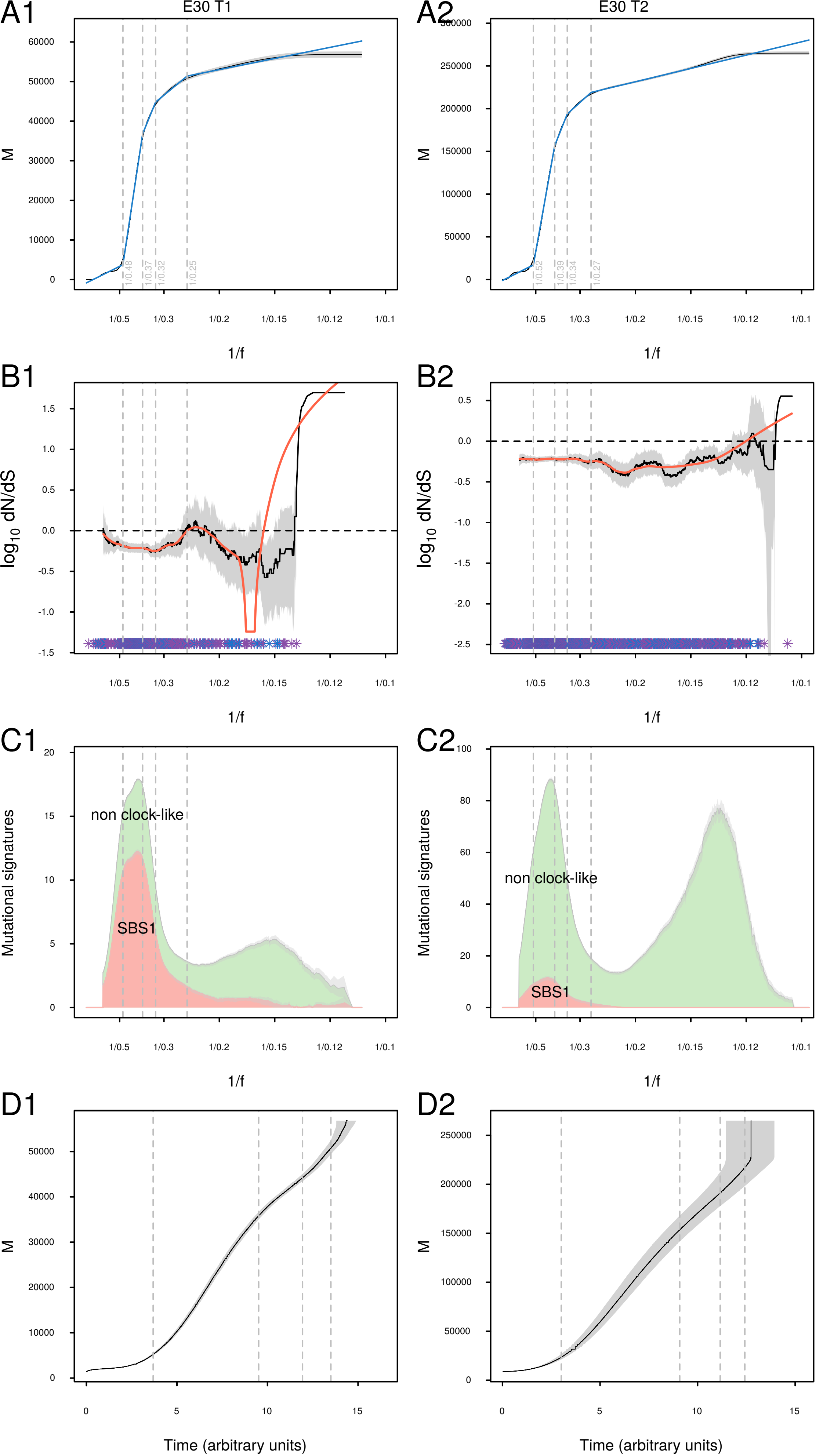

A1
E30 T1
M
1/f
A2
E30 T2
B1
log10 dN/dS
B2
C1
non clock-like
SBS1
Mutational signatures
C2
D1
Time (arbitrary units)
D2

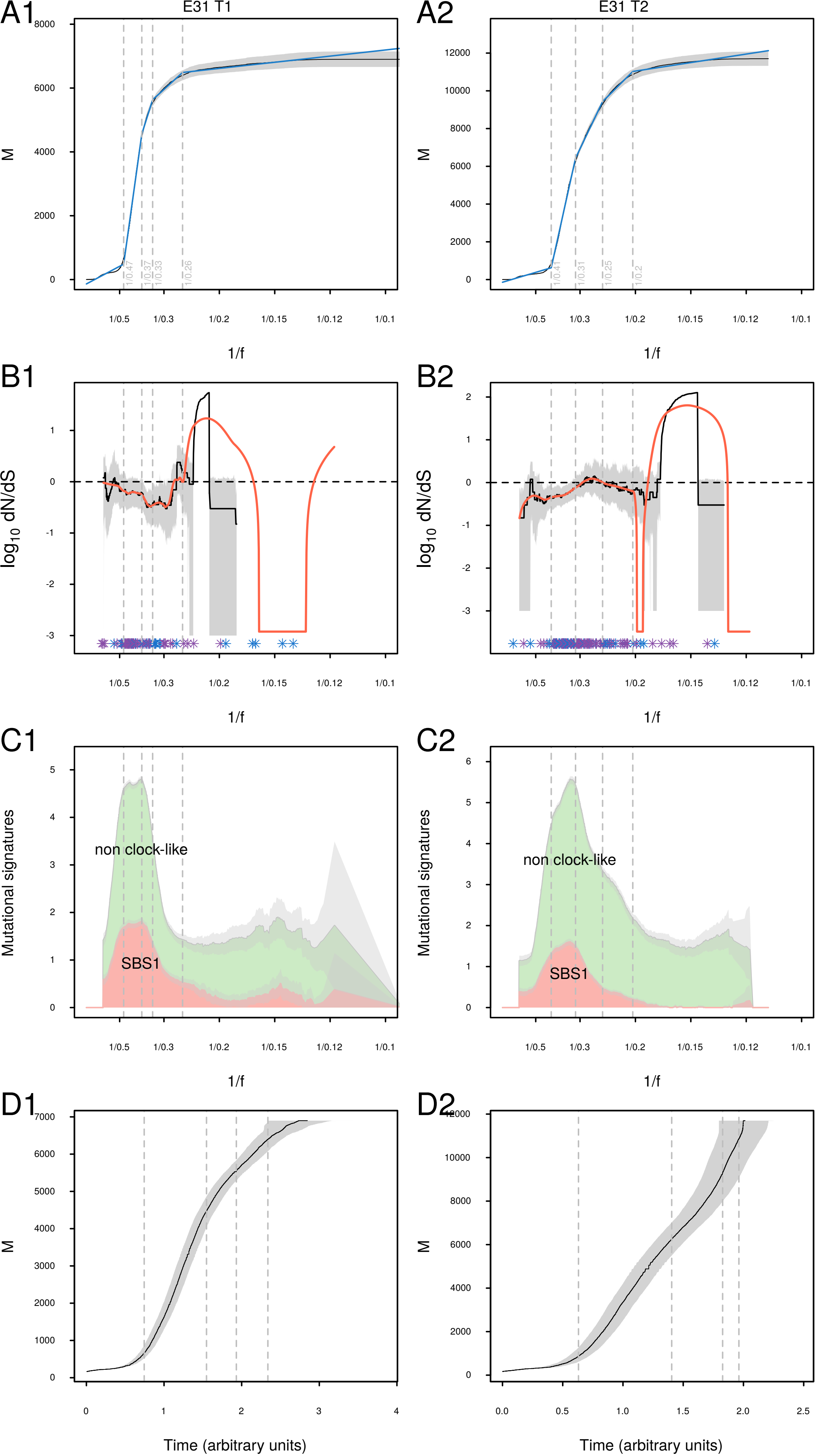
A1
E31 T1
A2
E31 T2
M
1/f
B1
B2
log10 dN/dS
C1
C2
Mutational signatures
non clock-like
SBS1
D1
D2
Time (arbitrary units)

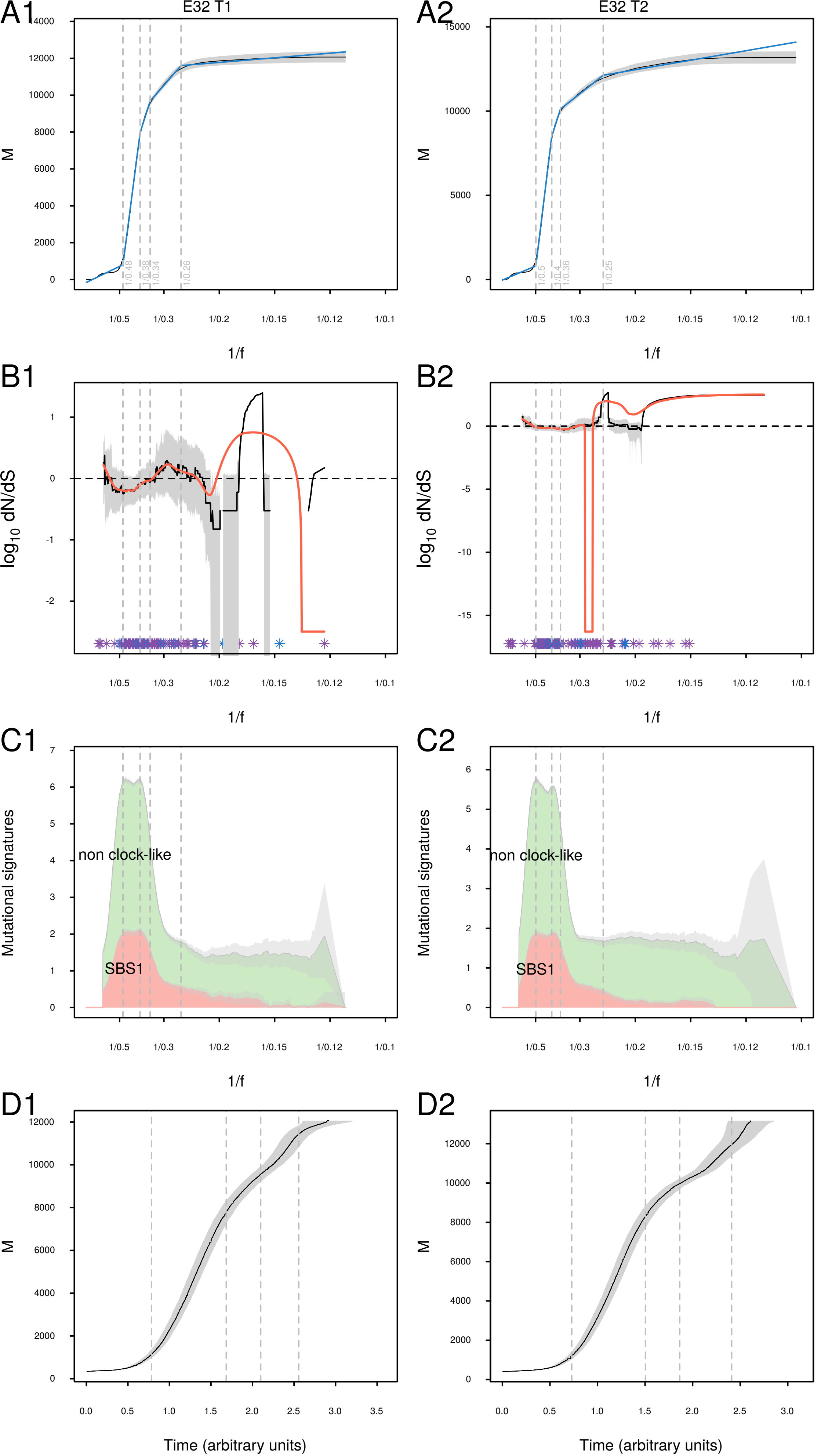

A1
E32 T1
A2
E32 T2
M
1/f
B1
B2
log10 dN/dS
C1
C2
Mutational signatures
non clock-like
SBS1
D1
D2
Time (arbitrary units)

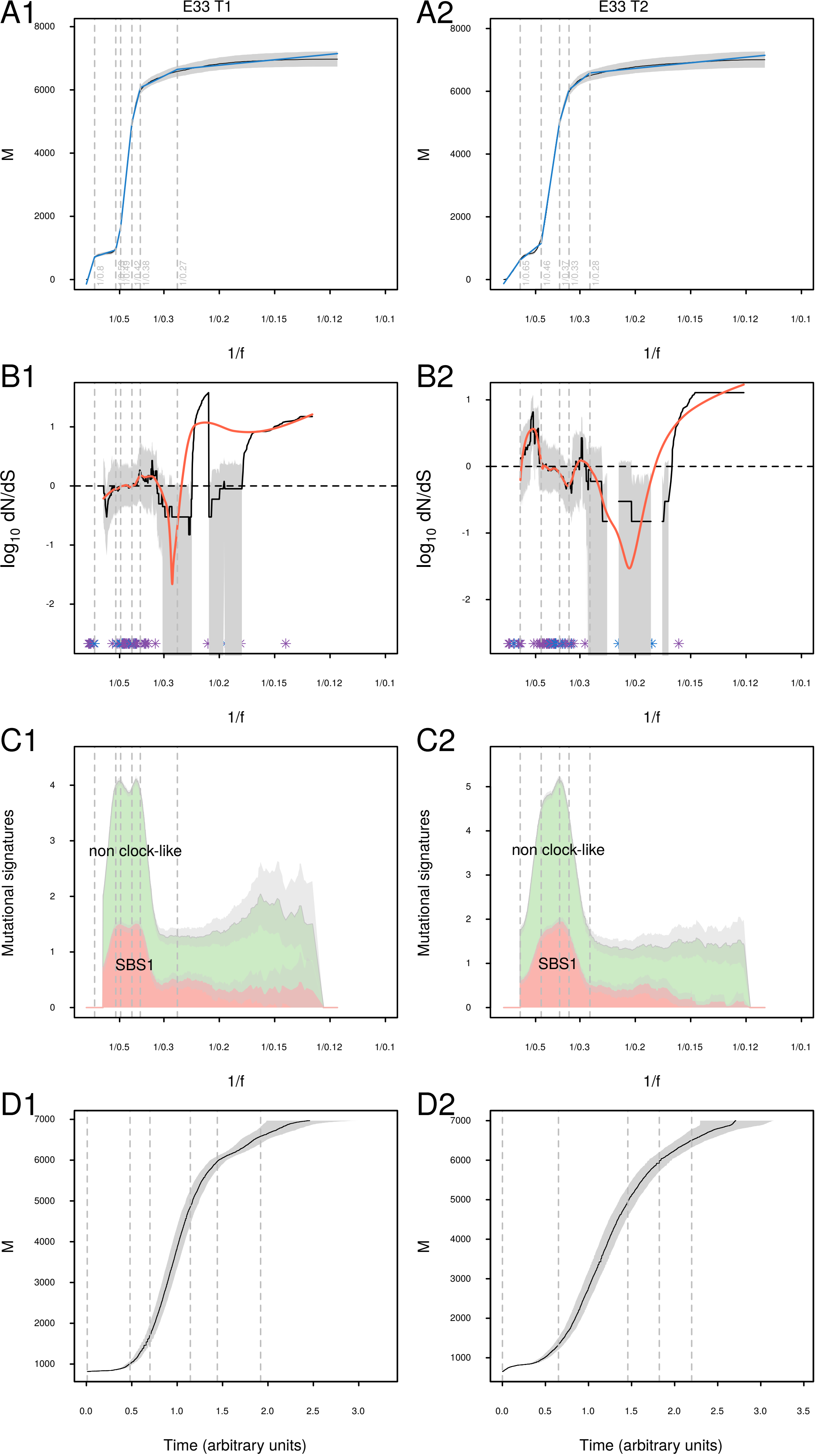
A1
E33 T1
A2
E33 T2
M
1/f
B1
B2
log10 dN/dS
C1
C2
Mutational signatures
non clock-like
SBS1
D1
D2
Time (arbitrary units)

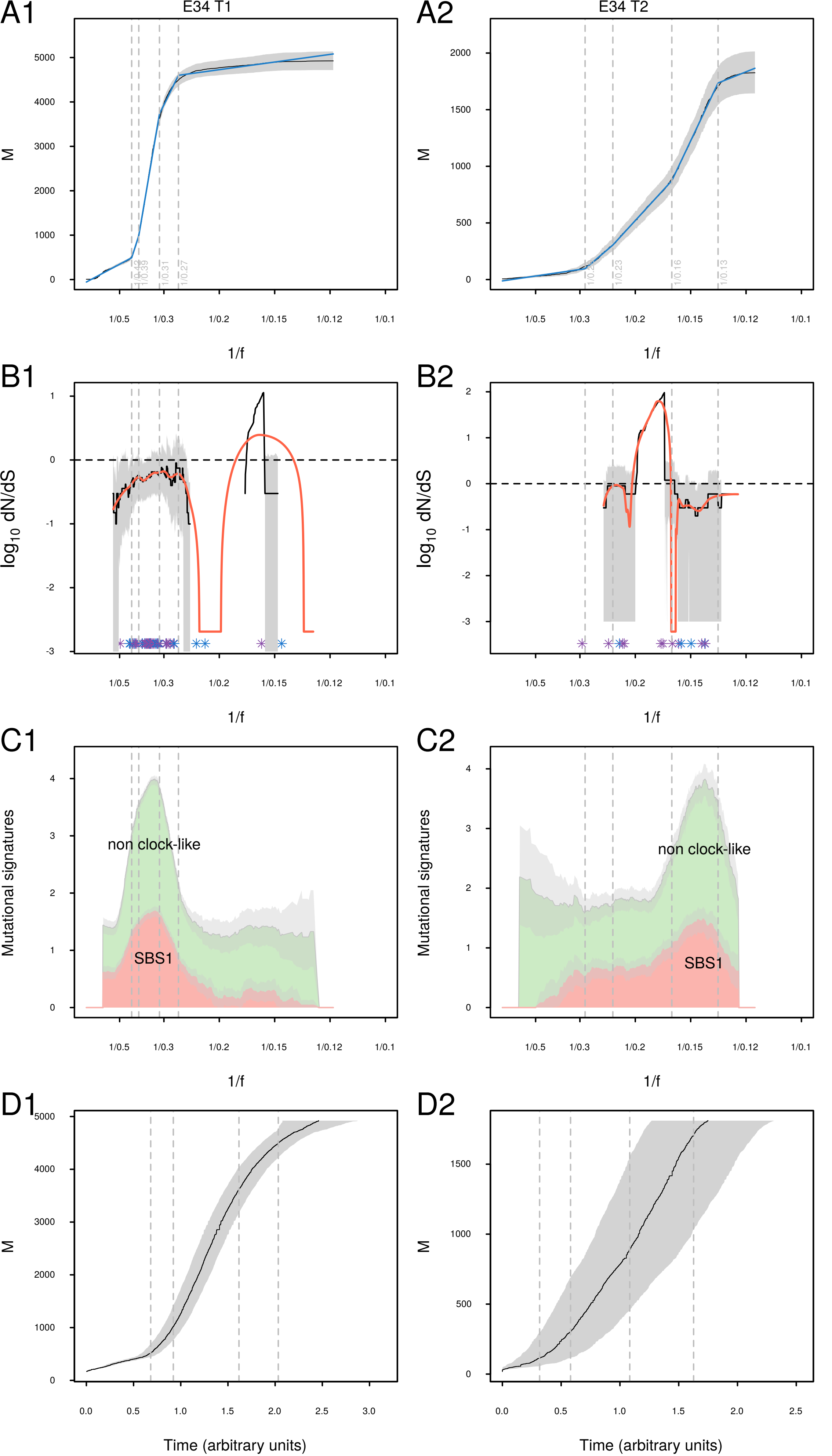
A1
E34 T1
A2
E34 T2
M
1/f
B1
B2
log10 dN/dS
C1
C2
Mutational signatures
non clock-like
SBS1
D1
D2
Time (arbitrary units)

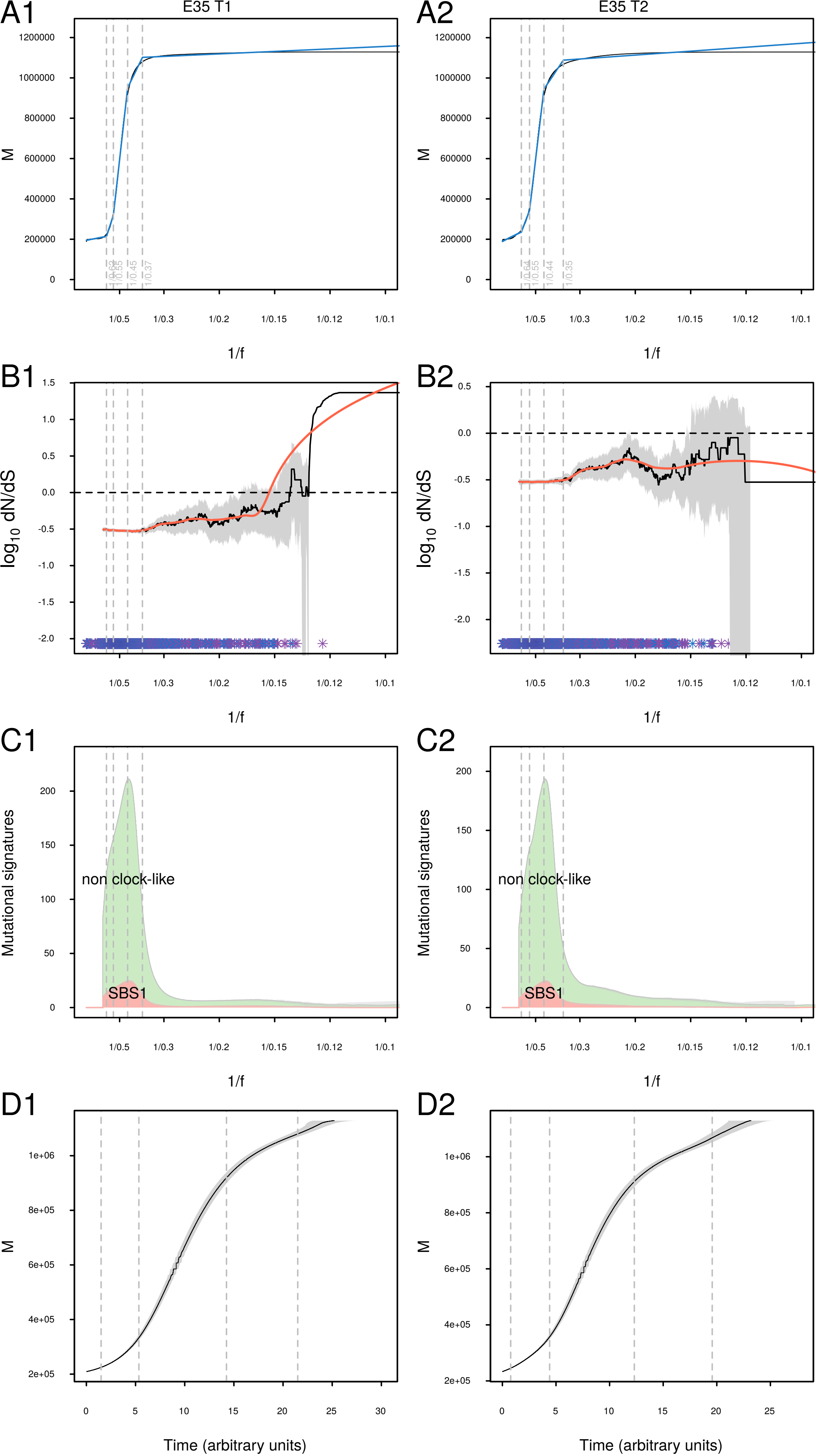
A1
E35 T1
A2
E35 T2
M
1/f
B1
B2
log10 dN/dS
C1
C2
Mutational signatures
non clock-like
SBS1
D1
D2
Time (arbitrary units)

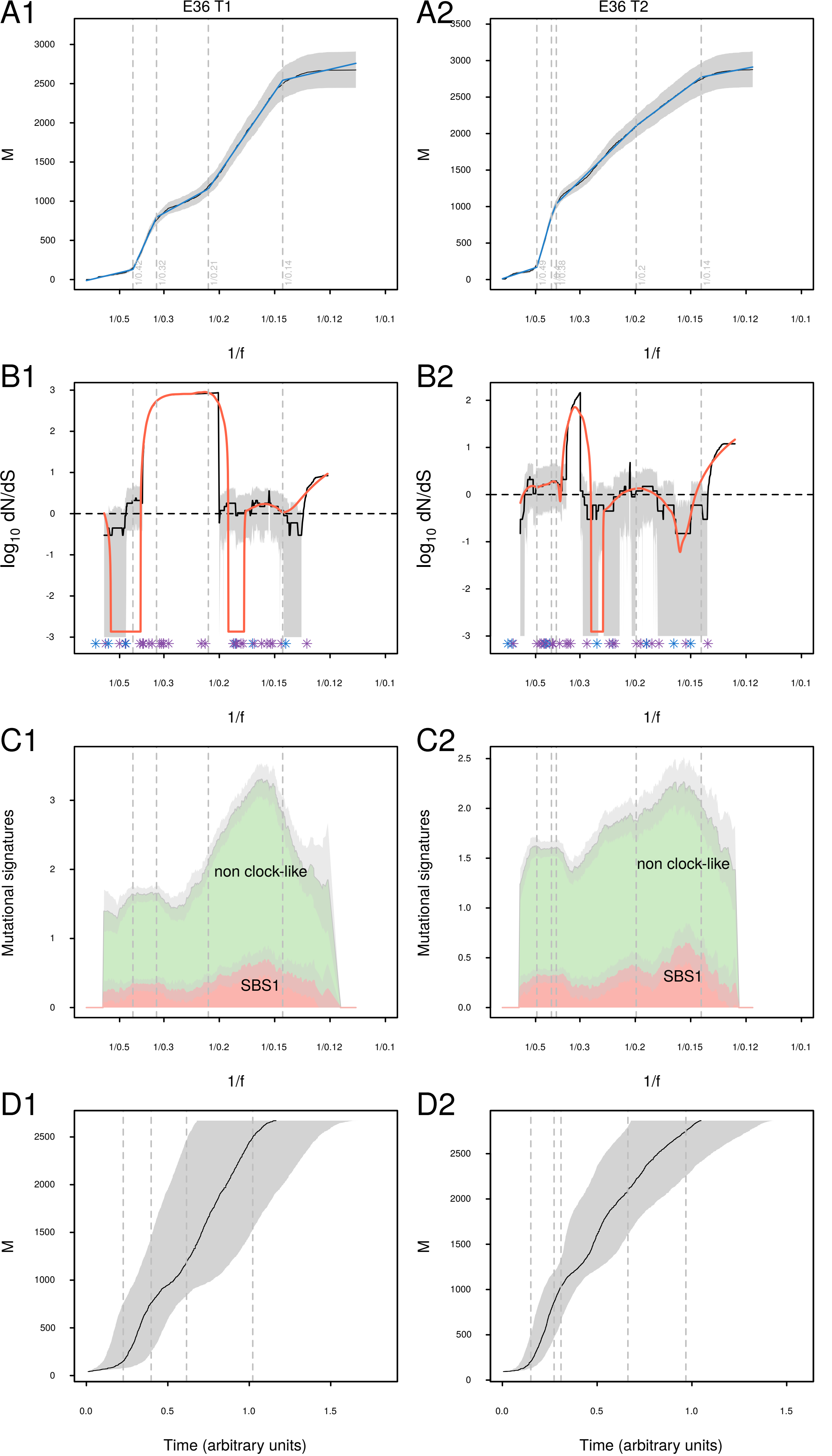

A1
E36 T1
A2
E36 T2
M
1/f
B1
B2
log10 dN/dS
C1
C2
Mutational signatures
non clock-like
SBS1
D1
D2
Time (arbitrary units)

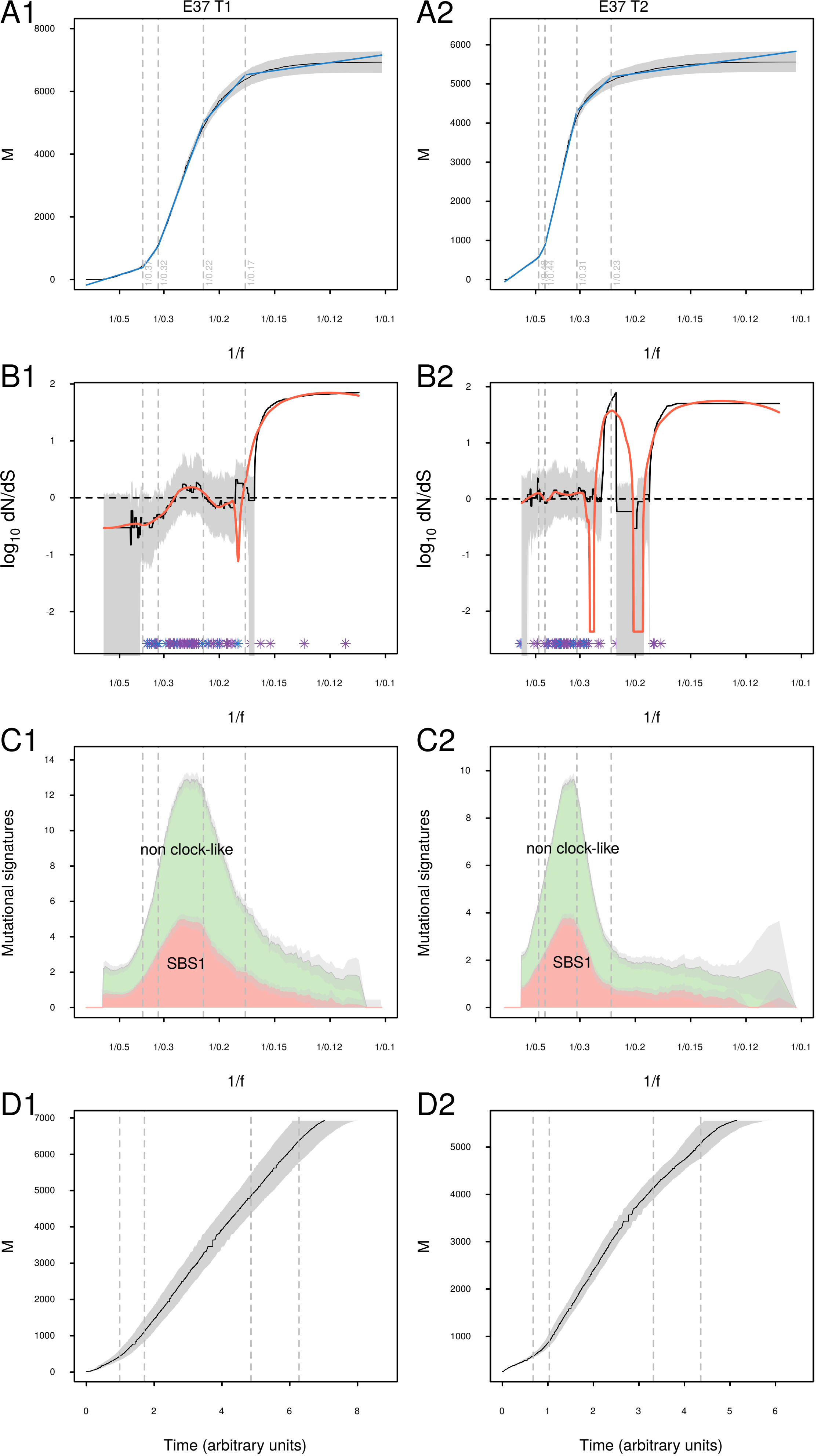

A1
E37 T1
M
1/f
A2
E37 T2
B1
log10 dN/dS
B2
C1
Mutational signatures
non clock-like
SBS1
C2
D1
Time (arbitrary units)
D2

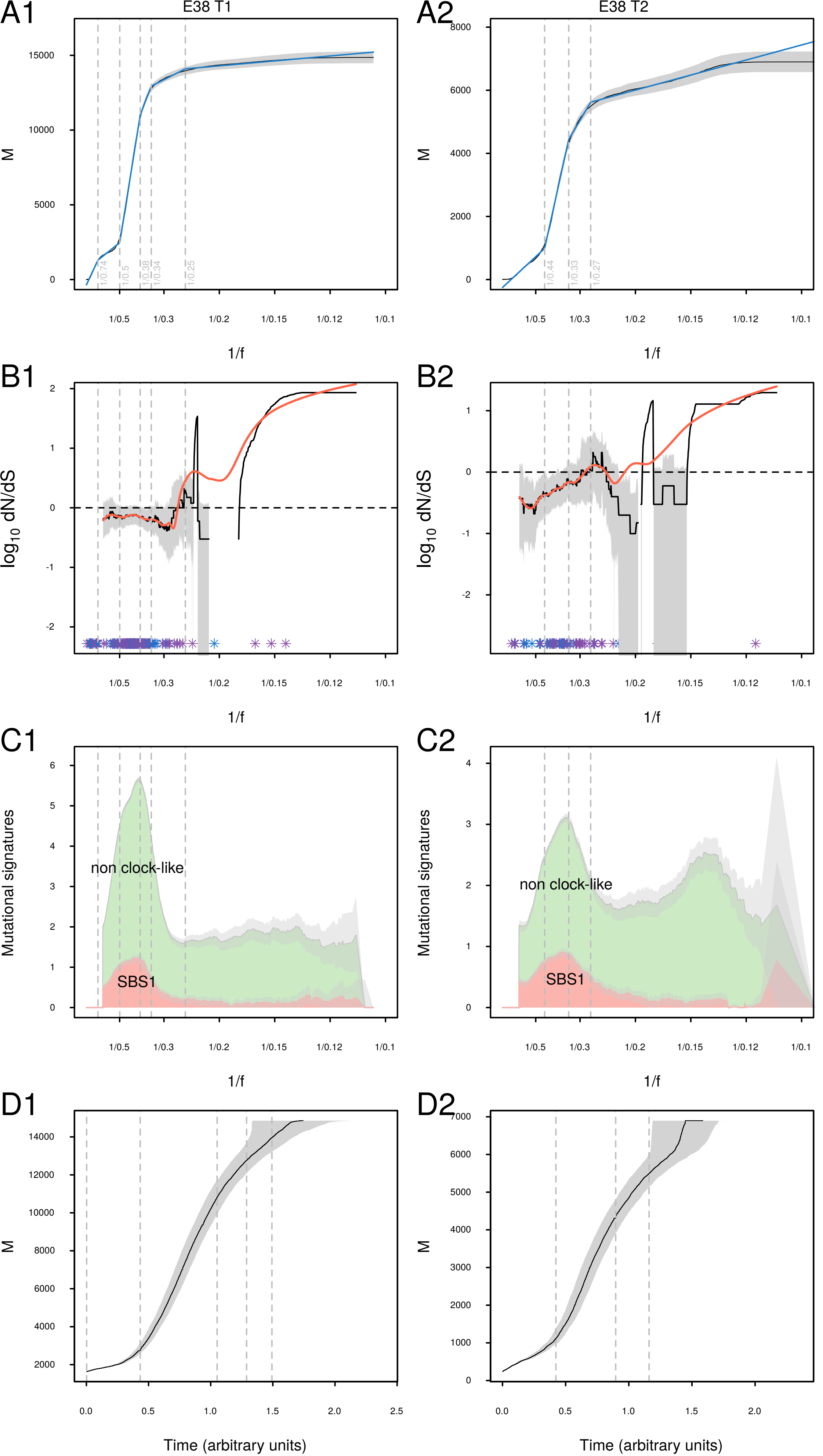

A1
E38 T1
A2
E38 T2
M
1/f
B1
B2
log10 dN/dS
C1
C2
Mutational signatures
non clock-like
SBS1
D1
D2
Time (arbitrary units)

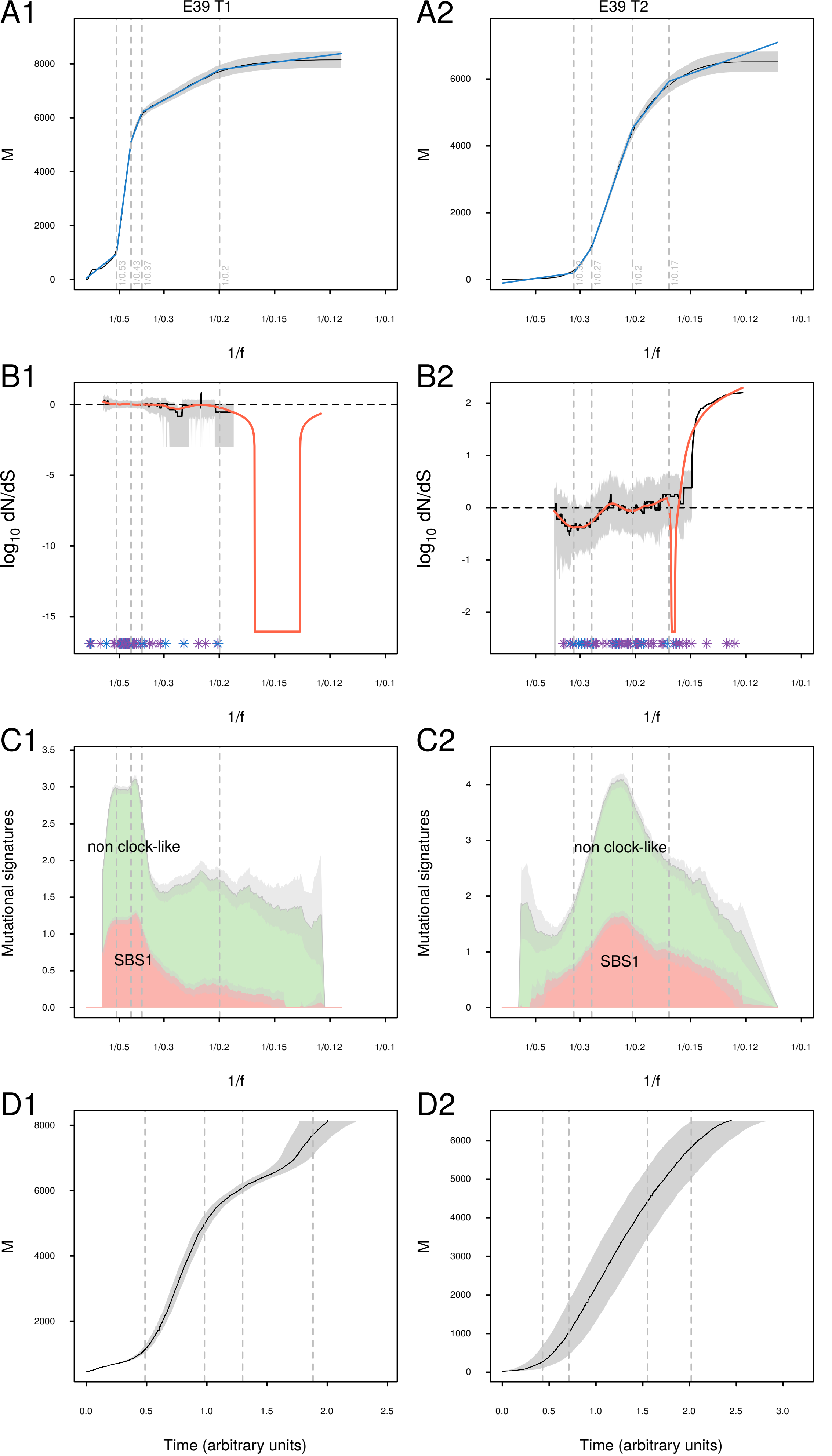

A1
E39 T1
A2
E39 T2
M
1/f
B1
B2
log10 dN/dS
C1
C2
non clock-like
SBS1
Mutational signatures
D1
D2
Time (arbitrary units)

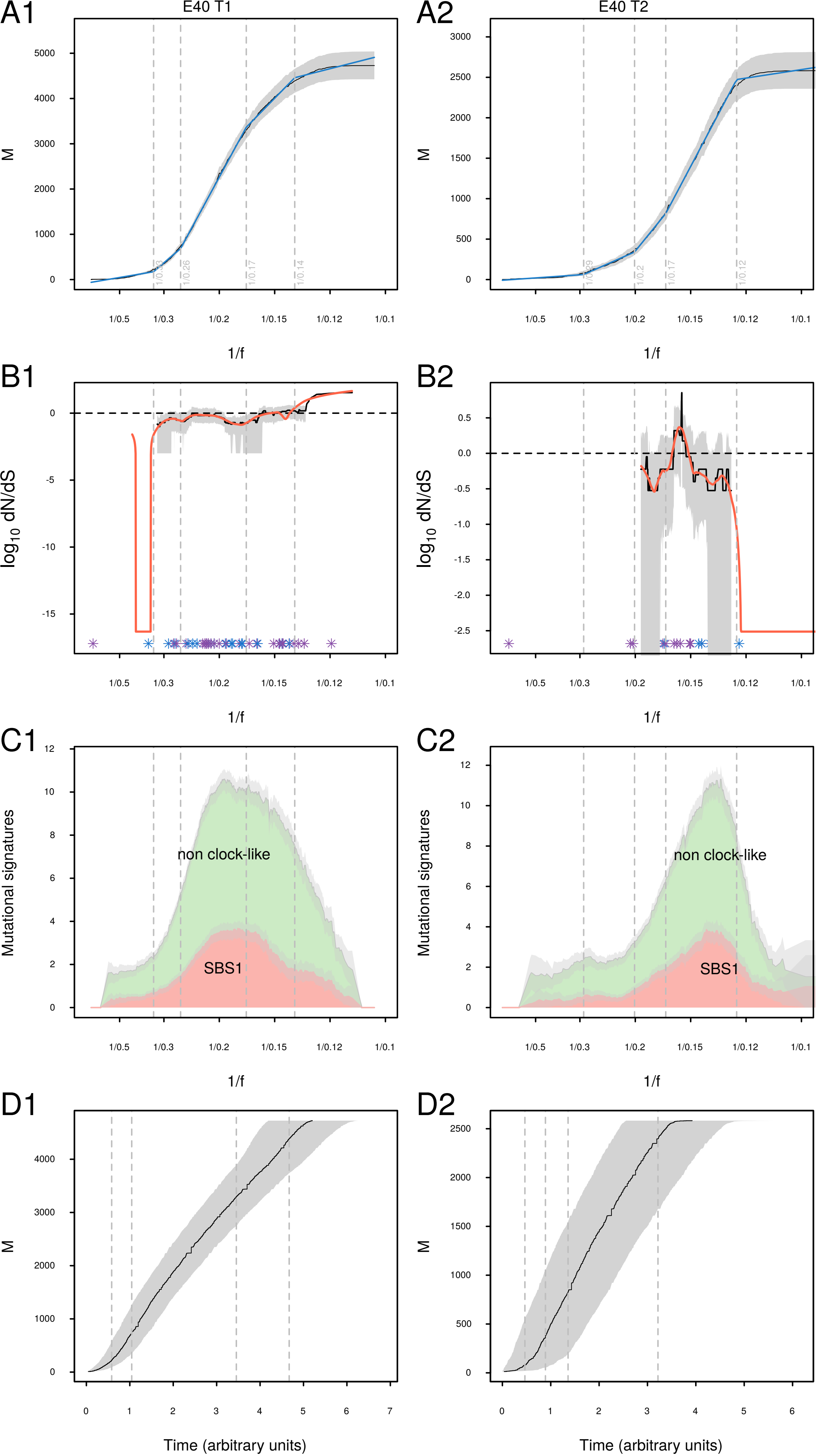

A1
E40 T1
A2
E40 T2
M
1/f
B1
B2
log10 dN/dS
C1
C2
Mutational signatures
non clock-like
SBS1
D1
D2
Time (arbitrary units)

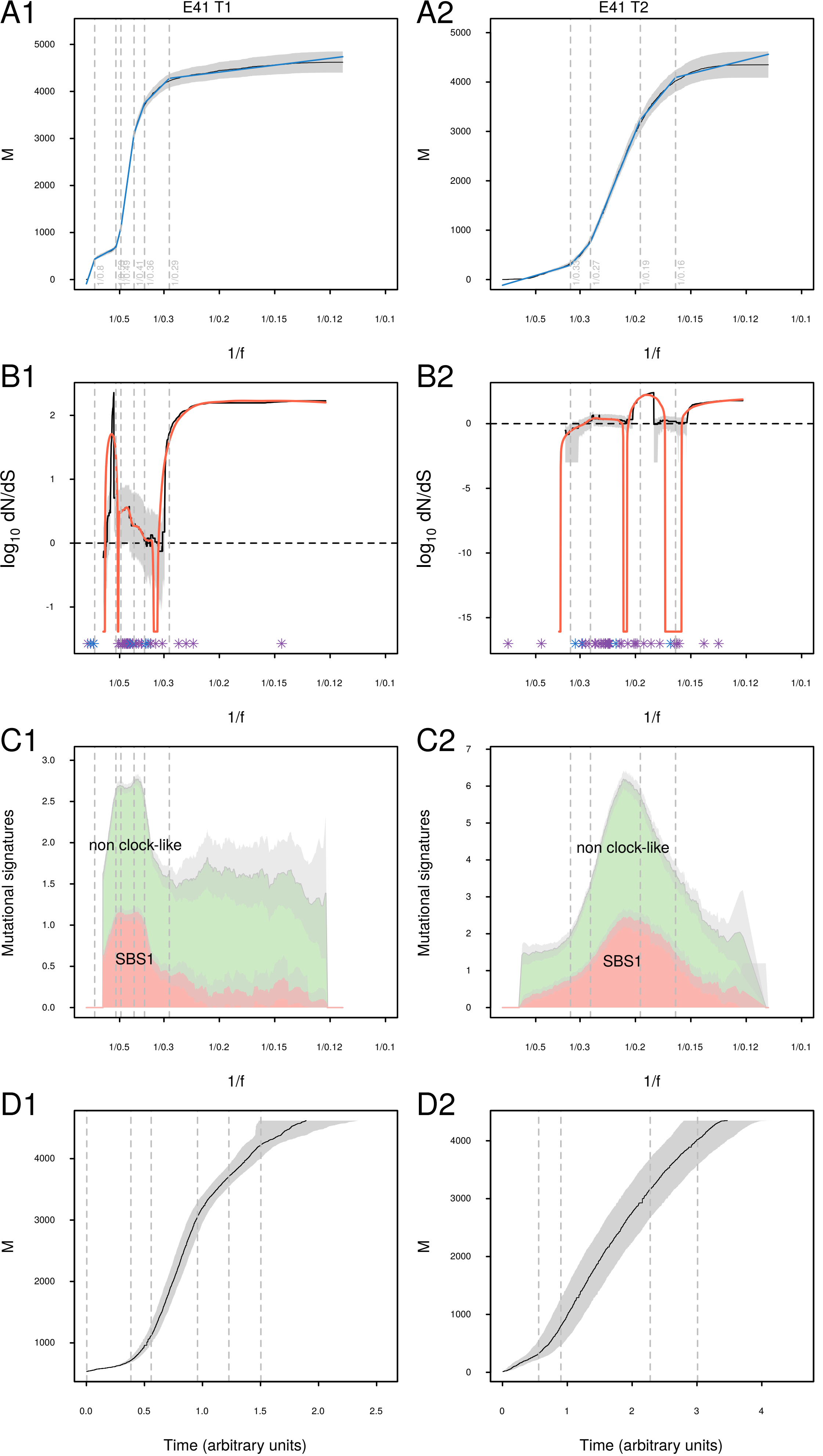

A1
E41 T1
A2
E41 T2
M
1/f
B1
B2
$\log_{10}$ dN/dS
C1
C2
Mutational signatures
non clock-like
SBS1
D1
D2
Time (arbitrary units)

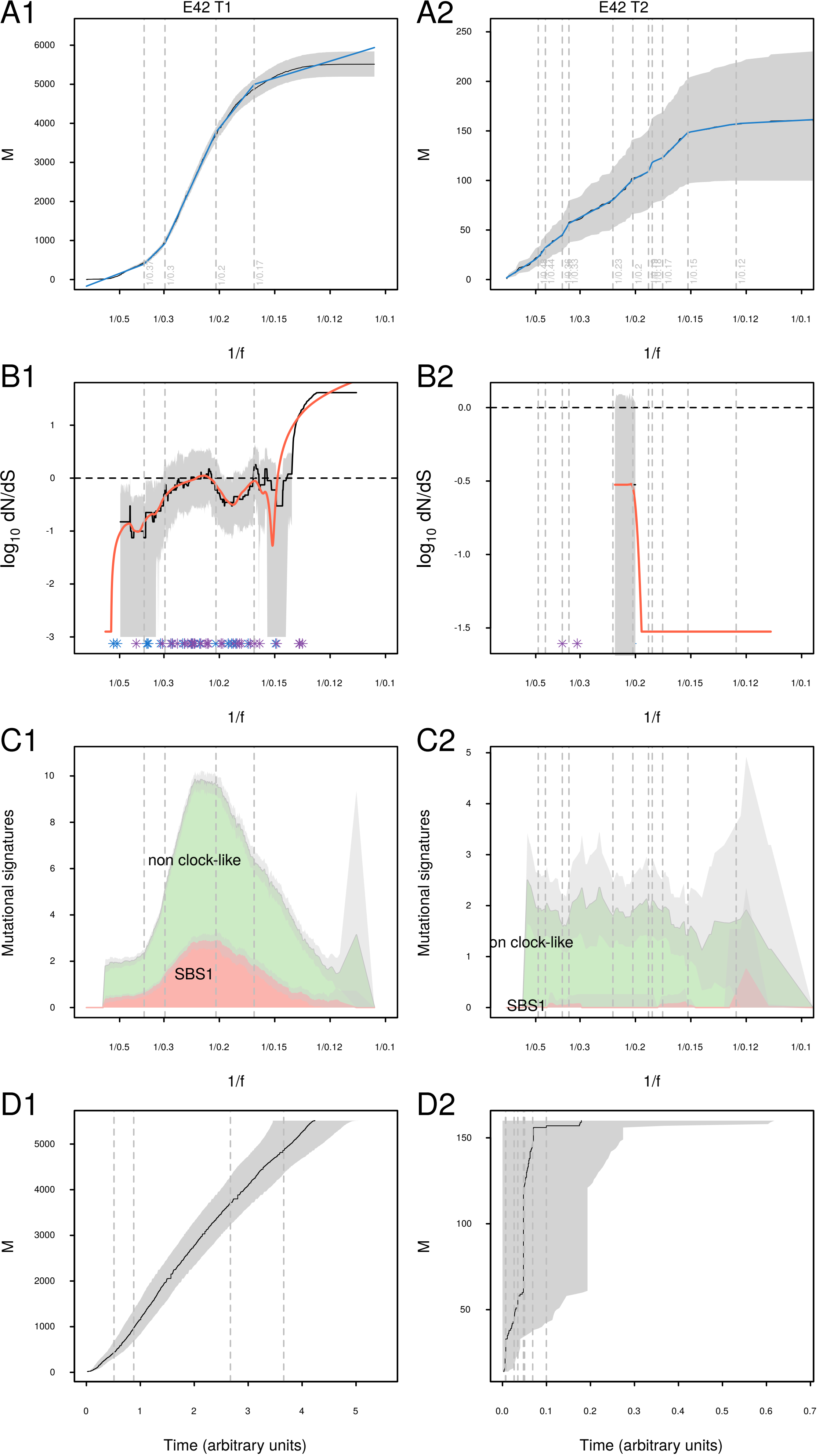
A1
E42 T1
A2
E42 T2
B1
B2
C1
C2
non clock-like
SBS1
on clock-like
D1
D2
M
log10 dN/dS
Mutational signatures
1/f
Time (arbitrary units)
1/0.5
1/0.3
1/0.2
1/0.15
1/0.12
1/0.1
