## Supplemental Figure 2 for "Time-resolved, integrated analysis of clonally evolving genomes"

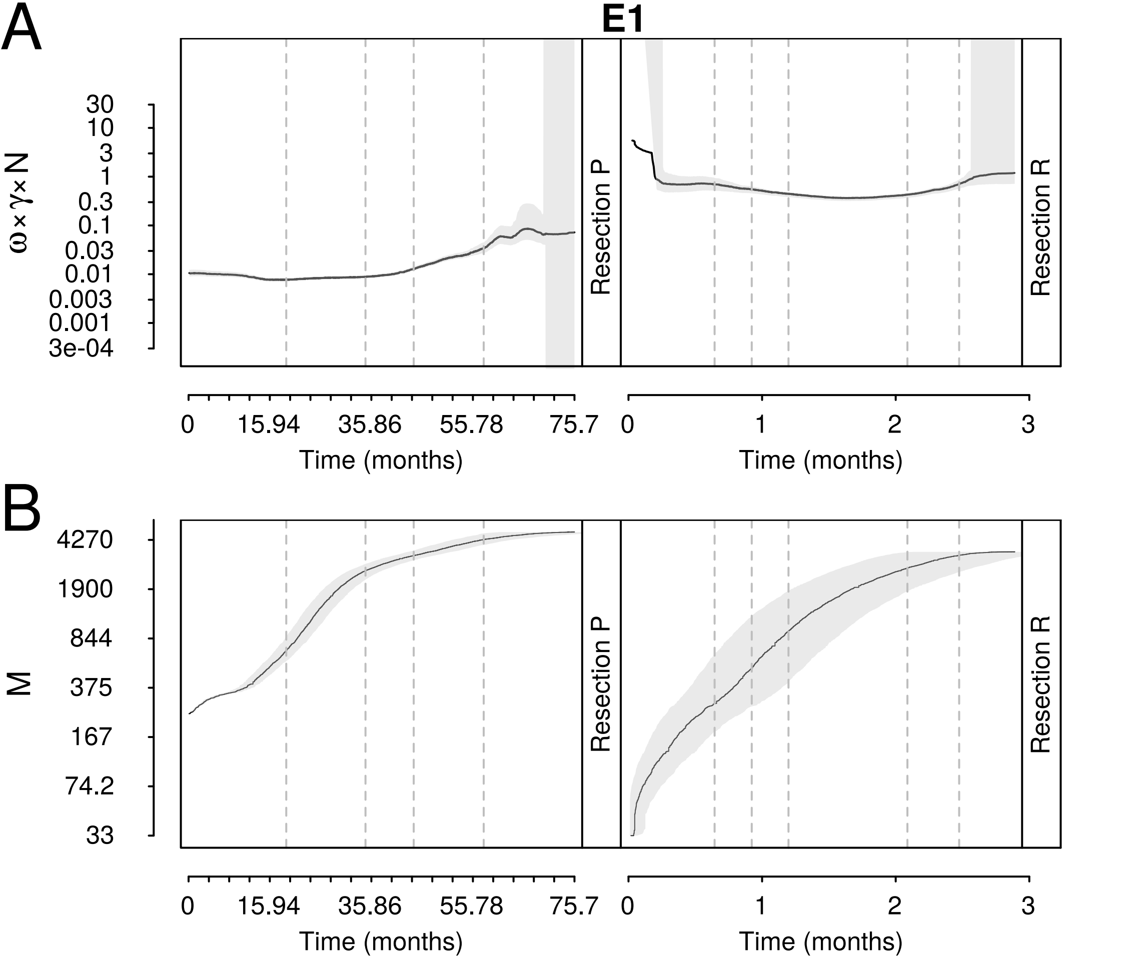
A
B
E1
ω×γ×N
30
10
3
1
0.3
0.1
0.03
0.01
0.003
0.001
3e-04
Resection P
Resection R
0
15.94
35.86
55.78
75.7
1
2
3
Time (months)
M
4270
1900
844
375
167
74.2
33

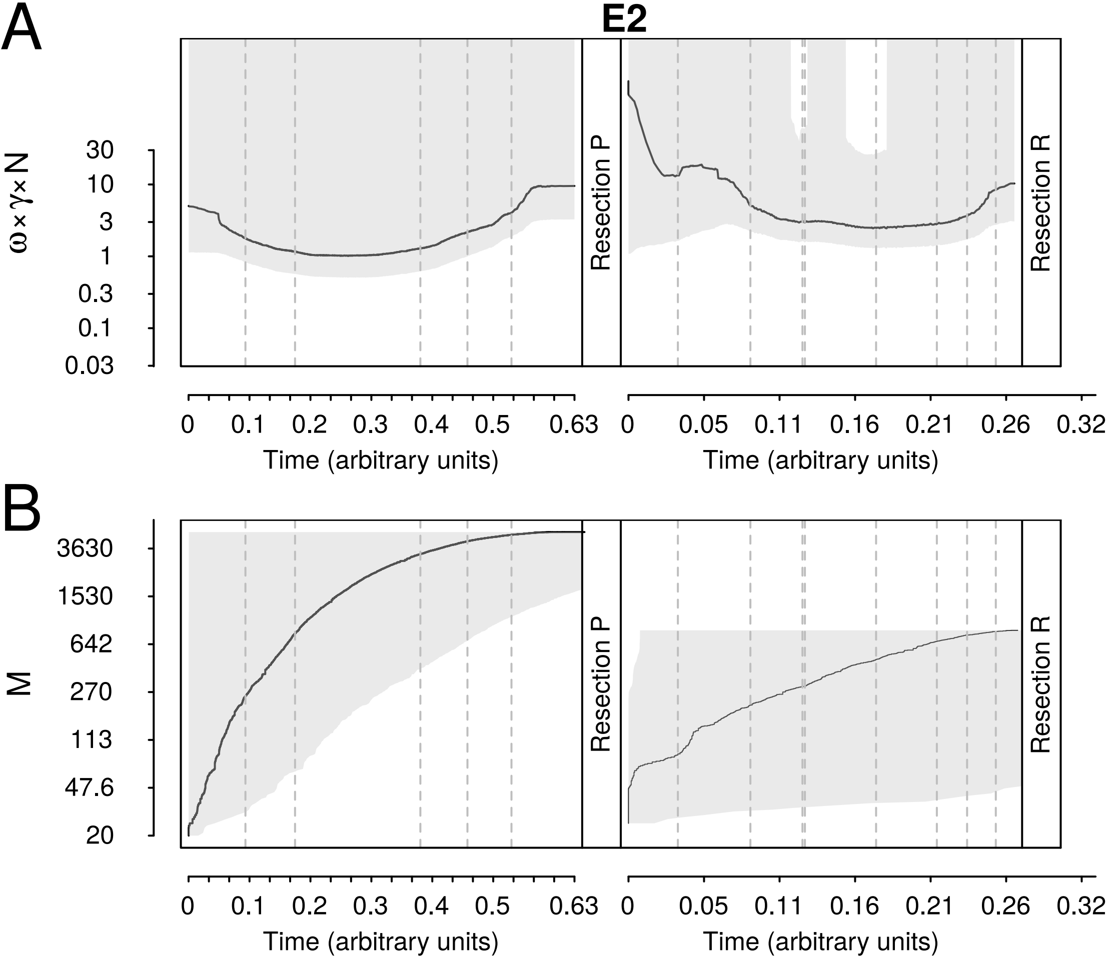
A
B
E2
ω×γ×N
30
10
3
1
0.3
0.1
0.03
M
3630
1530
642
270
113
47.6
20
Resection P
Resection R
0
0.1
0.2
0.3
0.4
0.5
0.63
0.05
0.11
0.16
0.21
0.26
0.32
Time (arbitrary units)

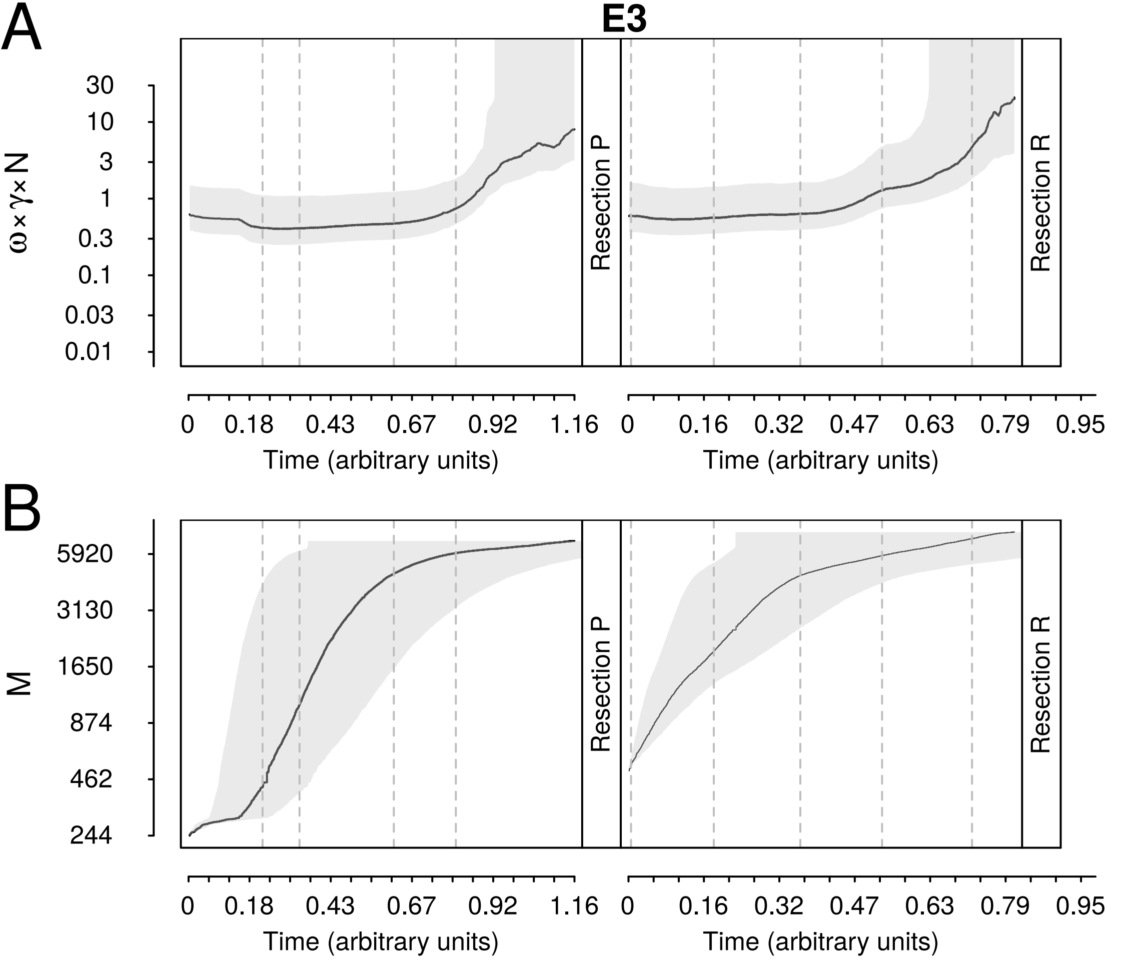
A
E3
ω×γ×N
30
10
3
1
0.3
0.1
0.03
0.01
Resection P
Resection R
0
0.18
0.43
0.67
0.92
1.16
0
0.16
0.32
0.47
0.63
0.79
0.95
Time (arbitrary units)
Time (arbitrary units)
B
M
5920
3130
1650
874
462
244
Resection P
Resection R
0
0.18
0.43
0.67
0.92
1.16
0
0.16
0.32
0.47
0.63
0.79
0.95
Time (arbitrary units)
Time (arbitrary units)

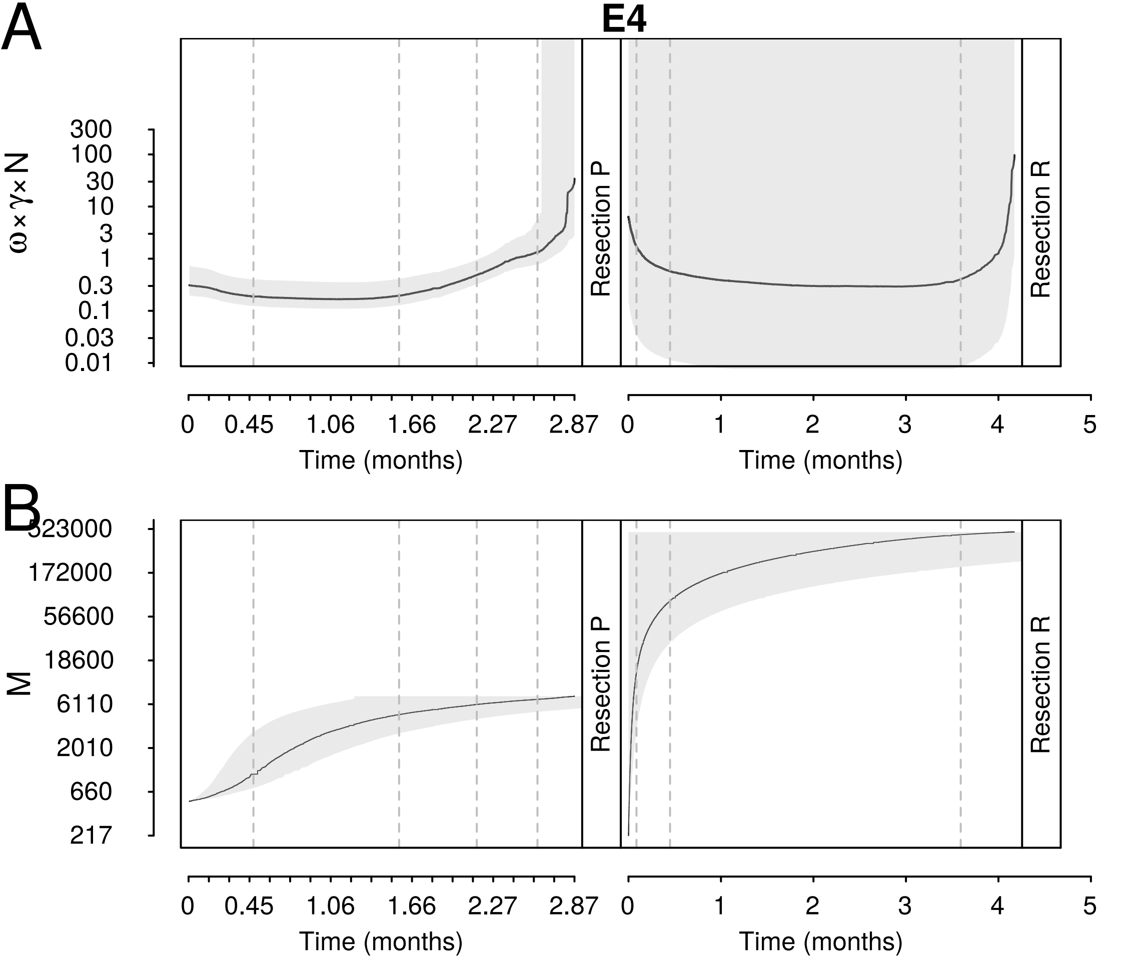
A
E4
ω×γ×N
300
100
30
10
3
1
0.3
0.1
0.03
0.01
Resection P
Resection R
0
0.45
1.06
1.66
2.27
2.87
Time (months)
0
1
2
3
4
5
Time (months)
B
323000
172000
56600
18600
6110
2010
660
217
M
Resection P
Resection R
0
0.45
1.06
1.66
2.27
2.87
Time (months)
0
1
2
3
4
5
Time (months)

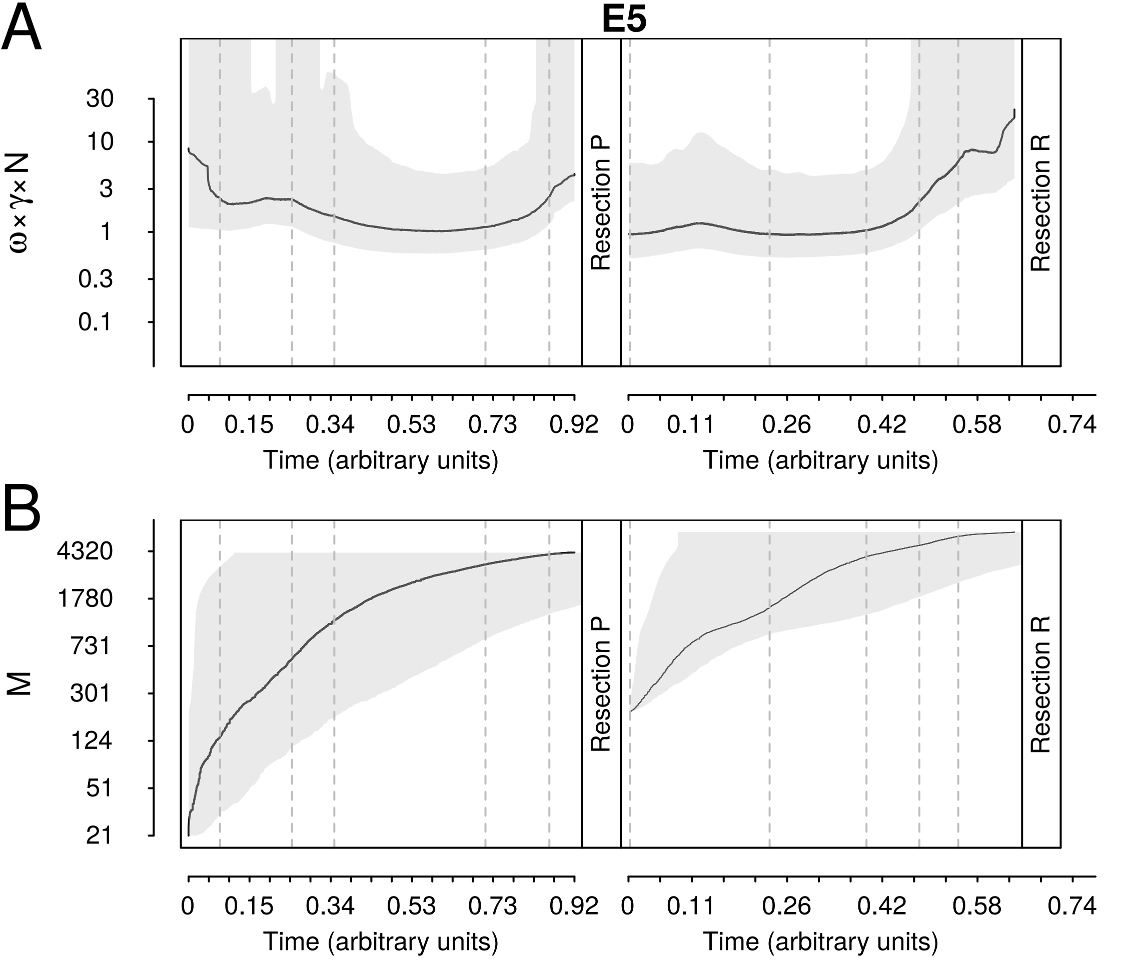
A
E5
30
10
3
1
0.3
0.1
ω×γ×N
Resection P
Resection R
0
0.15
0.34
0.53
0.73
0.92
Time (arbitrary units)
0
0.11
0.26
0.42
0.58
0.74
Time (arbitrary units)
B
4320
1780
731
301
124
51
21
M
Resection P
Resection R
0
0.15
0.34
0.53
0.73
0.92
Time (arbitrary units)
0
0.11
0.26
0.42
0.58
0.74
Time (arbitrary units)

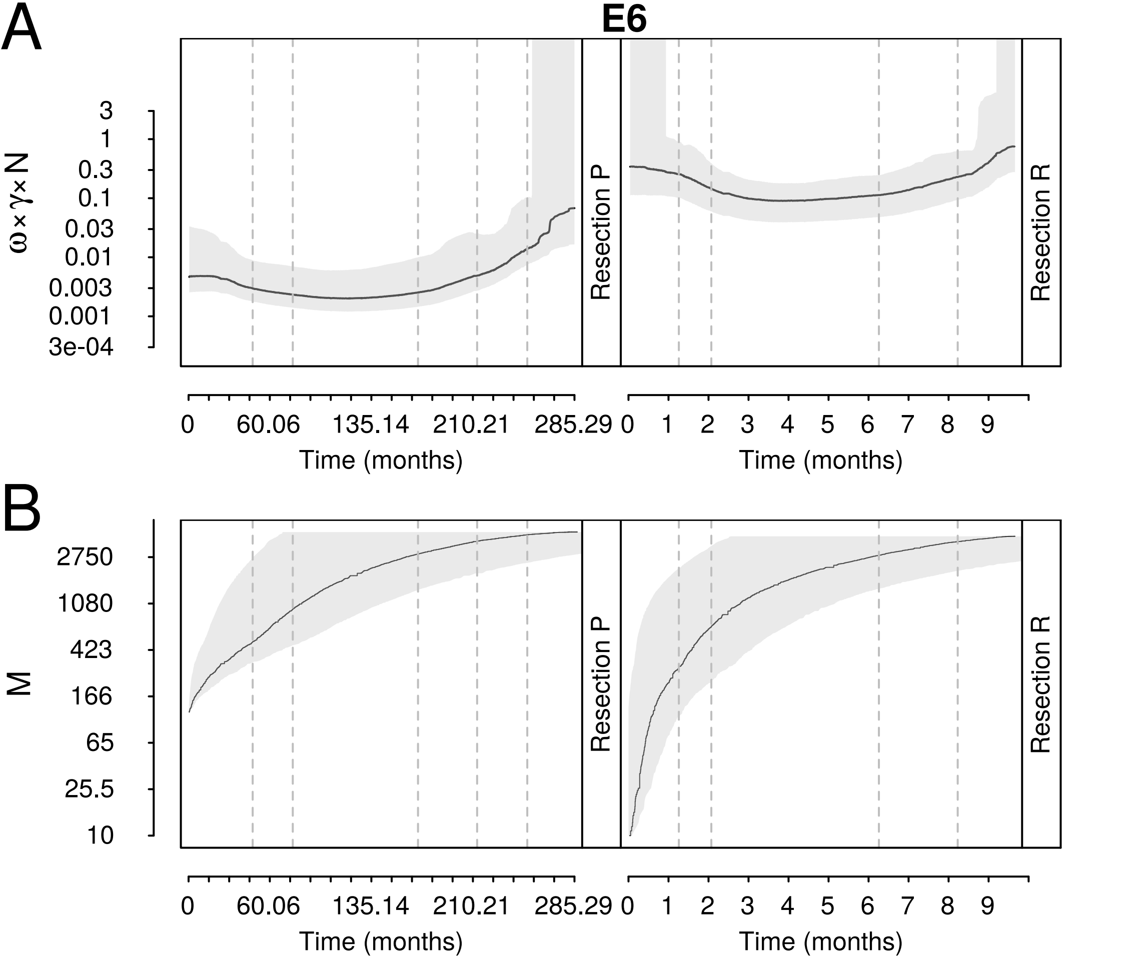

A
E6
ω×γ×N
3
1
0.3
0.1
0.03
0.01
0.003
0.001
3e-04
Resection P
Resection R
0
60.06
135.14
210.21
285.29
0 1 2 3 4 5 6 7 8 9
Time (months)
B
M
2750
1080
423
166
65
25.5
10

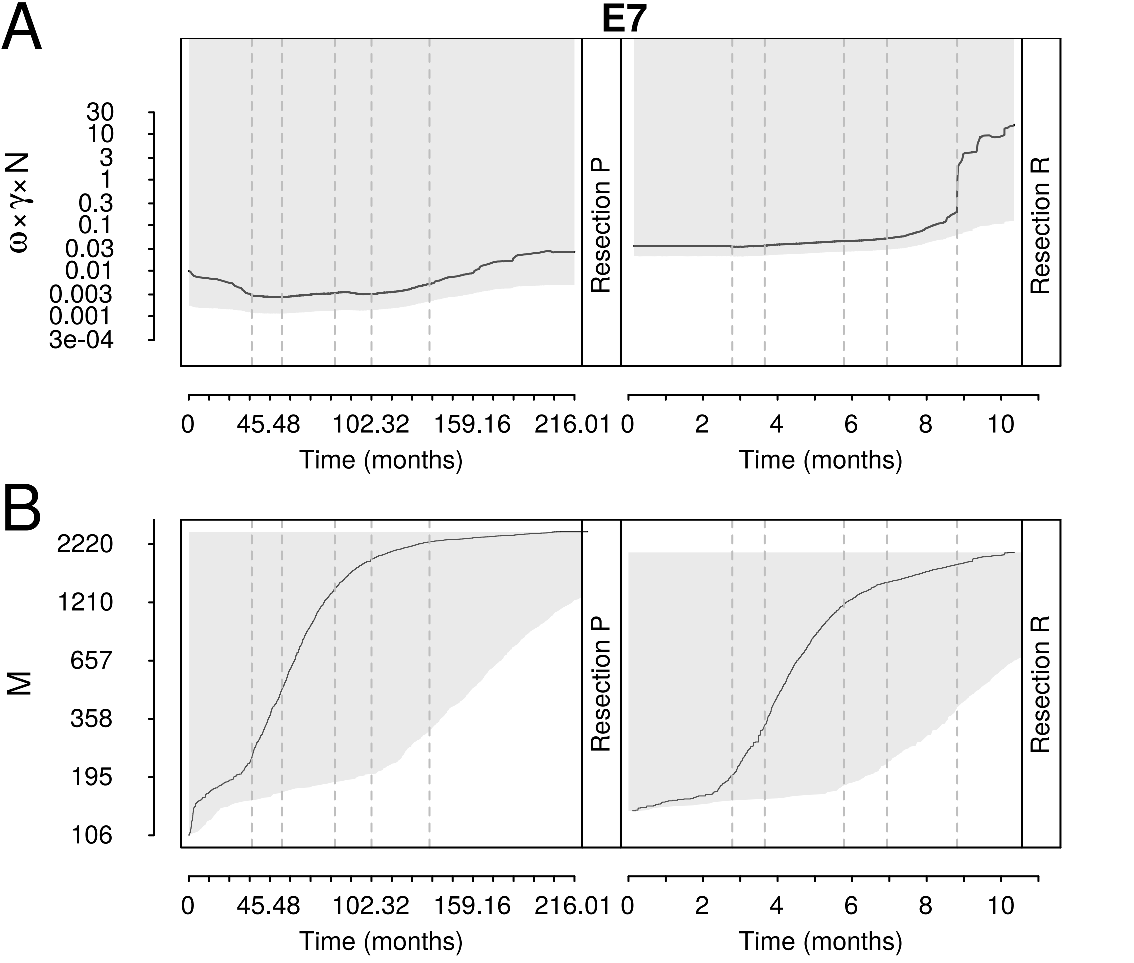
A
E7
ω×γ×N
30
10
3
1
0.3
0.1
0.03
0.01
0.003
0.001
3e-04
Resection P
Resection R
0
45.48
102.32
159.16
216.01
0
2
4
6
8
10
Time (months)
Time (months)
B
M
2220
1210
657
358
195
106
Resection P
Resection R
0
45.48
102.32
159.16
216.01
0
2
4
6
8
10
Time (months)
Time (months)

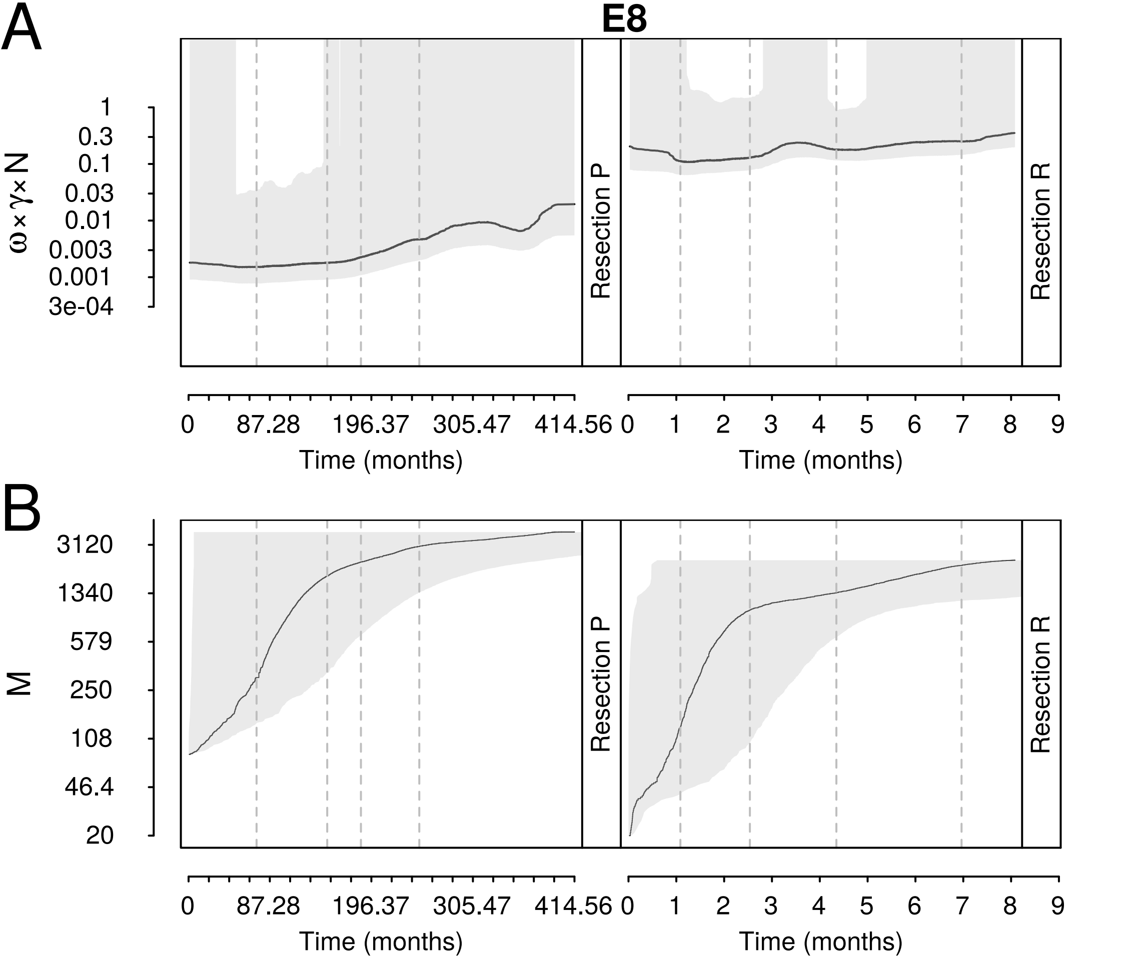
A
E8
B
ω×γ×N
1
0.3
0.1
0.03
0.01
0.003
0.001
3e-04
M
3120
1340
579
250
108
46.4
20
Resection P
Resection R
0
87.28
196.37
305.47
414.56
0
1
2
3
4
5
6
7
8
9
Time (months)

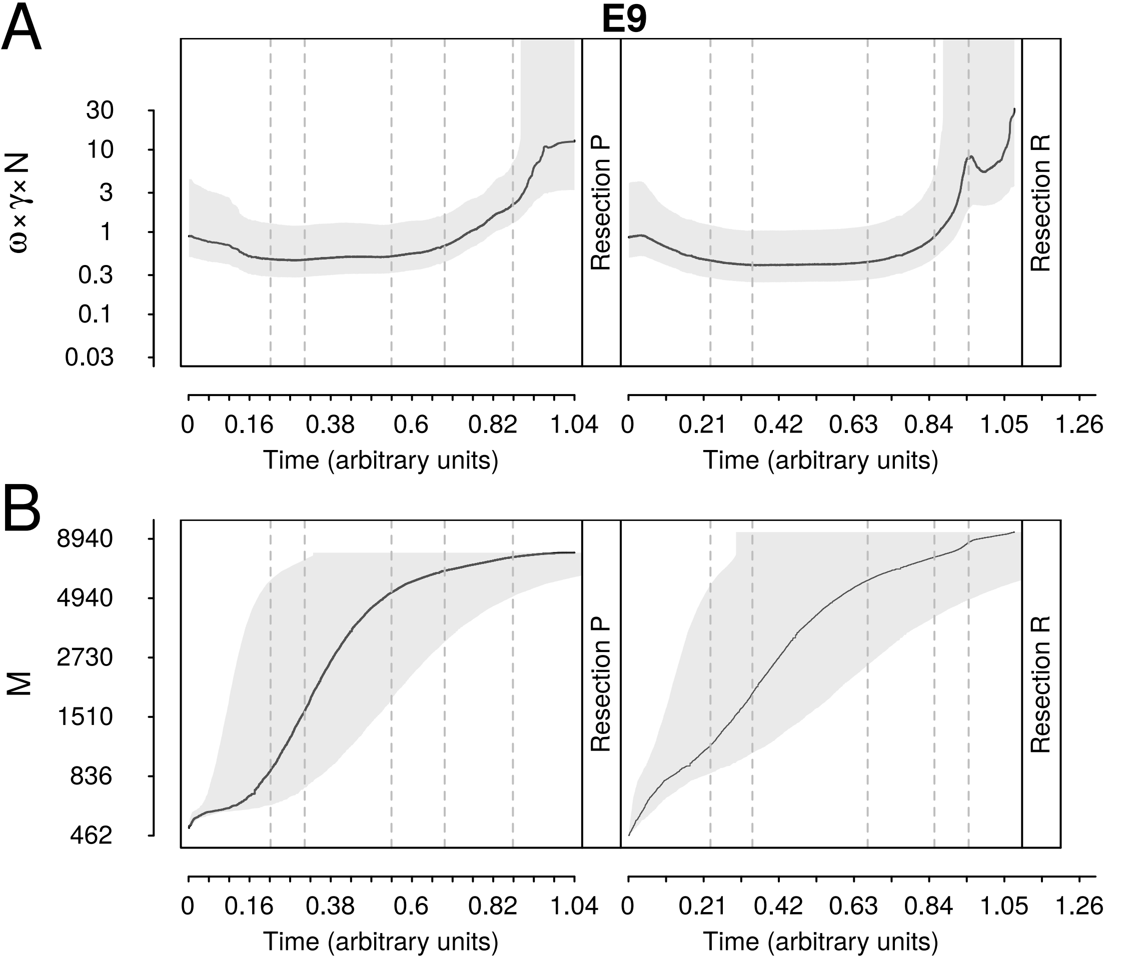
A
B
E9
ω×γ×N
30
10
3
1
0.3
0.1
0.03
Resection P
Resection R
0
0.16
0.38
0.6
0.82
1.04
0.21
0.42
0.63
0.84
1.05
1.26
Time (arbitrary units)
M
8940
4940
2730
1510
836
462

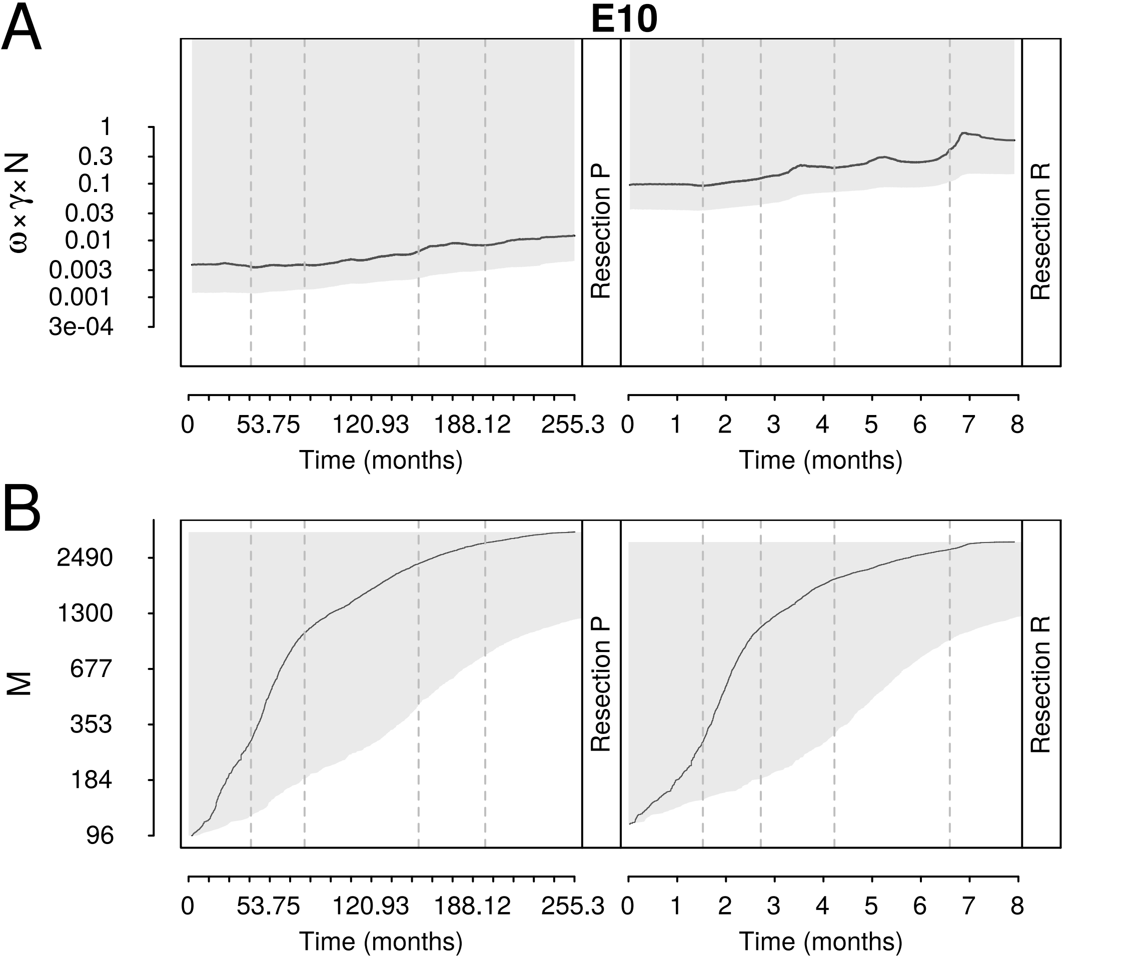

A
B
E10
ω×γ×N
1
0.3
0.1
0.03
0.01
0.003
0.001
3e-04
Resection P
Resection R
M
2490
1300
677
353
184
96
0
53.75
120.93
188.12
255.3
1
2
3
4
5
6
7
8
Time (months)

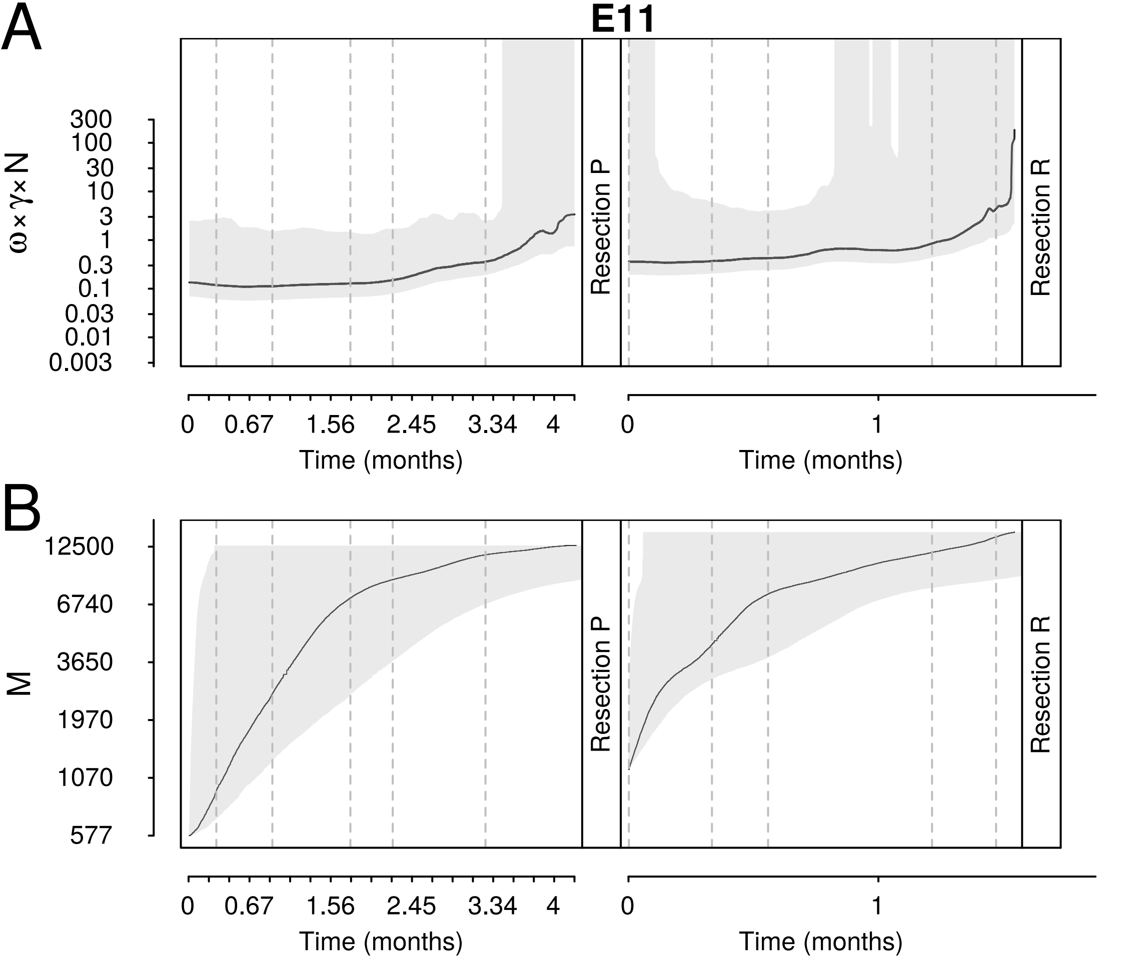

A
E11
300
100
30
10
3
1
0.3
0.1
0.03
0.01
0.003
ω×γ×N
Resection P
Resection R
0
0.67
1.56
2.45
3.34
4
Time (months)
0
1
Time (months)
B
12500
6740
3650
1970
1070
577
M
Resection P
Resection R
0
0.67
1.56
2.45
3.34
4
Time (months)
0
1
Time (months)

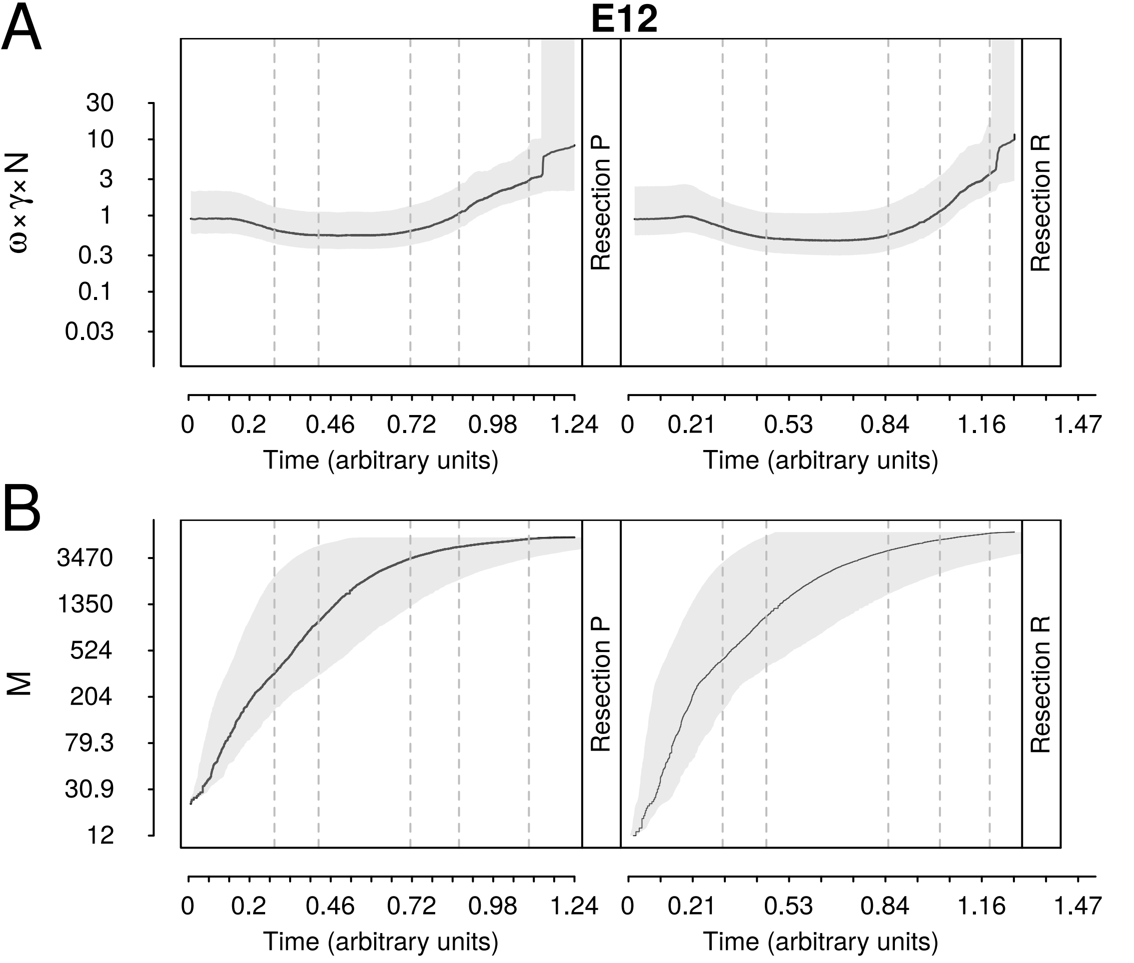
A
B
E12
ω×γ×N
30
10
3
1
0.3
0.1
0.03
Resection P
Resection R
0
0.2
0.46
0.72
0.98
1.24
0.21
0.53
0.84
1.16
1.47
Time (arbitrary units)
M
3470
1350
524
204
79.3
30.9
12

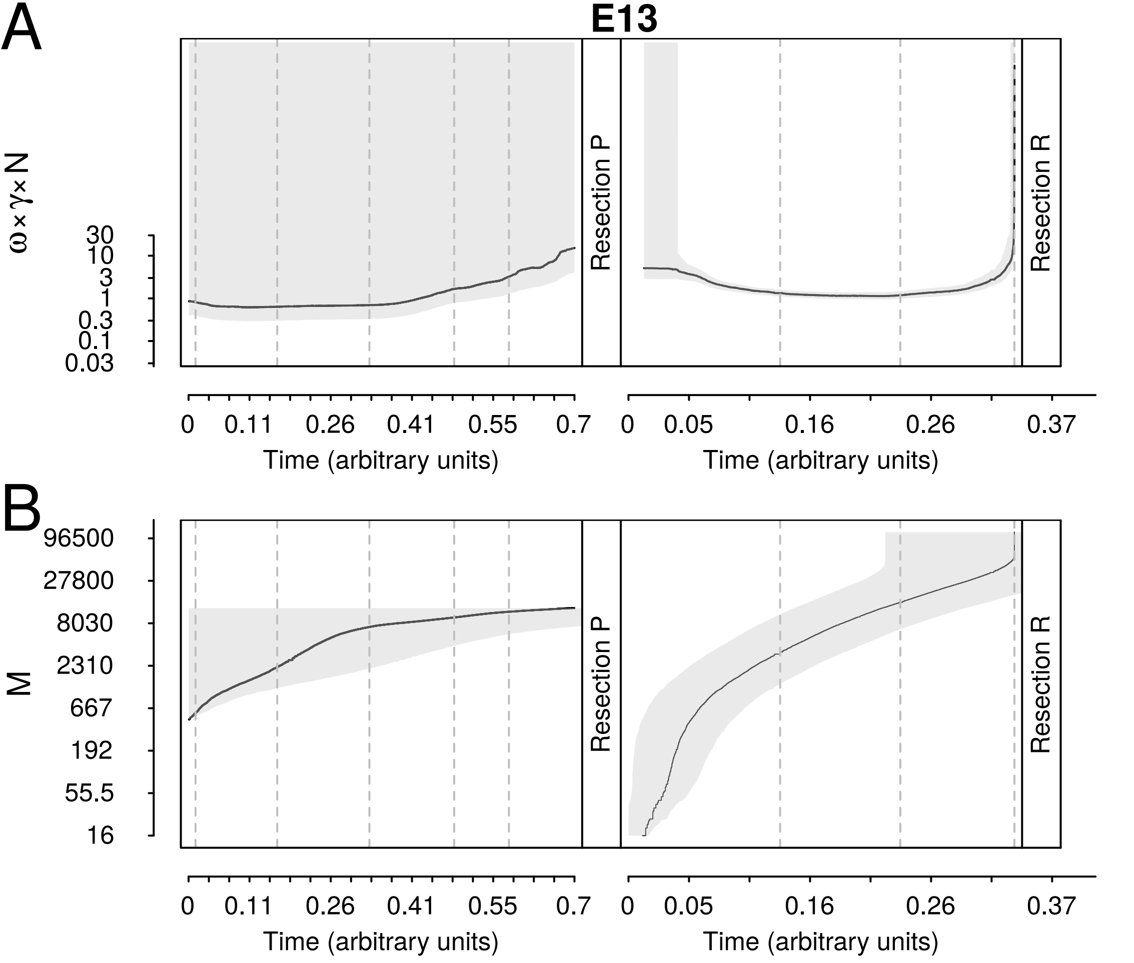

A
E13
ω×γ×N
30
10
3
1
0.3
0.1
0.03
Resection P
Resection R
0
0.11
0.26
0.41
0.55
0.7
Time (arbitrary units)
0
0.05
0.16
0.26
0.37
Time (arbitrary units)
B
96500
27800
8030
2310
667
192
55.5
16
M
Resection P
Resection R
0
0.11
0.26
0.41
0.55
0.7
Time (arbitrary units)
0
0.05
0.16
0.26
0.37
Time (arbitrary units)

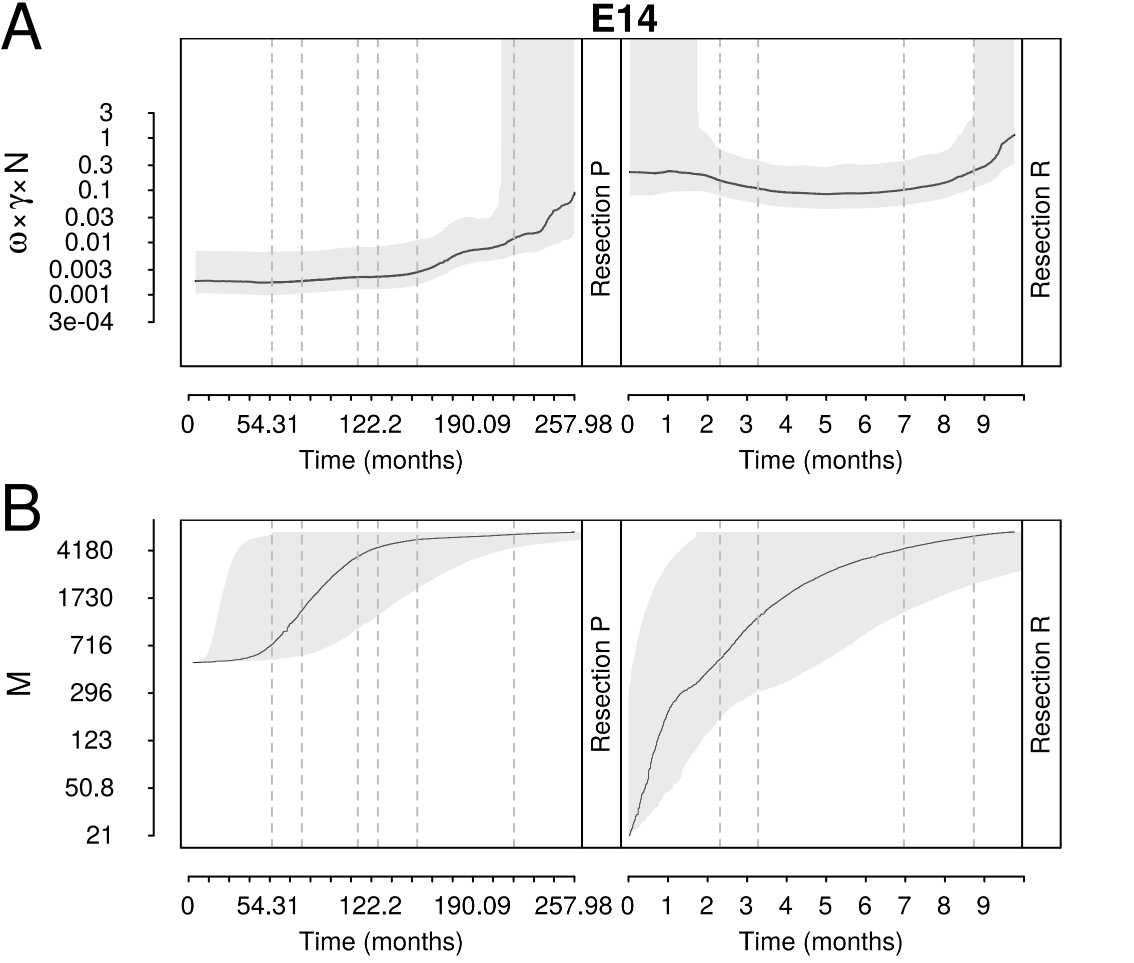
A
E14
B
ω×γ×N
3
1
0.3
0.1
0.03
0.01
0.003
0.001
3e-04
M
4180
1730
716
296
123
50.8
21
Resection P
Resection R
0
54.31
122.2
190.09
257.98
0 1 2 3 4 5 6 7 8 9
Time (months)

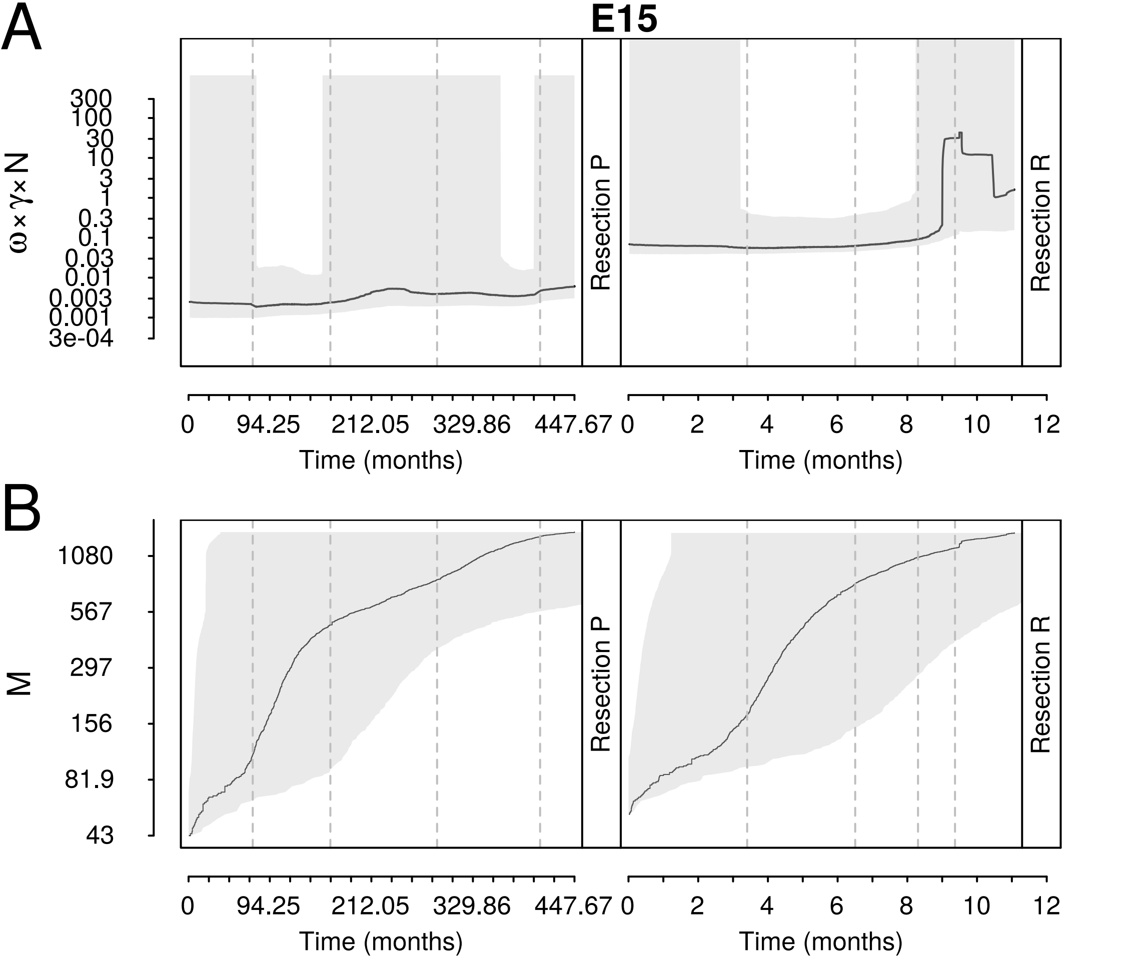
A
E15
ω×γ×N
300
100
30
10
3
1
0.3
0.1
0.03
0.01
0.003
0.001
3e-04
Resection P
Resection R
0
94.25
212.05
329.86
447.67
Time (months)
0
2
4
6
8
10
12
Time (months)
B
M
1080
567
297
156
81.9
43
Resection P
Resection R
0
94.25
212.05
329.86
447.67
Time (months)
0
2
4
6
8
10
12
Time (months)

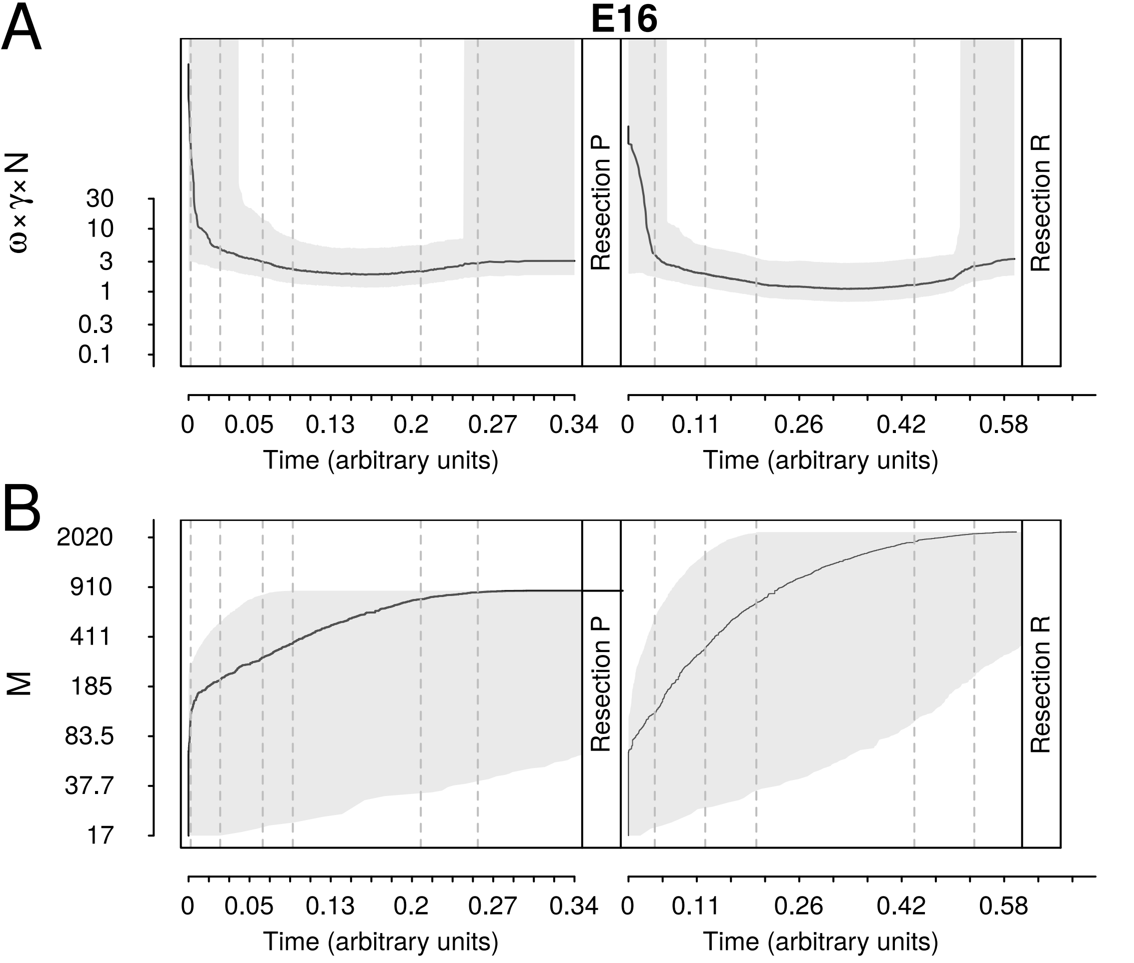

A
E16
ω×γ×N
30
10
3
1
0.3
0.1
Resection P
Resection R
0
0.05
0.13
0.2
0.27
0.34
0
0.11
0.26
0.42
0.58
Time (arbitrary units)
Time (arbitrary units)
B
M
2020
910
411
185
83.5
37.7
17
Resection P
Resection R
0
0.05
0.13
0.2
0.27
0.34
0
0.11
0.26
0.42
0.58
Time (arbitrary units)
Time (arbitrary units)

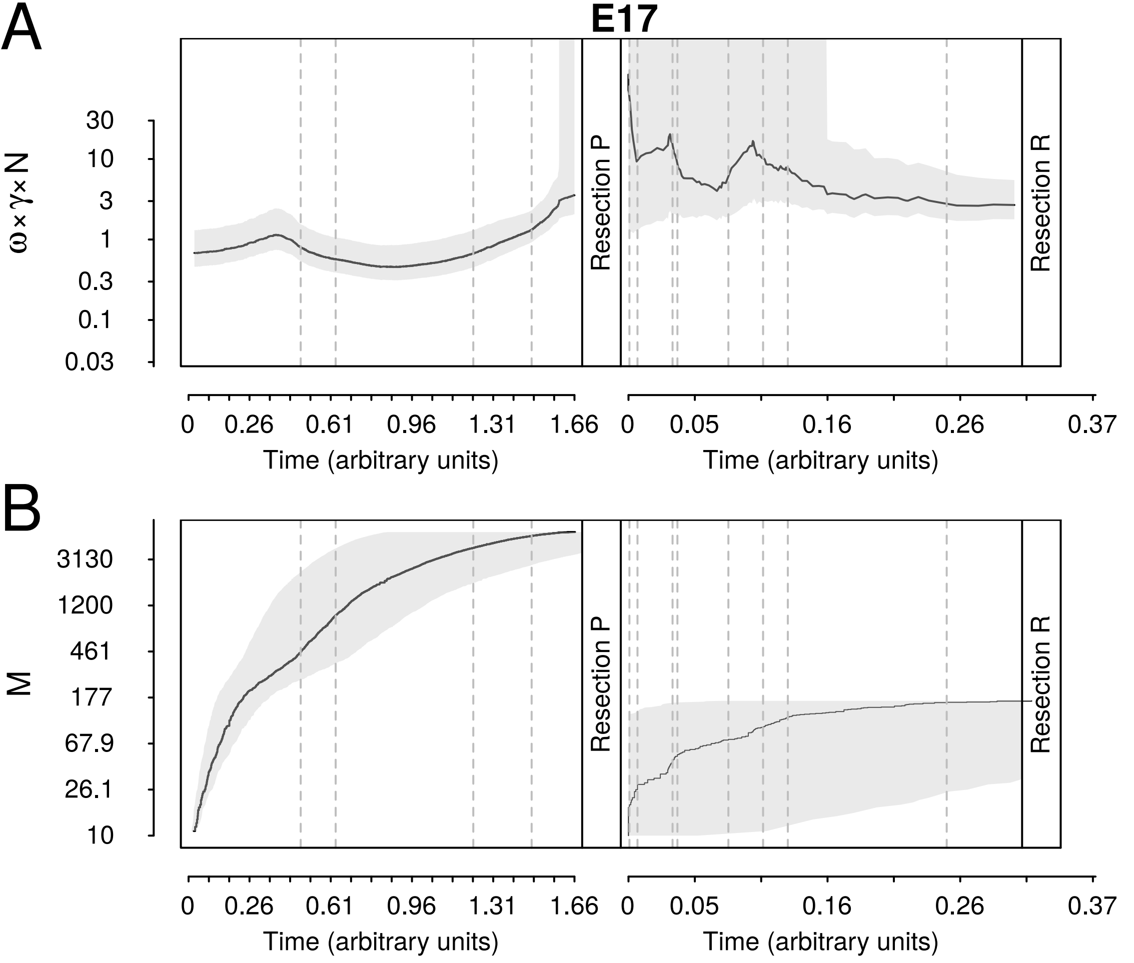
A
E17
ω×γ×N
30
10
3
1
0.3
0.1
0.03
Resection P
Resection R
0
0.26
0.61
0.96
1.31
1.66
Time (arbitrary units)
0
0.05
0.16
0.26
0.37
Time (arbitrary units)
B
M
3130
1200
461
177
67.9
26.1
10
Resection P
Resection R
0
0.26
0.61
0.96
1.31
1.66
Time (arbitrary units)
0
0.05
0.16
0.26
0.37
Time (arbitrary units)

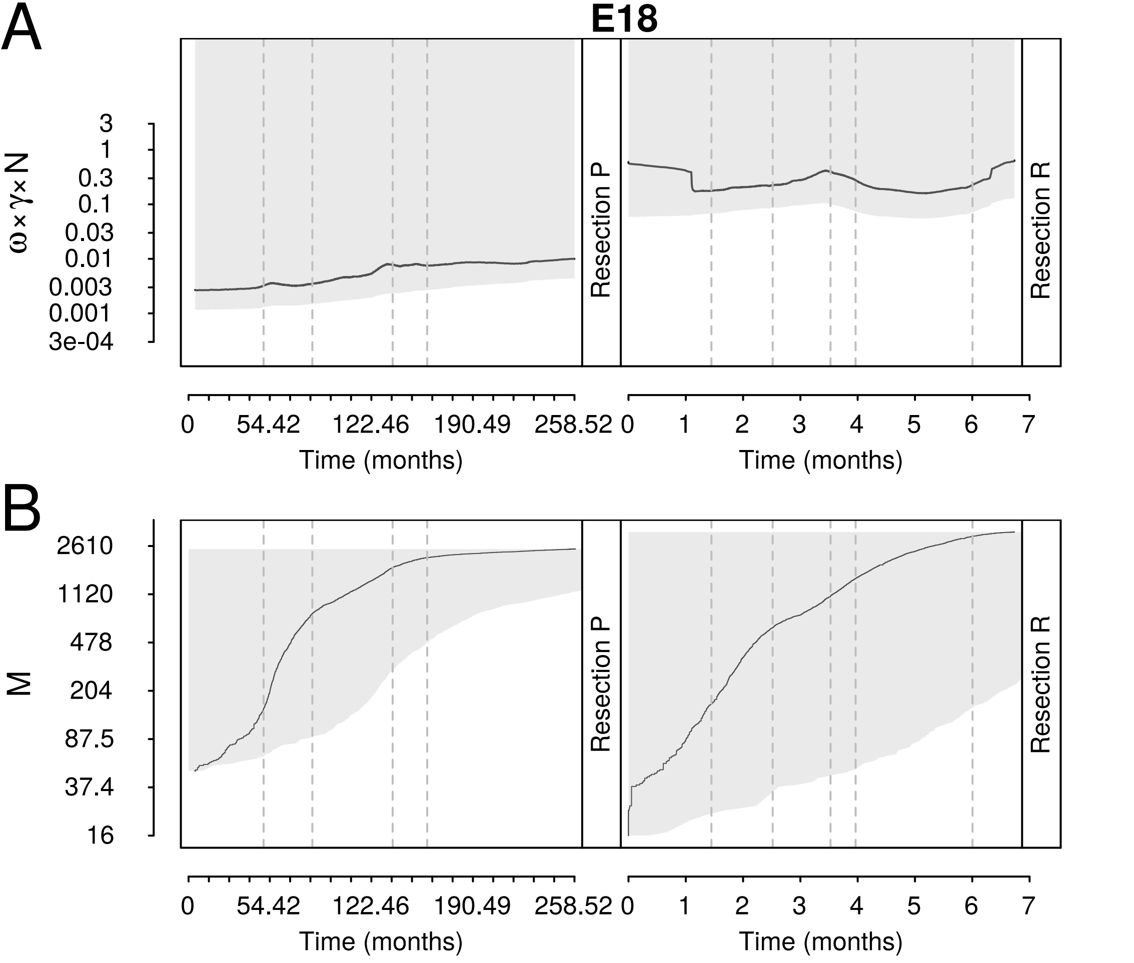
A
B
E18
ω×γ×N
3
1
0.3
0.1
0.03
0.01
0.003
0.001
3e-04
M
2610
1120
478
204
87.5
37.4
16
Resection P
Resection R
0
54.42
122.46
190.49
258.52
1
2
3
4
5
6
7
Time (months)

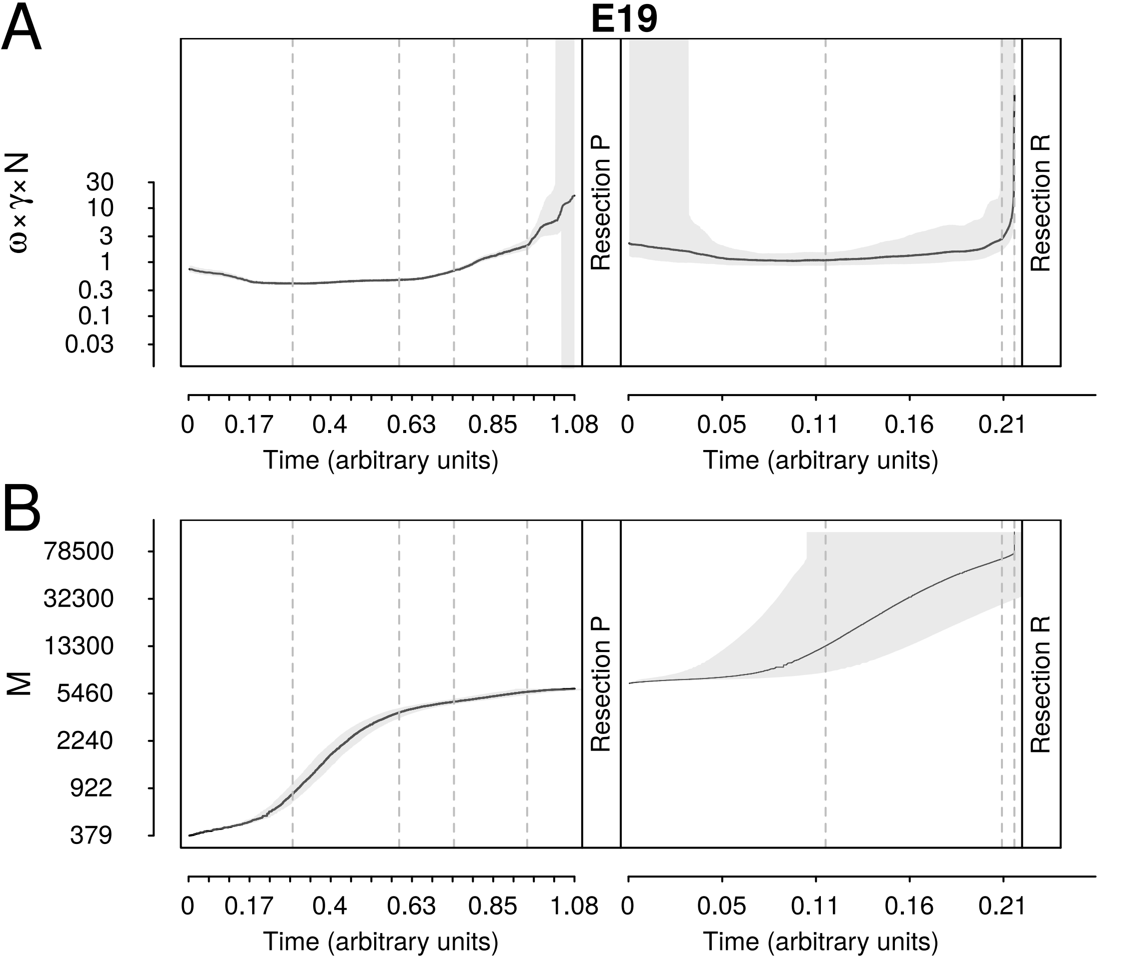
A
B
E19
ω×γ×N
30
10
3
1
0.3
0.1
0.03
M
78500
32300
13300
5460
2240
922
379
Resection P
Resection R
0
0.17
0.4
0.63
0.85
1.08
0
0.05
0.11
0.16
0.21
Time (arbitrary units)

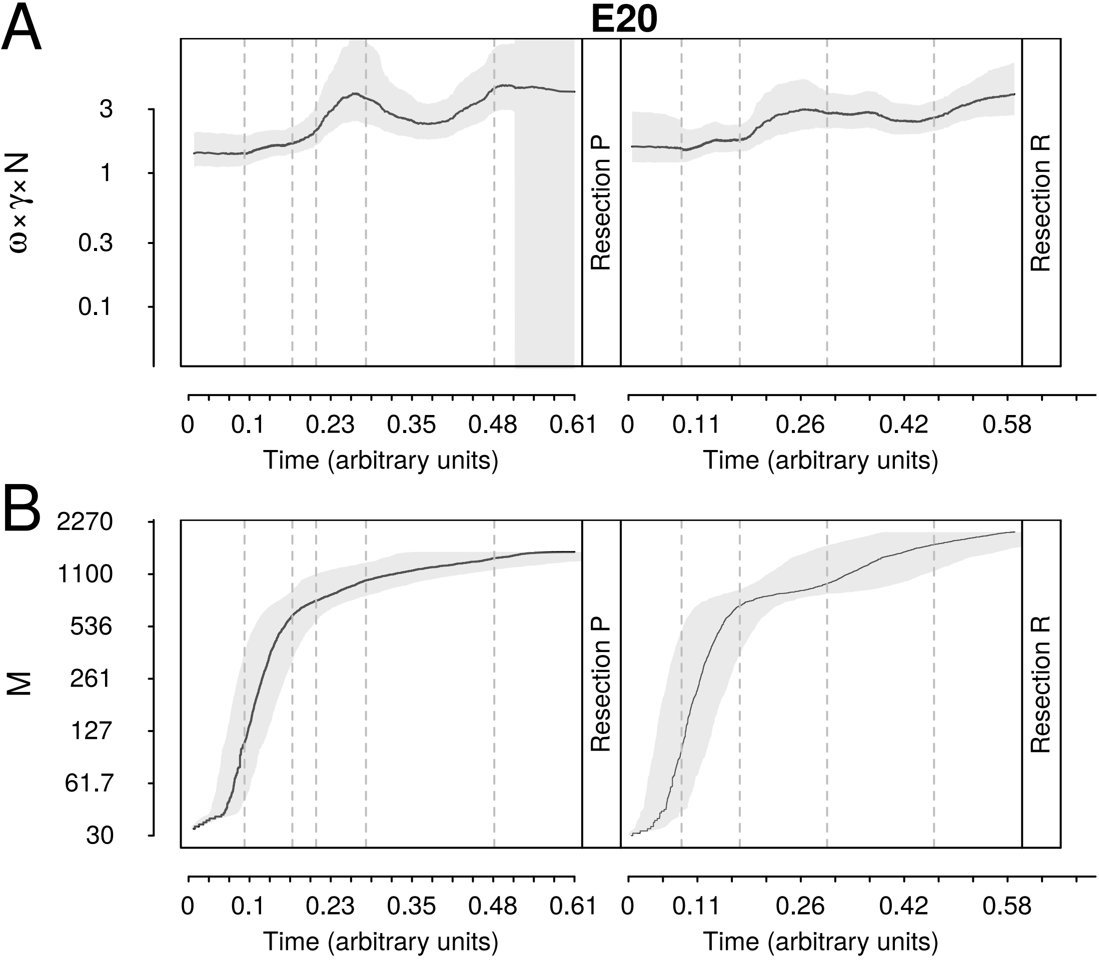
A
B
E20
ω×γ×N
3
1
0.3
0.1
M
2270
1100
536
261
127
61.7
30
Resection P
Resection R
0
0.1
0.23
0.35
0.48
0.61
0.11
0.26
0.42
0.58
Time (arbitrary units)

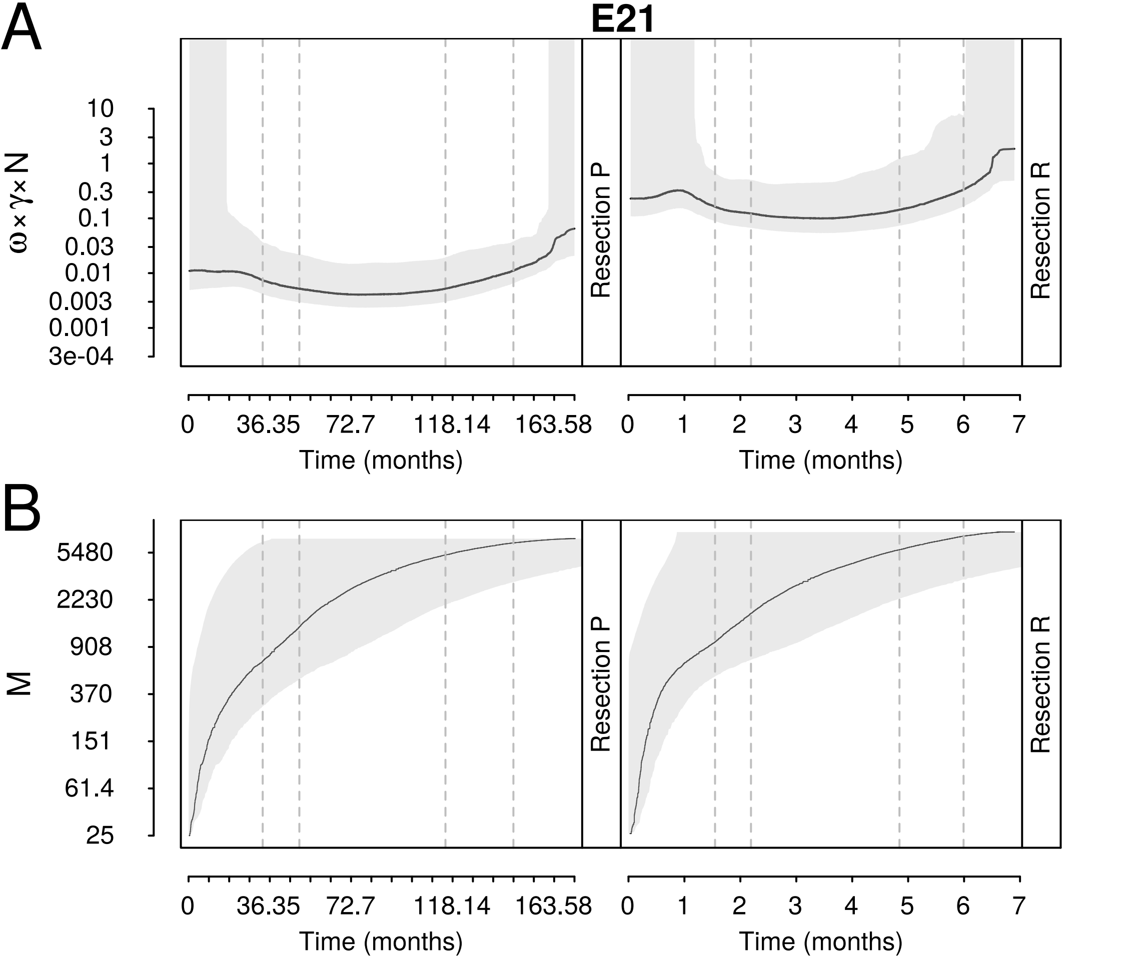
A
E21
ω×γ×N
10
3
1
0.3
0.1
0.03
0.01
0.003
0.001
3e-04
Resection P
Resection R
0
36.35
72.7
118.14
163.58
Time (months)
0
1
2
3
4
5
6
7
Time (months)
B
M
5480
2230
908
370
151
61.4
25
Resection P
Resection R
0
36.35
72.7
118.14
163.58
Time (months)
0
1
2
3
4
5
6
7
Time (months)

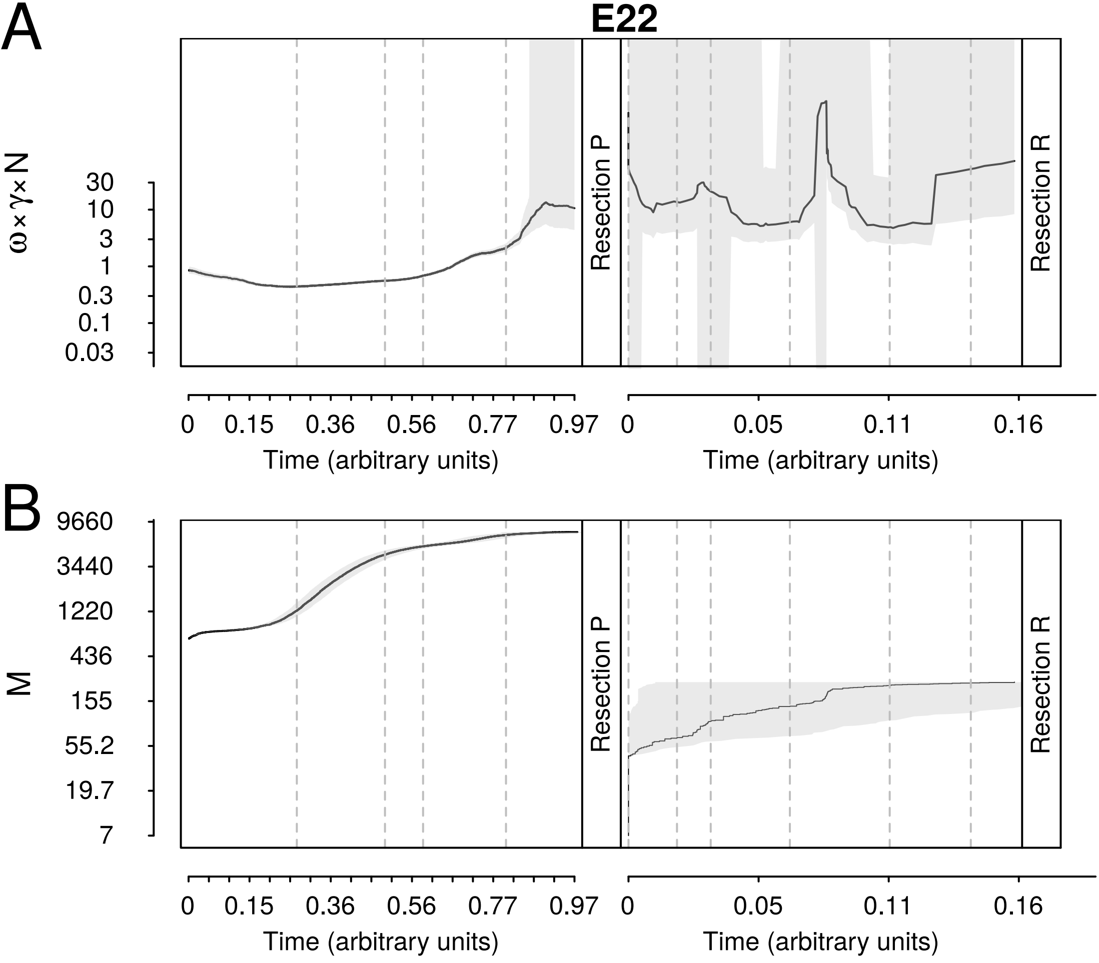

A
E22
B
ω×γ×N
30
10
3
1
0.3
0.1
0.03
Resection P
Resection R
0
0.15
0.36
0.56
0.77
0.97
0
0.05
0.11
0.16
Time (arbitrary units)
M
9660
3440
1220
436
155
55.2
19.7
7

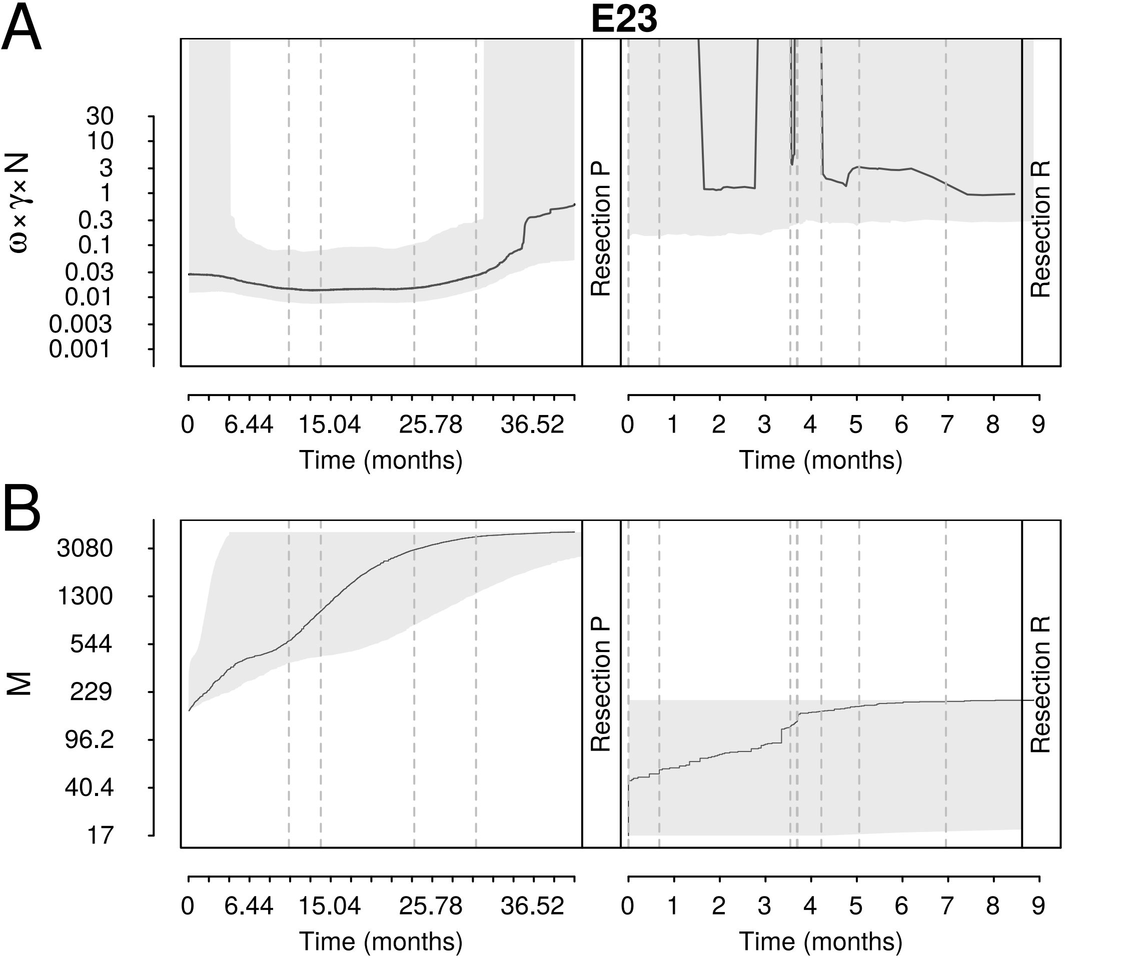
A
E23
ω×γ×N
30
10
3
1
0.3
0.1
0.03
0.01
0.003
0.001
Resection P
Resection R
0
6.44
15.04
25.78
36.52
Time (months)
0
1
2
3
4
5
6
7
8
9
Time (months)
B
M
3080
1300
544
229
96.2
40.4
17
Resection P
Resection R
0
6.44
15.04
25.78
36.52
Time (months)
0
1
2
3
4
5
6
7
8
9
Time (months)

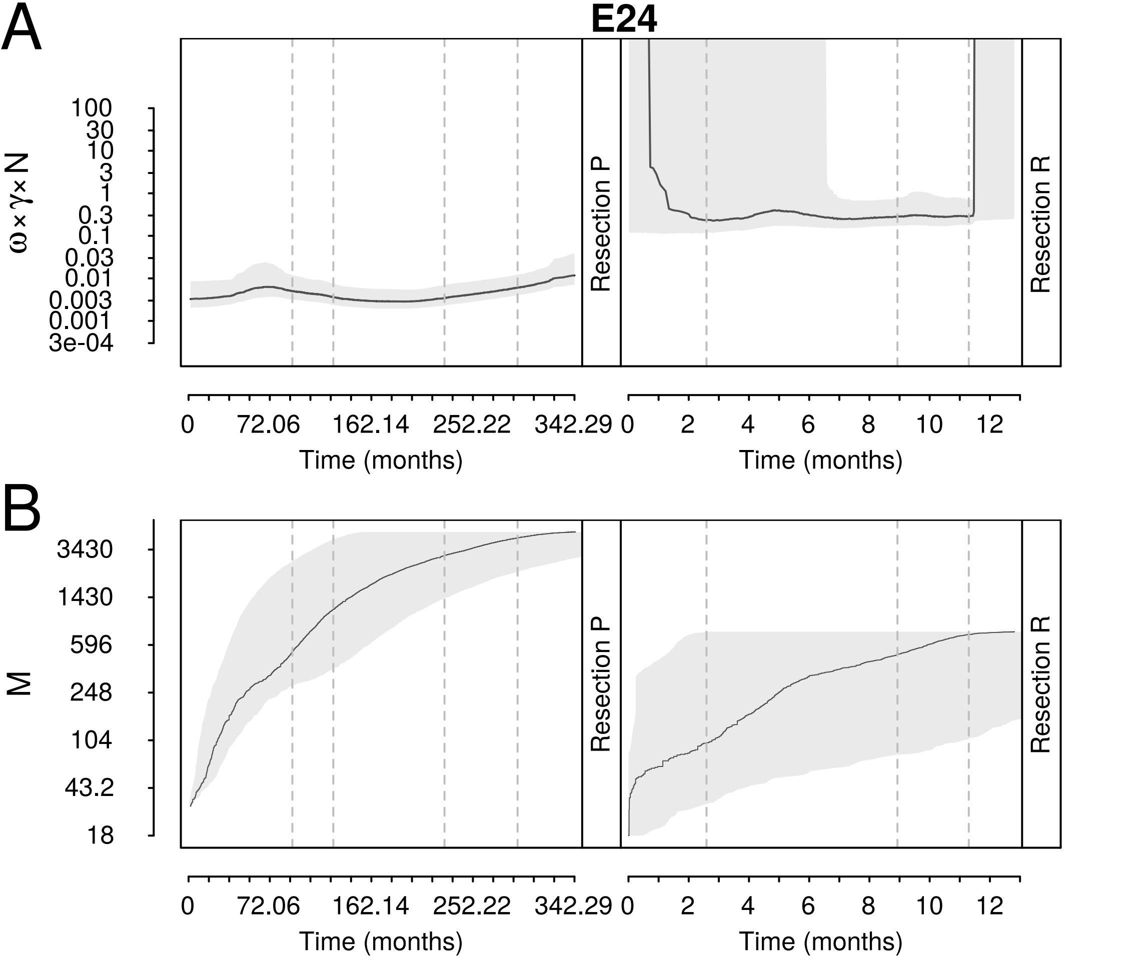

A
E24
ω×γ×N
100
30
10
3
1
0.3
0.1
0.03
0.01
0.003
0.001
3e-04
Resection P
Resection R
0
72.06
162.14
252.22
342.29
0
2
4
6
8
10
12
Time (months)
Time (months)
B
M
3430
1430
596
248
104
43.2
18
Resection P
Resection R
0
72.06
162.14
252.22
342.29
0
2
4
6
8
10
12
Time (months)
Time (months)

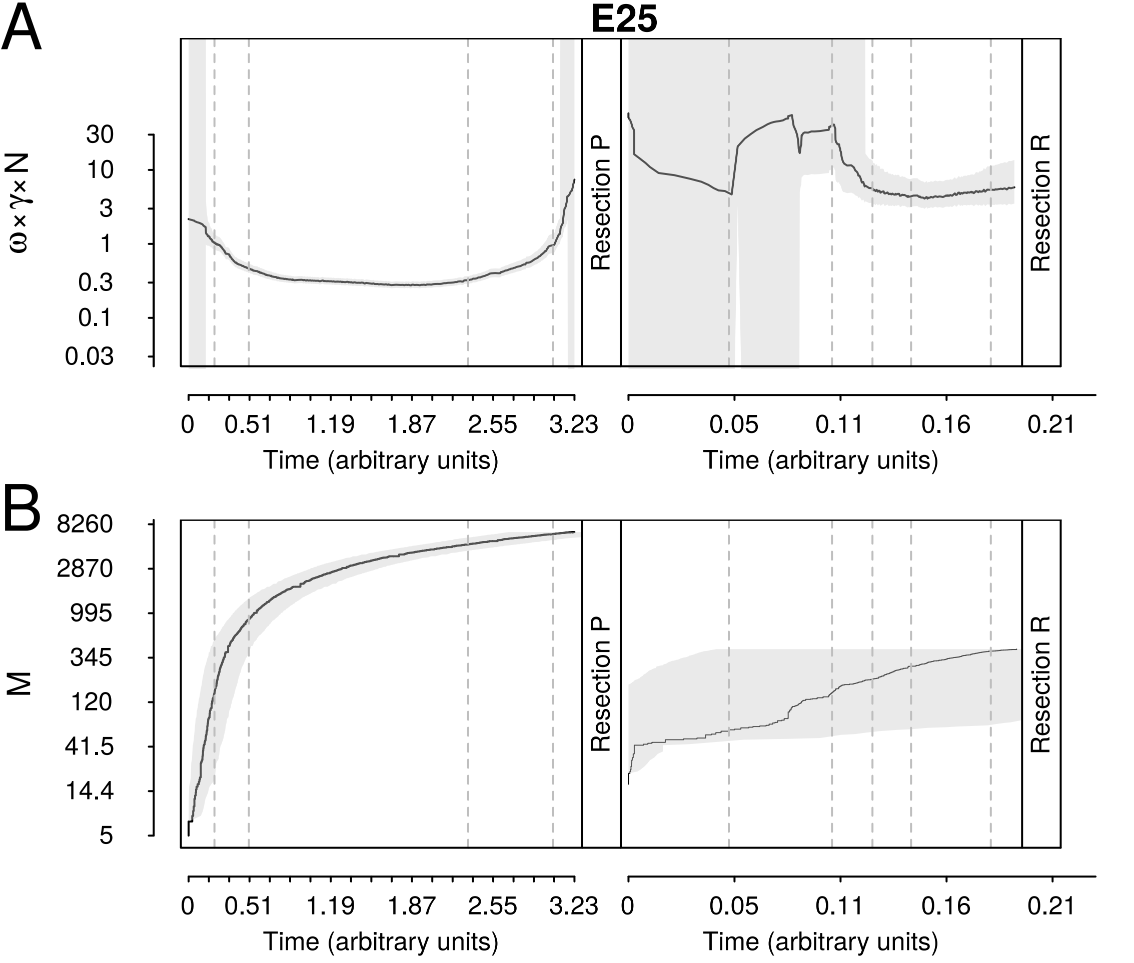
A
E25
ω×γ×N
30
10
3
1
0.3
0.1
0.03
Resection P
Resection R
0
0.51
1.19
1.87
2.55
3.23
Time (arbitrary units)
0
0.05
0.11
0.16
0.21
Time (arbitrary units)
B
M
8260
2870
995
345
120
41.5
14.4
5
Resection P
Resection R
0
0.51
1.19
1.87
2.55
3.23
Time (arbitrary units)
0
0.05
0.11
0.16
0.21
Time (arbitrary units)

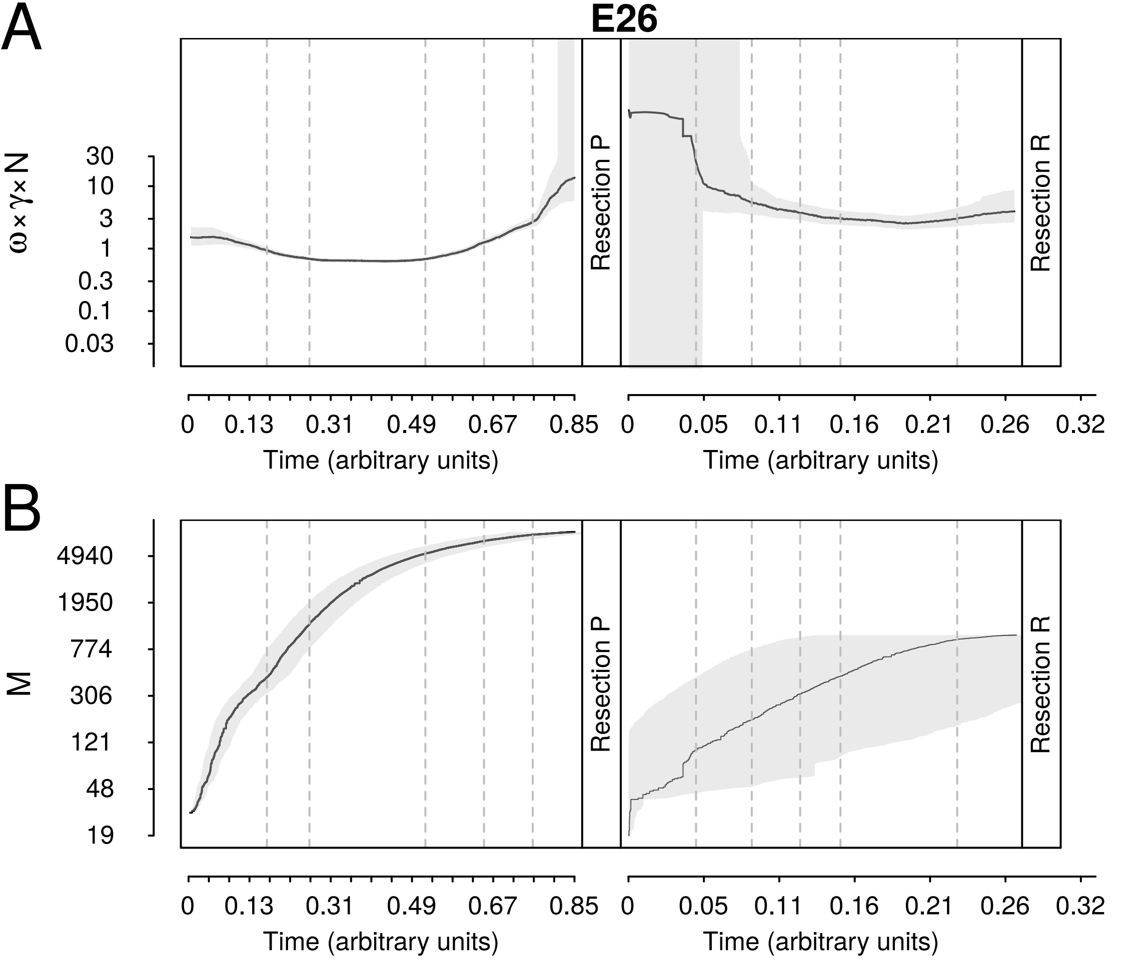

A
E26
ω×γ×N
30
10
3
1
0.3
0.1
0.03
Resection P
Resection R
0
0.13
0.31
0.49
0.67
0.85
Time (arbitrary units)
0
0.05
0.11
0.16
0.21
0.26
0.32
Time (arbitrary units)
B
M
4940
1950
774
306
121
48
19
Resection P
Resection R
0
0.13
0.31
0.49
0.67
0.85
Time (arbitrary units)
0
0.05
0.11
0.16
0.21
0.26
0.32
Time (arbitrary units)

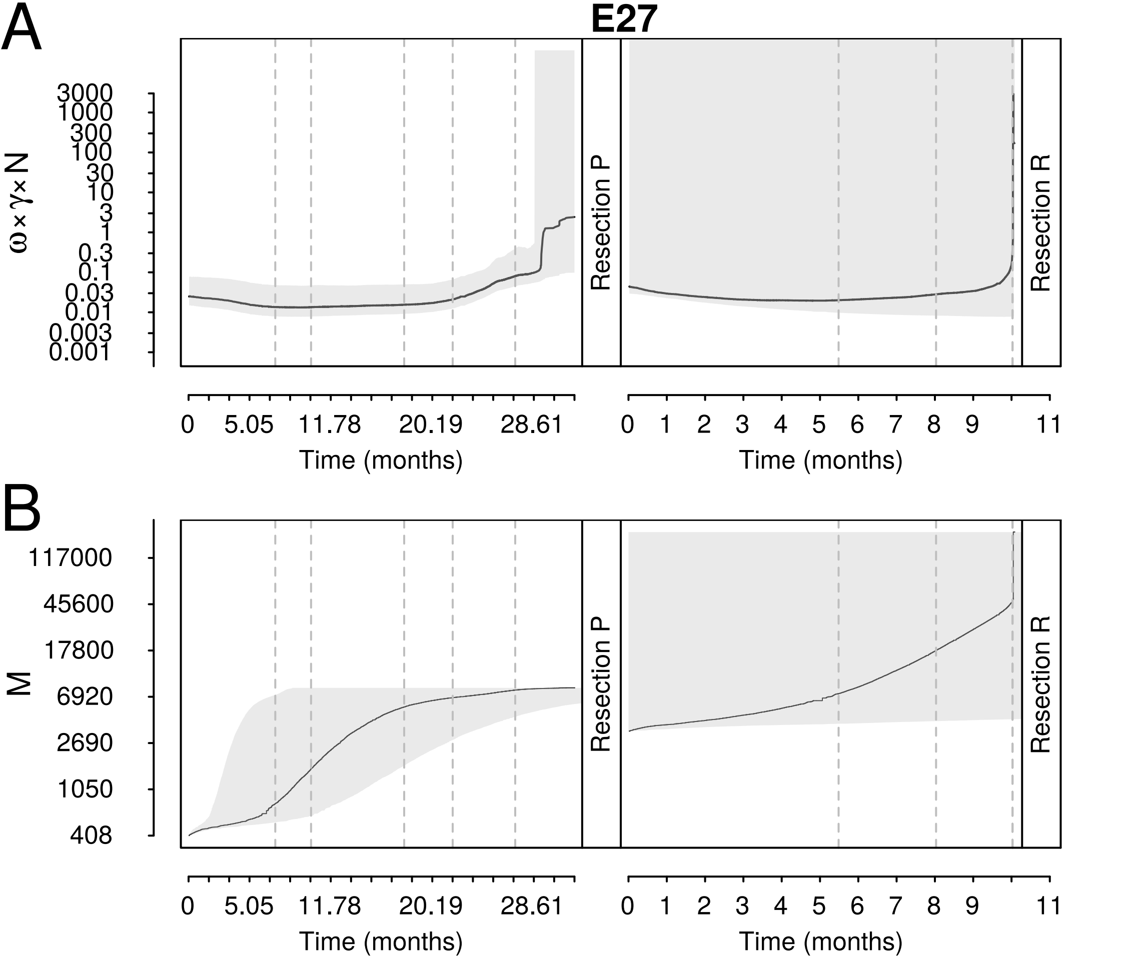

A
E27
3000
1000
300
100
30
10
3
1
0.3
0.1
0.03
0.01
0.003
0.001
ω×γ×N
Resection P
Resection R
0
5.05
11.78
20.19
28.61
Time (months)
0
1
2
3
4
5
6
7
8
9
11
Time (months)
B
117000
45600
17800
6920
2690
1050
408
M
Resection P
Resection R
0
5.05
11.78
20.19
28.61
Time (months)
0
1
2
3
4
5
6
7
8
9
11
Time (months)

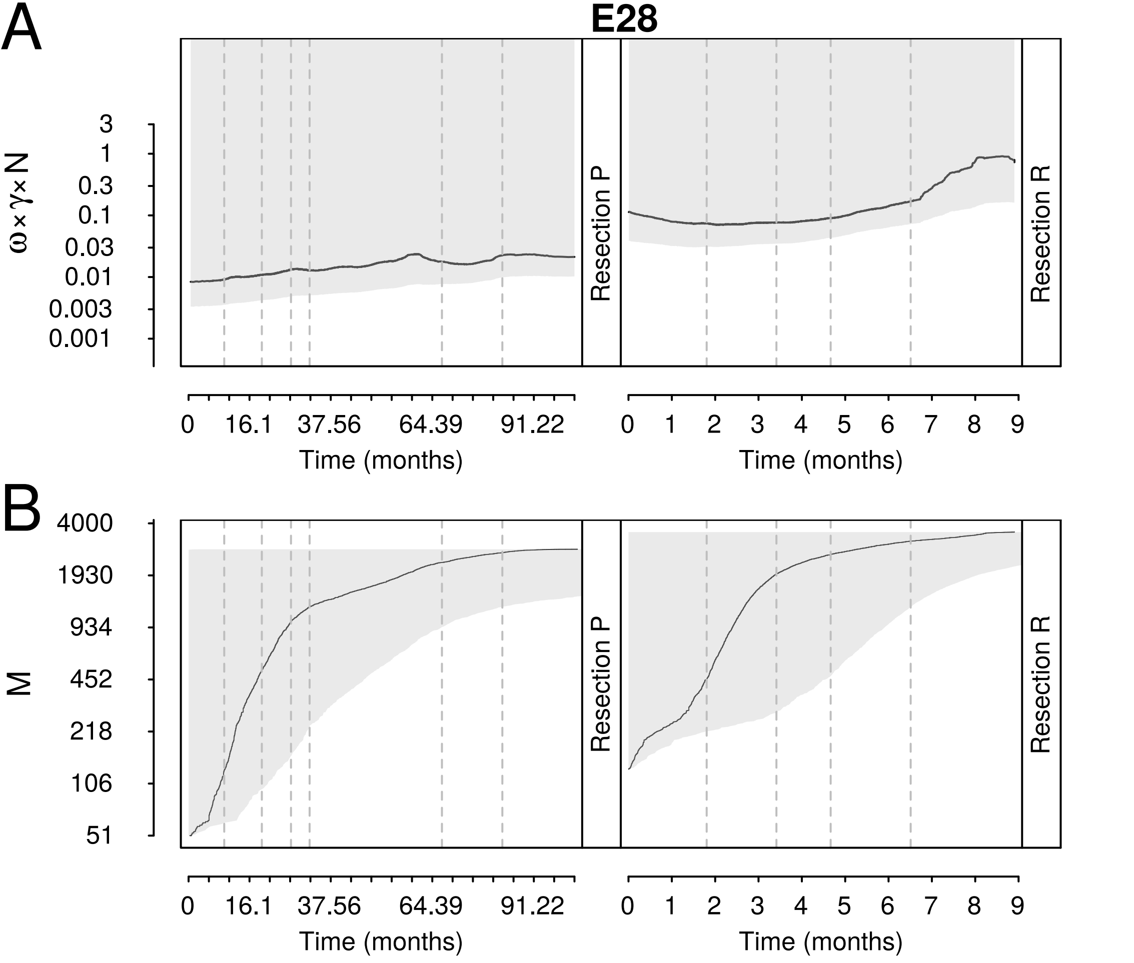
A
B
E28
ω×γ×N
3
1
0.3
0.1
0.03
0.01
0.003
0.001
Resection P
Resection R
0
16.1
37.56
64.39
91.22
Time (months)
0
1
2
3
4
5
6
7
8
9
Time (months)
M
4000
1930
934
452
218
106
51

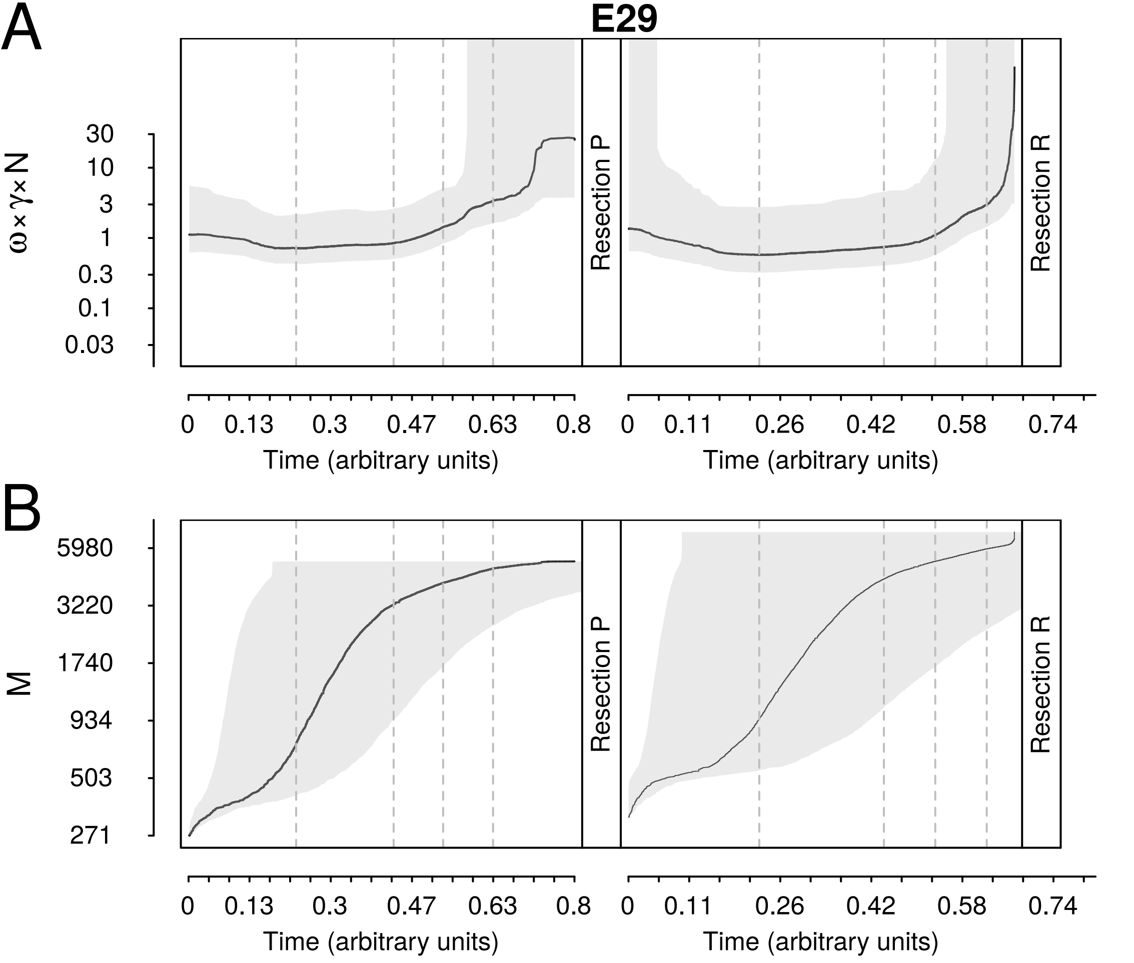
A
E29
ω×γ×N
30
10
3
1
0.3
0.1
0.03
Resection P
Resection R
0
0.13
0.3
0.47
0.63
0.8
Time (arbitrary units)
0
0.11
0.26
0.42
0.58
0.74
Time (arbitrary units)
B
M
5980
3220
1740
934
503
271
Resection P
Resection R
0
0.13
0.3
0.47
0.63
0.8
Time (arbitrary units)
0
0.11
0.26
0.42
0.58
0.74
Time (arbitrary units)

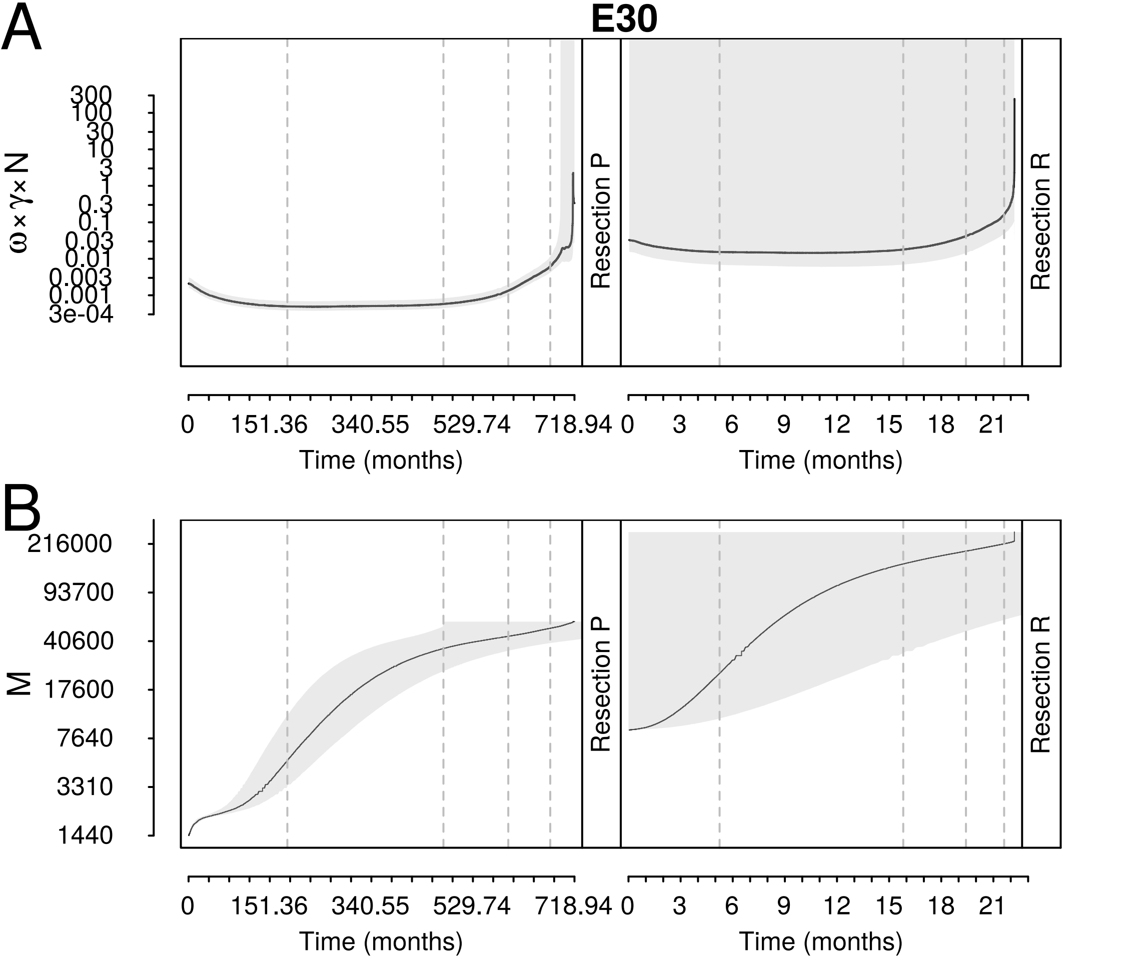
A
E30
ω×γ×N
300
100
30
10
3
1
0.3
0.1
0.03
0.01
0.003
0.001
3e-04
Resection P
Resection R
0
151.36
340.55
529.74
718.94
Time (months)
0
3
6
9
12
15
18
21
Time (months)
B
M
216000
93700
40600
17600
7640
3310
1440
Resection P
Resection R
0
151.36
340.55
529.74
718.94
Time (months)
0
3
6
9
12
15
18
21
Time (months)

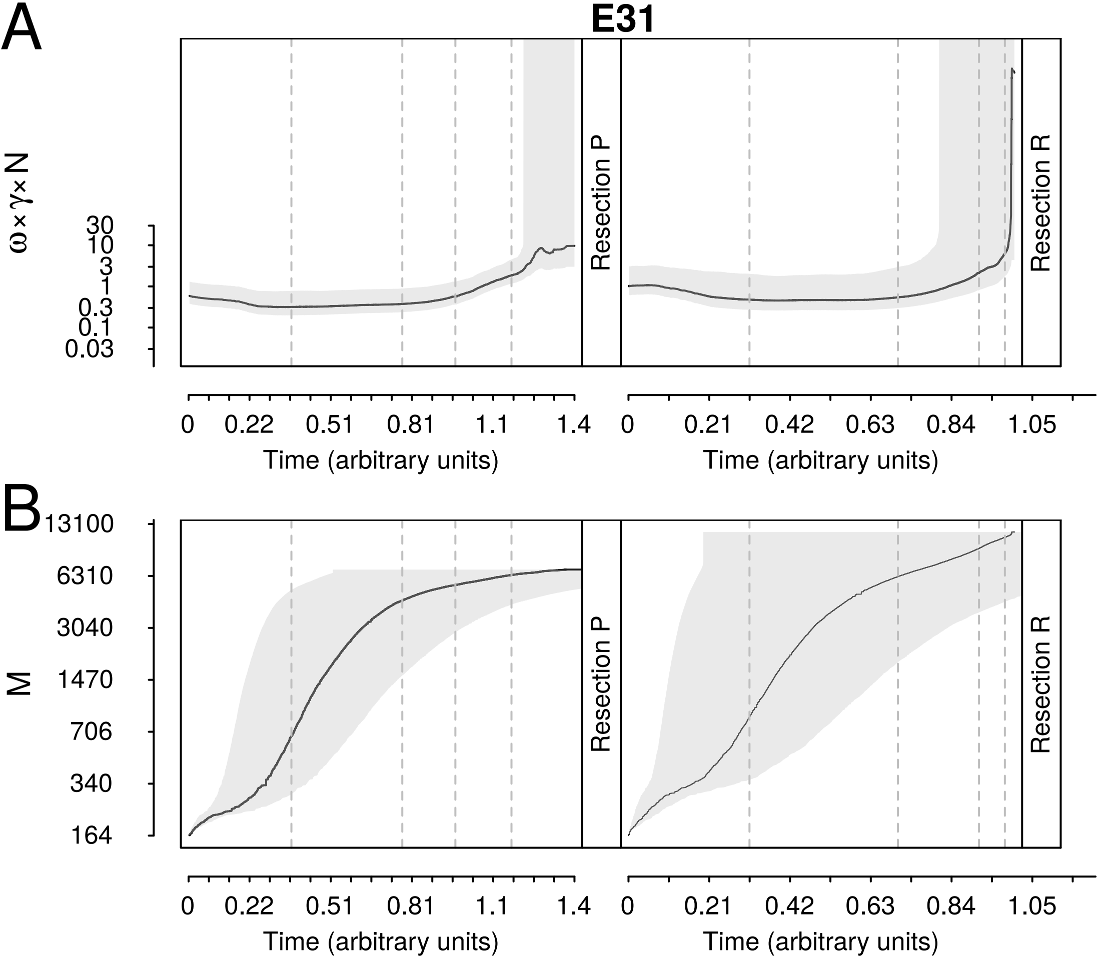
A
E31
ω×γ×N
30
10
3
1
0.3
0.1
0.03
Resection P
Resection R
0
0.22
0.51
0.81
1.1
1.4
Time (arbitrary units)
0
0.21
0.42
0.63
0.84
1.05
Time (arbitrary units)
B
M
13100
6310
3040
1470
706
340
164
Resection P
Resection R
0
0.22
0.51
0.81
1.1
1.4
Time (arbitrary units)
0
0.21
0.42
0.63
0.84
1.05
Time (arbitrary units)

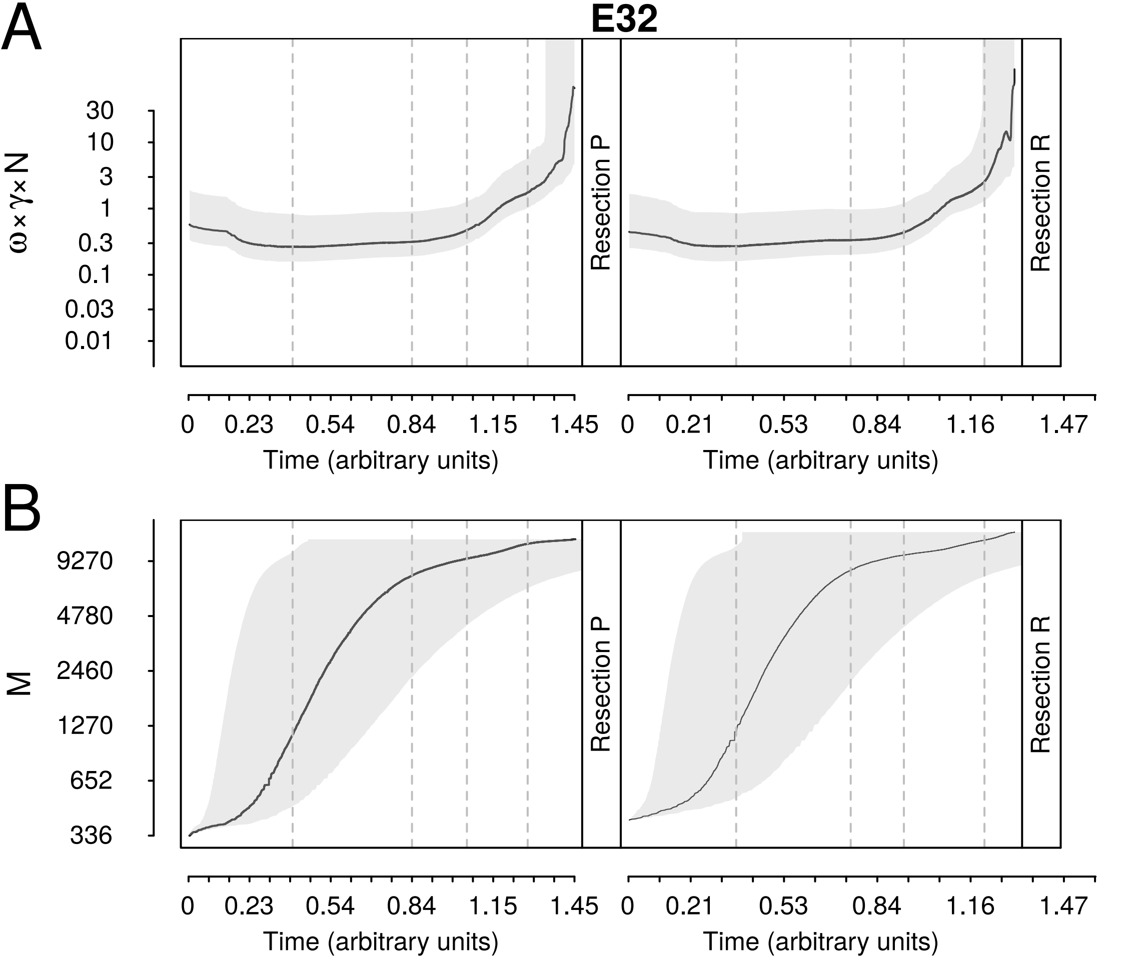

A
E32
ω×γ×N
30
10
3
1
0.3
0.1
0.03
0.01
Resection P
Resection R
0
0.23
0.54
0.84
1.15
1.45
0
0.21
0.53
0.84
1.16
1.47
Time (arbitrary units)
Time (arbitrary units)
B
M
9270
4780
2460
1270
652
336
Resection P
Resection R
0
0.23
0.54
0.84
1.15
1.45
0
0.21
0.53
0.84
1.16
1.47
Time (arbitrary units)
Time (arbitrary units)

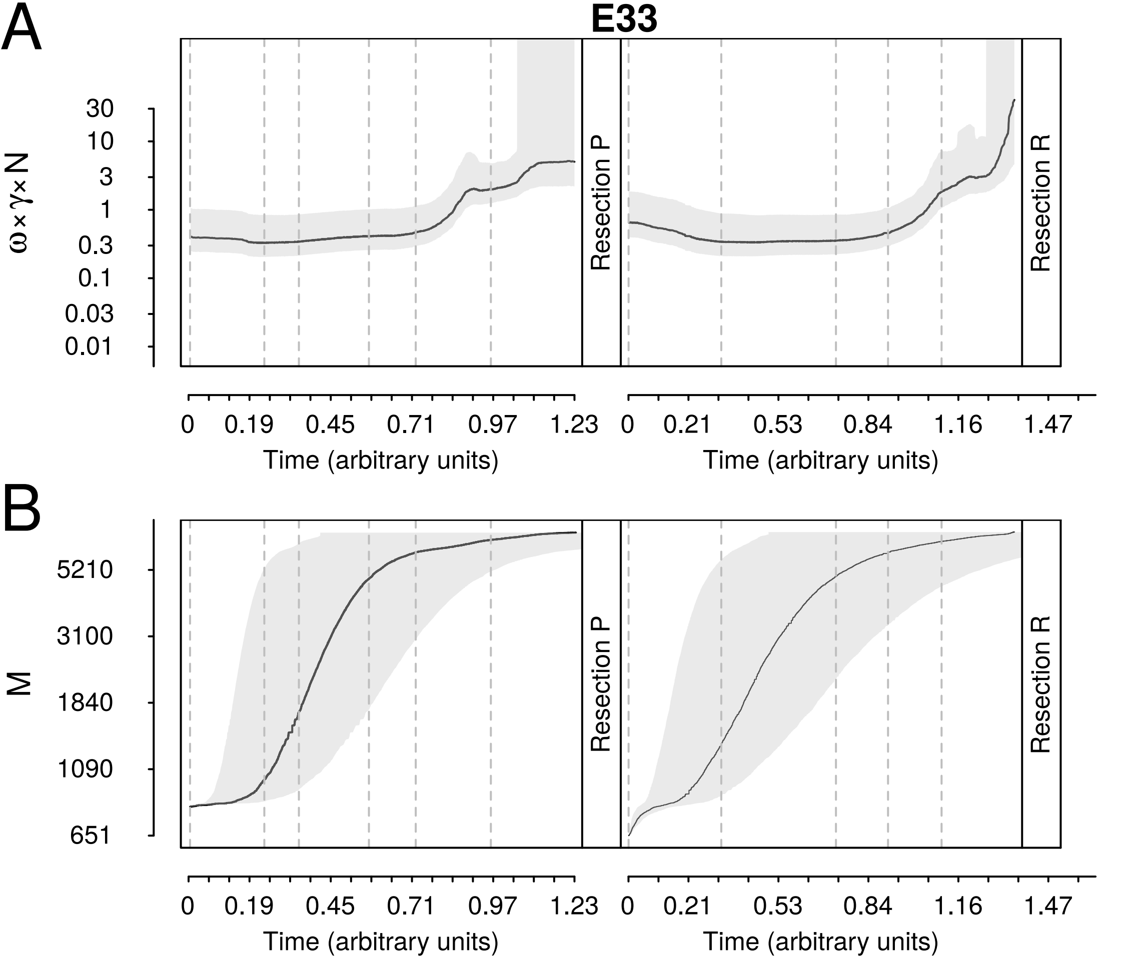
A
E33
ω×γ×N
30
10
3
1
0.3
0.1
0.03
0.01
Resection P
Resection R
0
0.19
0.45
0.71
0.97
1.23
Time (arbitrary units)
0
0.21
0.53
0.84
1.16
1.47
Time (arbitrary units)
B
M
5210
3100
1840
1090
651
Resection P
Resection R
0
0.19
0.45
0.71
0.97
1.23
Time (arbitrary units)
0
0.21
0.53
0.84
1.16
1.47
Time (arbitrary units)

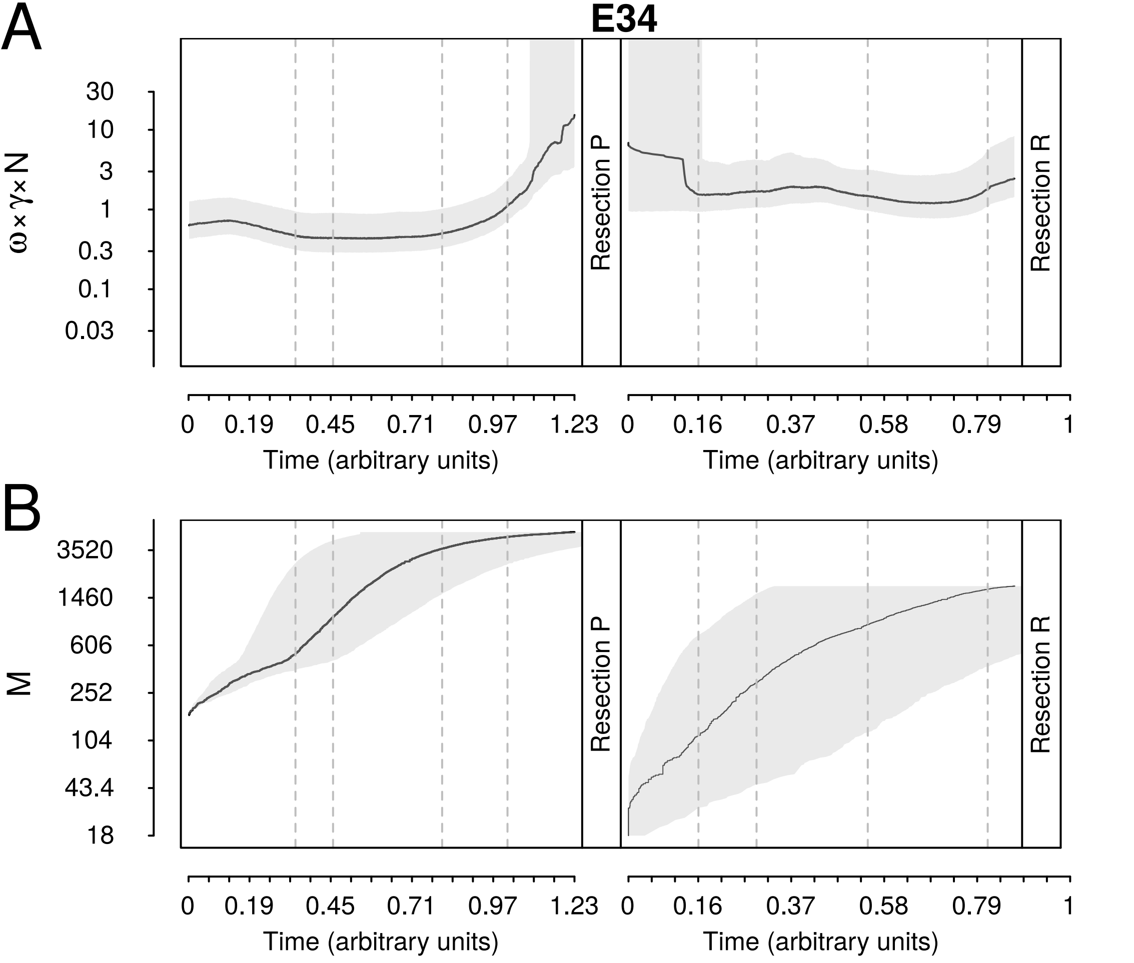

A
E34
30
10
3
1
0.3
0.1
0.03
ω×γ×N
Resection P
Resection R
0
0.19
0.45
0.71
0.97
1.23
Time (arbitrary units)
0
0.16
0.37
0.58
0.79
1
Time (arbitrary units)
B
3520
1460
606
252
104
43.4
18
M
Resection P
Resection R
0
0.19
0.45
0.71
0.97
1.23
Time (arbitrary units)
0
0.16
0.37
0.58
0.79
1
Time (arbitrary units)

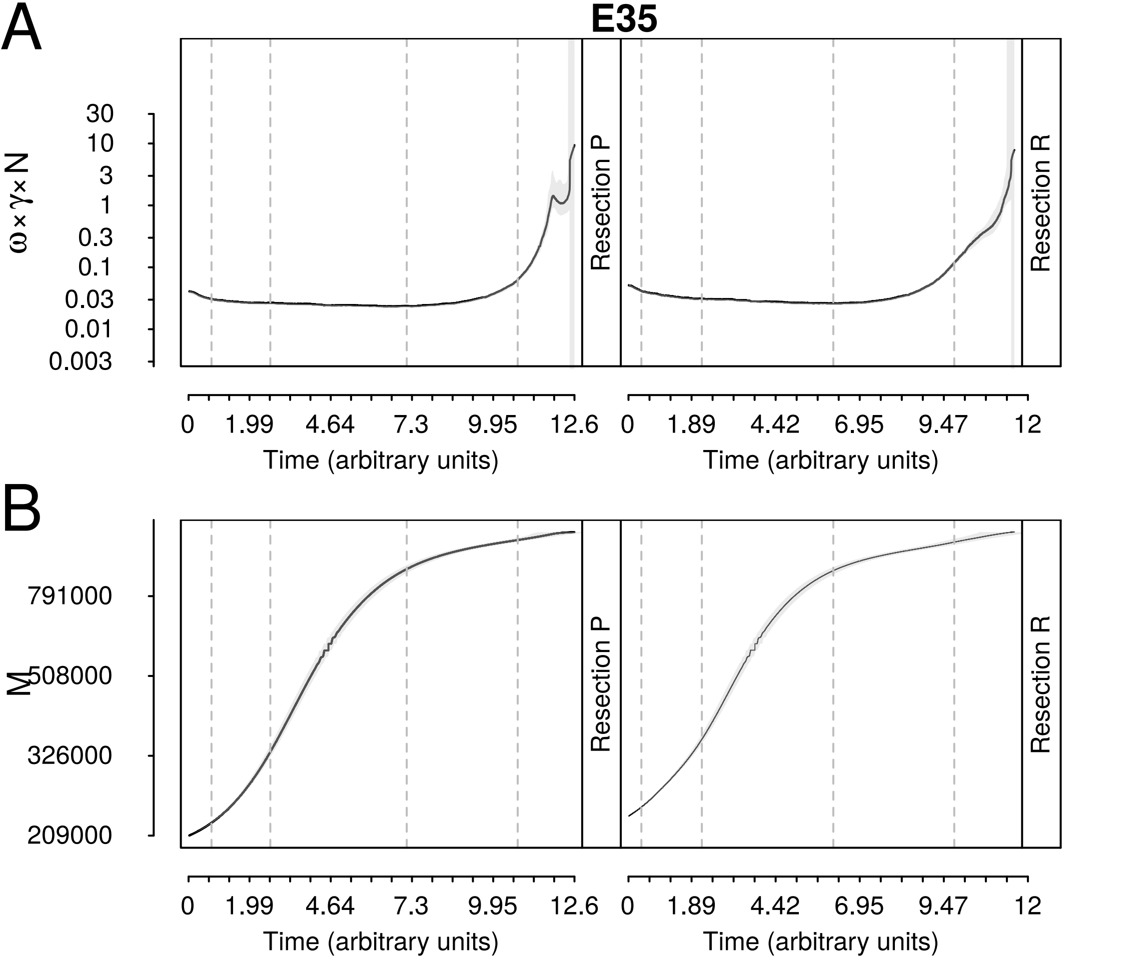
A
E35
ω×γ×N
30
10
3
1
0.3
0.1
0.03
0.01
0.003
Resection P
Resection R
0
1.99
4.64
7.3
9.95
12.6
Time (arbitrary units)
0
1.89
4.42
6.95
9.47
12
Time (arbitrary units)
B
M
791000
508000
326000
209000
Resection P
Resection R
0
1.99
4.64
7.3
9.95
12.6
Time (arbitrary units)
0
1.89
4.42
6.95
9.47
12
Time (arbitrary units)

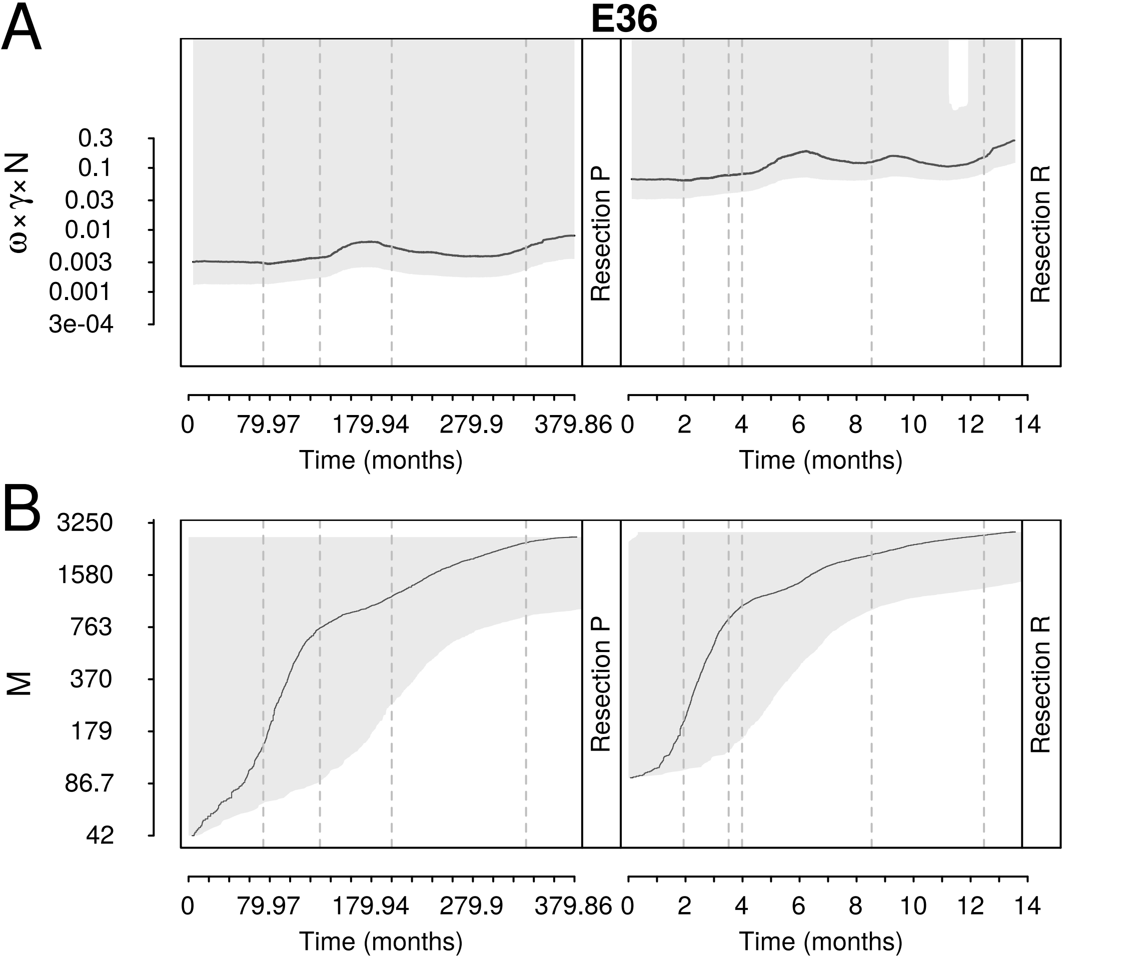

A
E36
ω×γ×N
0.3
0.1
0.03
0.01
0.003
0.001
3e-04
Resection P
Resection R
0
79.97
179.94
279.9
379.86
0
2
4
6
8
10
12
14
Time (months)
Time (months)
B
M
3250
1580
763
370
179
86.7
42
Resection P
Resection R
0
79.97
179.94
279.9
379.86
0
2
4
6
8
10
12
14
Time (months)
Time (months)

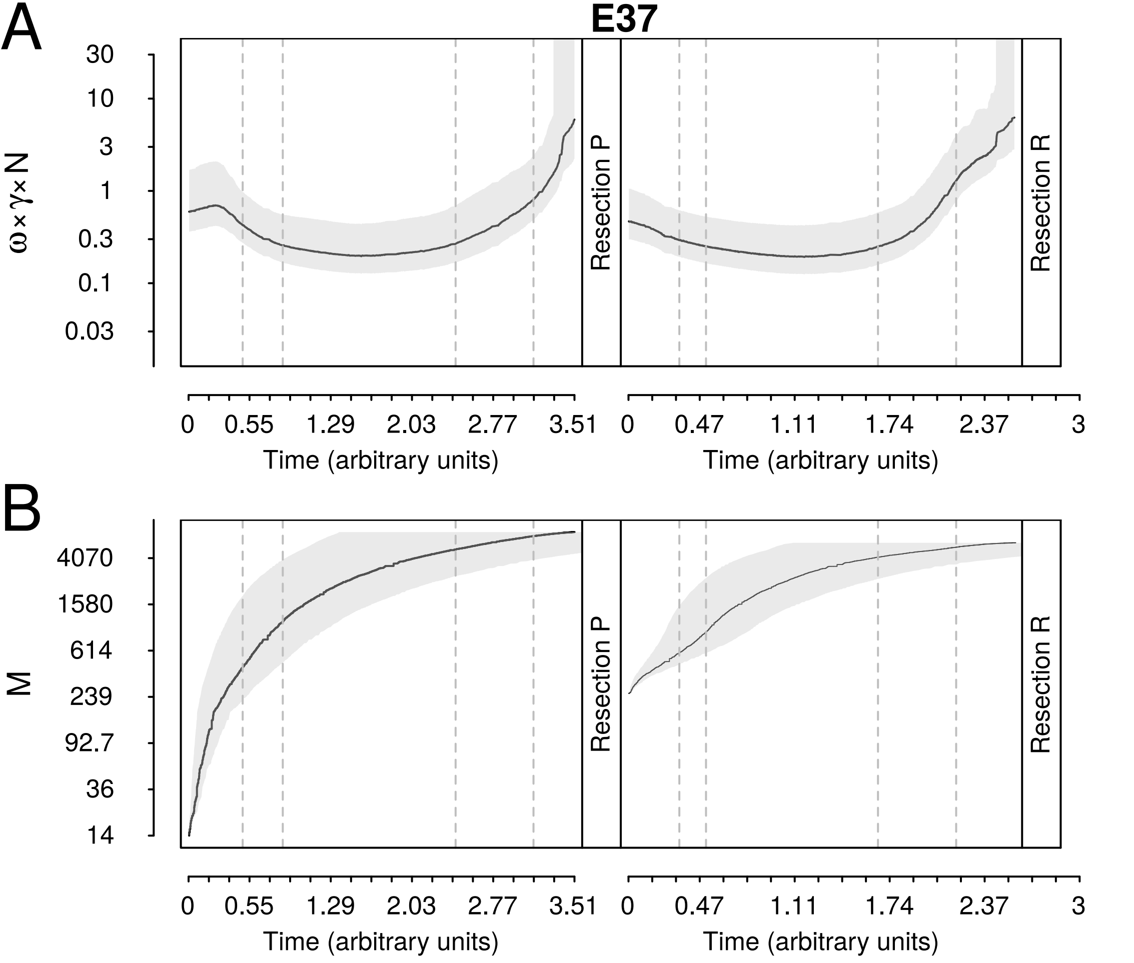

A
E37
30
10
3
1
0.3
0.1
0.03
ω×γ×N
Resection P
Resection R
0
0.55
1.29
2.03
2.77
3.51
Time (arbitrary units)
0
0.47
1.11
1.74
2.37
3
Time (arbitrary units)
B
4070
1580
614
239
92.7
36
14
M
Resection P
Resection R
0
0.55
1.29
2.03
2.77
3.51
Time (arbitrary units)
0
0.47
1.11
1.74
2.37
3
Time (arbitrary units)

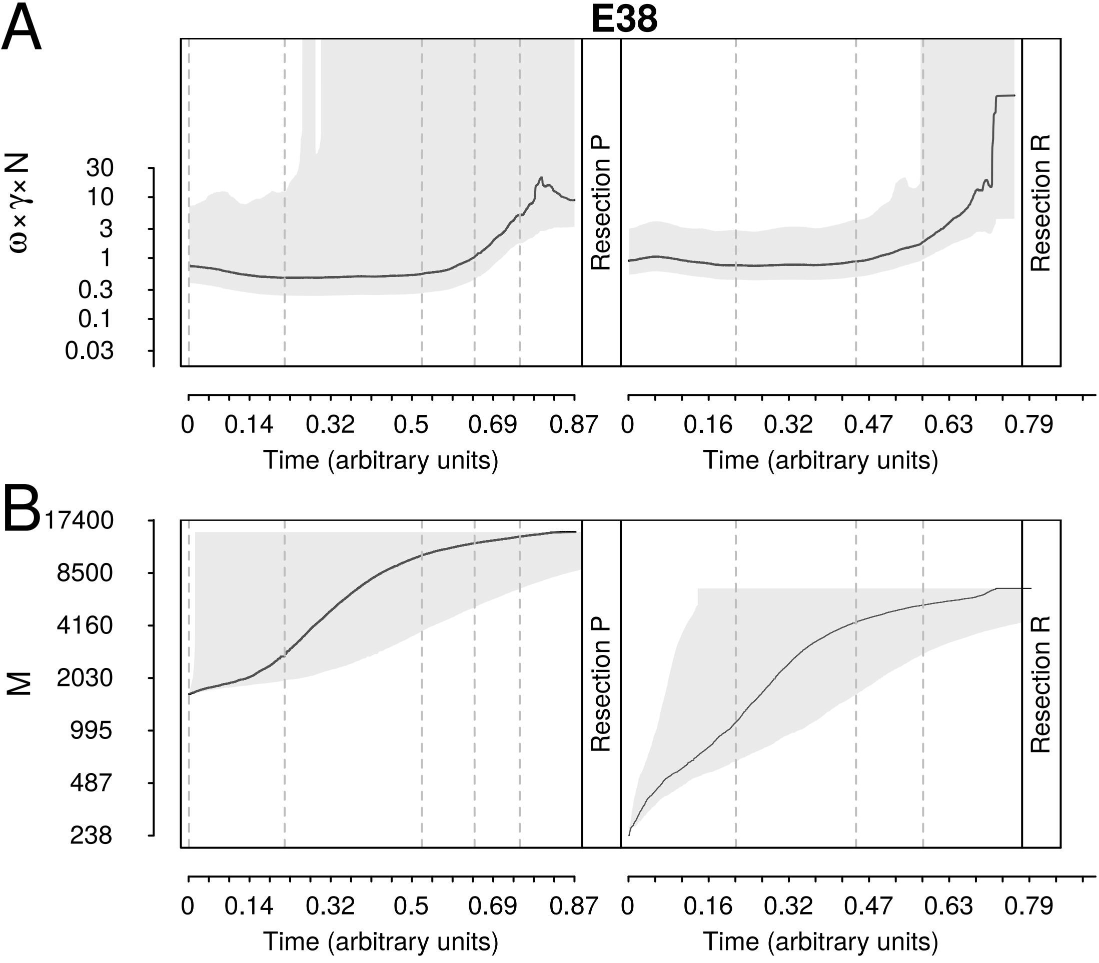
A
E38
ω×γ×N
30
10
3
1
0.3
0.1
0.03
Resection P
Resection R
0
0.14
0.32
0.5
0.69
0.87
Time (arbitrary units)
0
0.16
0.32
0.47
0.63
0.79
Time (arbitrary units)
B
M
17400
8500
4160
2030
995
487
238
Resection P
Resection R
0
0.14
0.32
0.5
0.69
0.87
Time (arbitrary units)
0
0.16
0.32
0.47
0.63
0.79
Time (arbitrary units)

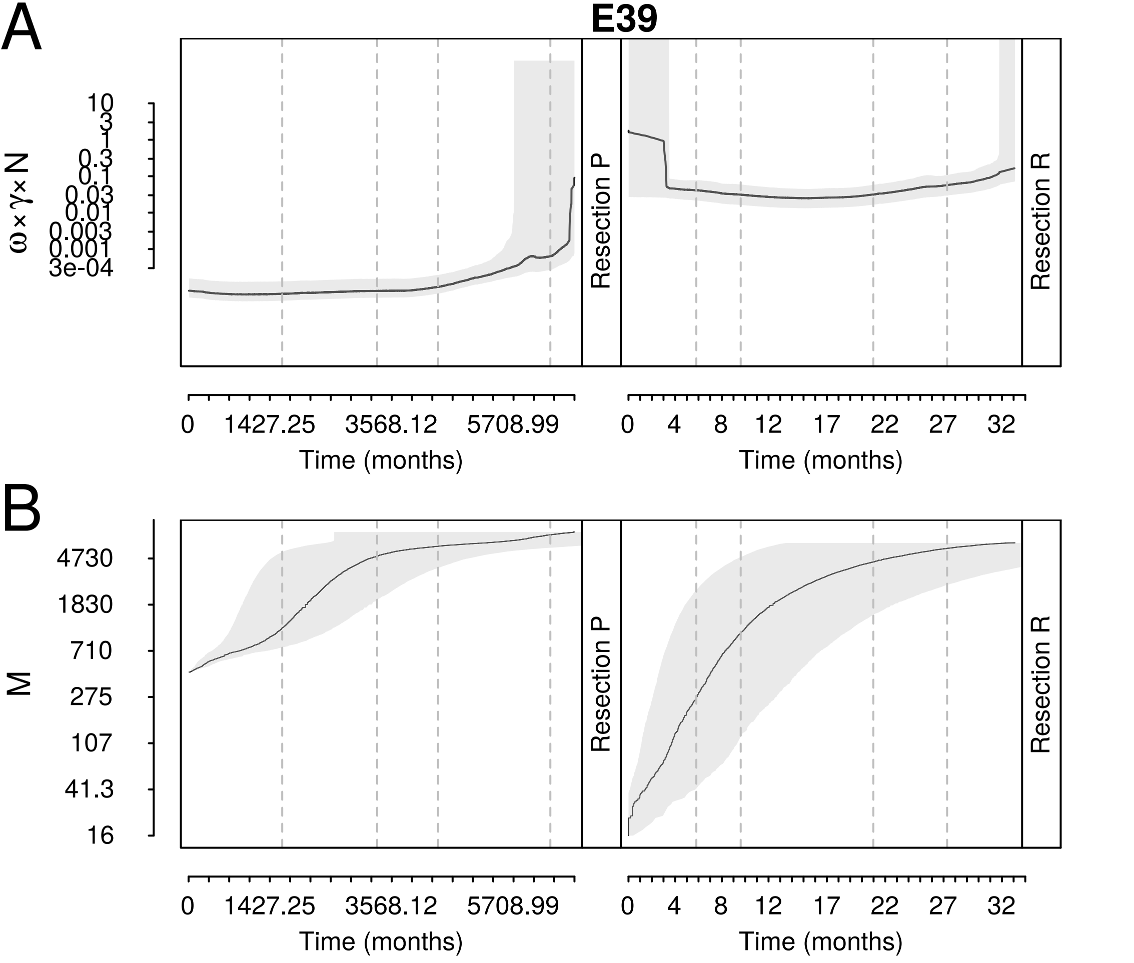

A
E39
ω×γ×N
10
3
1
0.3
0.1
0.03
0.01
0.003
0.001
3e-04
Resection P
Resection R
0
1427.25
3568.12
5708.99
Time (months)
0
4
8
12
17
22
27
32
Time (months)
B
M
4730
1830
710
275
107
41.3
16
Resection P
Resection R
0
1427.25
3568.12
5708.99
Time (months)
0
4
8
12
17
22
27
32
Time (months)

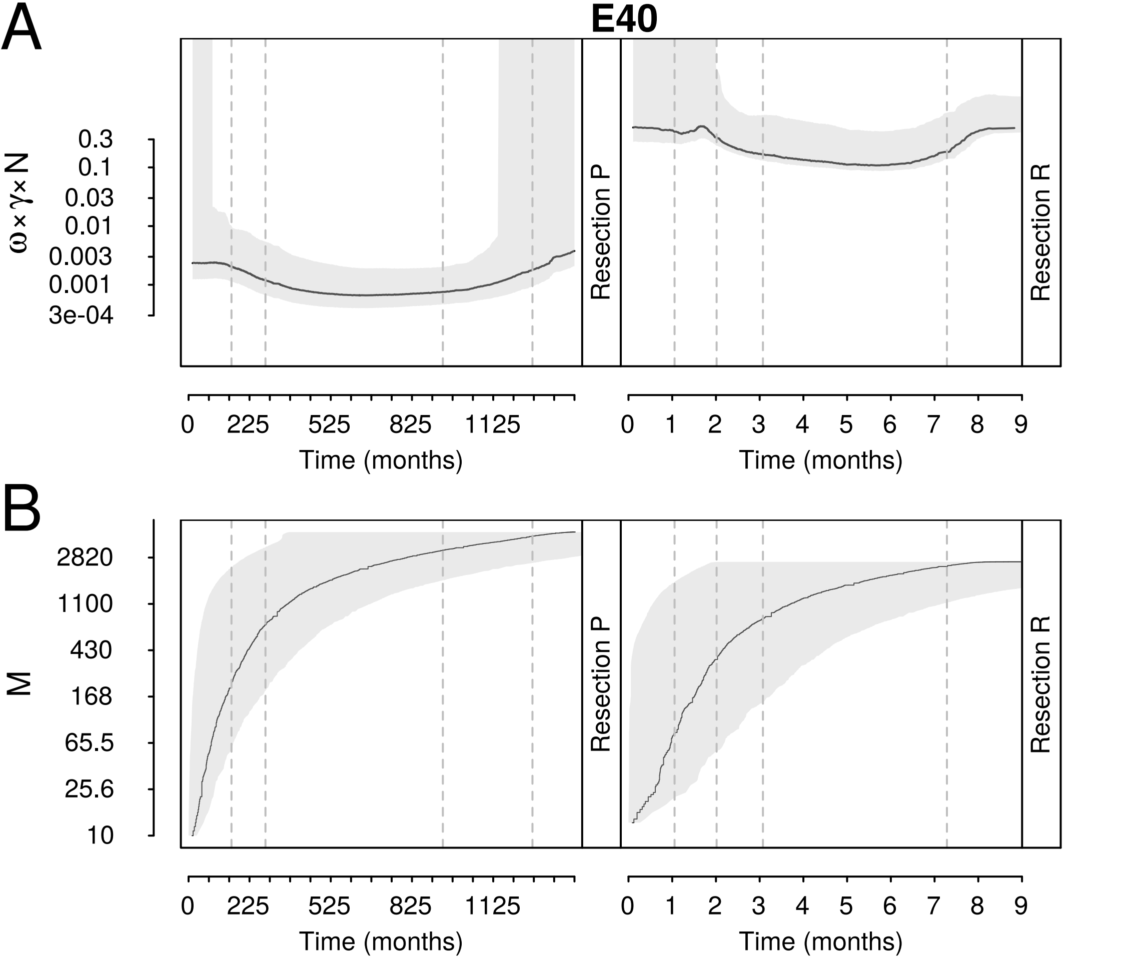

A
E40
ω×γ×N
0.3
0.1
0.03
0.01
0.003
0.001
3e-04
Resection P
Resection R
0
225
525
825
1125
Time (months)
0 1 2 3 4 5 6 7 8 9
Time (months)
B
M
2820
1100
430
168
65.5
25.6
10
Resection P
Resection R
0
225
525
825
1125
Time (months)
0 1 2 3 4 5 6 7 8 9
Time (months)

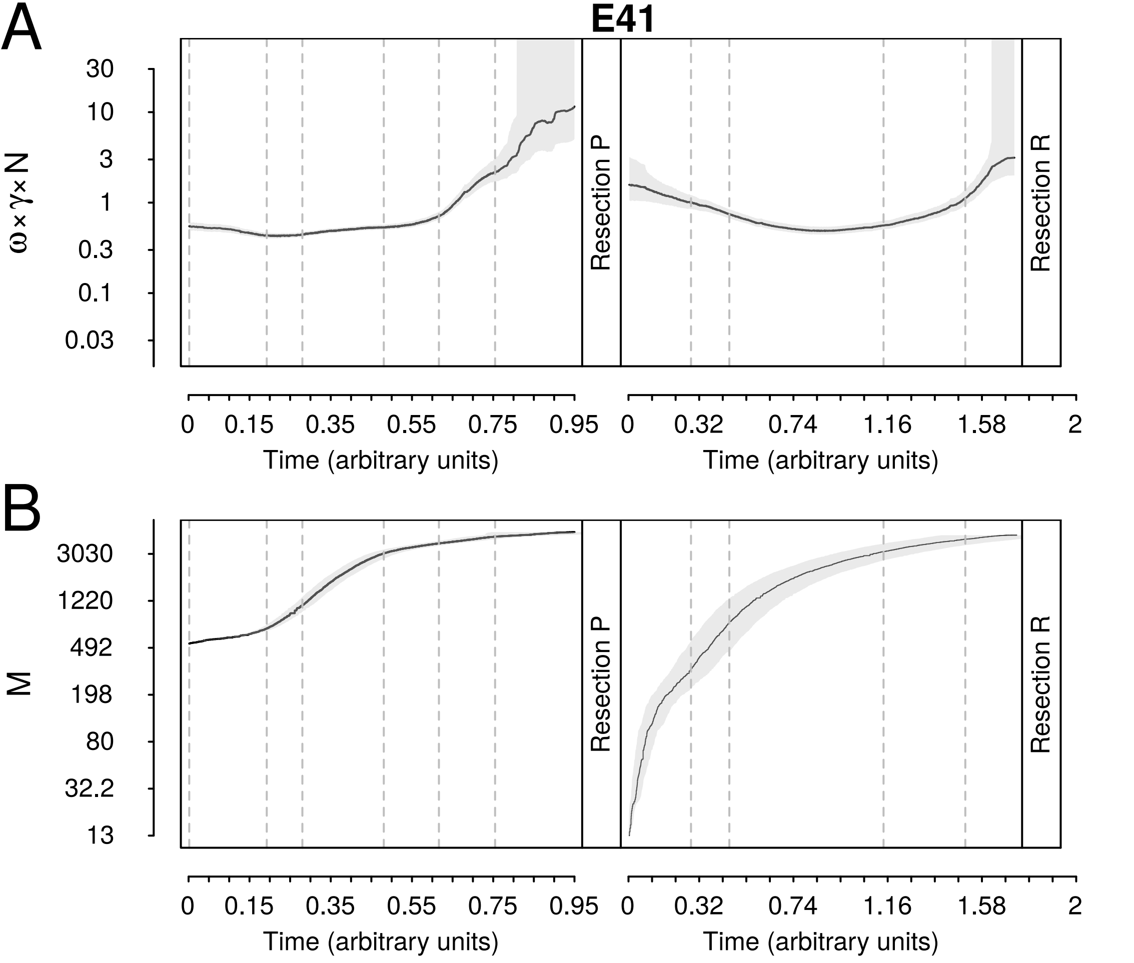

A
E41
30
10
3
1
0.3
0.1
0.03
ω×γ×N
Resection P
Resection R
0
0.15
0.35
0.55
0.75
0.95
Time (arbitrary units)
0
0.32
0.74
1.16
1.58
2
Time (arbitrary units)
B
3030
1220
492
198
80
32.2
13
M
Resection P
Resection R
0
0.15
0.35
0.55
0.75
0.95
Time (arbitrary units)
0
0.32
0.74
1.16
1.58
2
Time (arbitrary units)

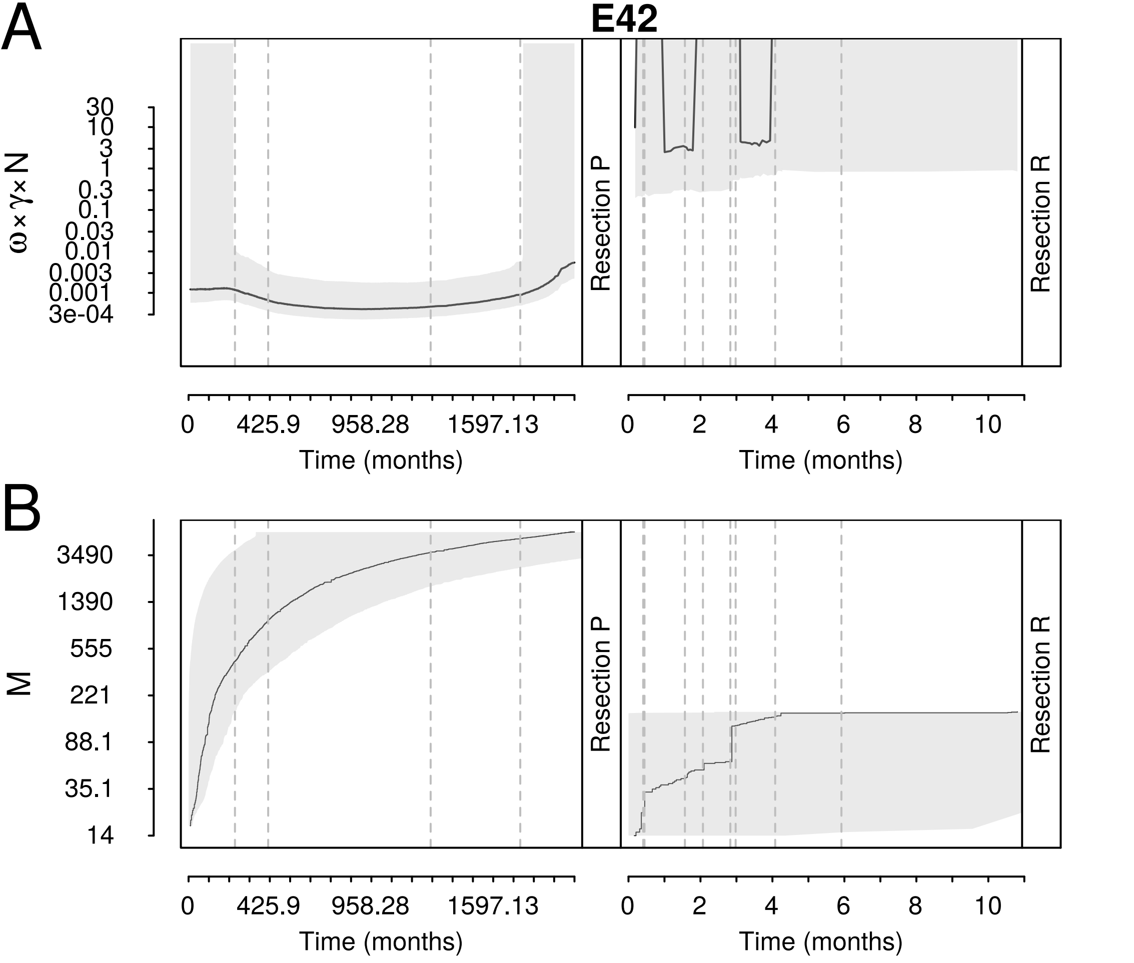
A
E42
ω×γ×N
30
10
3
1
0.3
0.1
0.03
0.01
0.003
0.001
3e-04
Resection P
Resection R
0
425.9
958.28
1597.13
Time (months)
0
2
4
6
8
10
Time (months)
B
M
3490
1390
555
221
88.1
35.1
14
Resection P
Resection R
0
425.9
958.28
1597.13
Time (months)
0
2
4
6
8
10
Time (months)
