## Supplementary Methods for "Time-resolved, integrated analysis of clonally evolving genomes"

#### Study of Mutation Accumulation

3.2 Mutation accumulation as a function of time for all subclones . . 5

### 1 Expressing mutation accumulation as a function of time

#### 1.1 Mutation accumulation with no subclones

The number of mutations at time $t$ is denoted $M(t)$. Taking relevant parameters in consideration, the increment of mutations $dM(t)$ at instant $t$ can be expressed as follows:

$$dM(t) = \mu(t) \cdot \pi(t) \cdot \omega(t) \cdot \gamma(t) \cdot N(t) \cdot dt \ ,$$

where :

- $\mu(t)$ is the mutation rate at instant $t$, per division, per haploid genome,
- $\pi(t)$ is the ploidy at instant $t$,
- $\omega(t)$ is the number of divisions per unit time, at instant $t$,
- $\gamma(t)$ is the survival rate, at instant $t$,
- $N(t)$ is the number of cells at instant $t$.

The number of divisions per unit time is scaled by the number of effectively surviving cells, through the use of survival rate $\gamma(t)$. This is because only surviving cells give rise to observable mutations.

#### 1.2 Mutation accumulation stratified by subclone

The expression in the latter section is limited to situations where cells are sufficiently homogeneous to be represented by a single set of parameters $\mu(t)$, $\pi(t)$, $\omega(t)$. This is seldom the case in tumors, where these parameters can vary from subclone to subclone or from cell to cell. Hence the increment of mutations is stratified by subclone $i$:

$$dM_i(t) = \mu_i(t) \cdot \pi_i(t) \cdot \omega_i(t) \cdot \gamma_i(t) \cdot N_i(t) \cdot dt \ , \quad (1)$$

where $i$ in $1, ..., n_c$ denotes a specific subclone, and $n_c$ is the number of subclones.

### 2 Expressing allele frequency as a function of time

#### 2.1 Number of cells and allele frequency

The number of cells, $N_i(t)$ can be expressed as a function of allele frequency $f_i(t)$ [1], when one considers that a mutation appearing in subclone $i$ at instant $t$ must correspond to one mutation among $N_i(t) \cdot \pi_i(t)$ chromosome copies, at that specific instant $t$. This can be written as follows:

$$f_i(t) = \frac{1}{N_i(t) \cdot \pi_i(t)} \ . \tag{2}$$

However, allele frequency is observed not only in subclone $i$, but over all subclones. Furthermore, allele frequency is observed not when the mutation occurred, but later at sample retrieval or resection time point, $\tau_r$. It should be noted that $\tau_r$ is a constant.

Therefore we have to introduce a generalized version of equation (2.1), by using the observed allele frequency $f_i(t; \tau_r)$, for a mutation appeared in subclone $i$ at time $t$ and observed at time $\tau_r$ (resection time or sampling time).

To obtain the formulation of $f_i(t; \tau_r)$, we make the following assumptions :

- If a mutation is under selection, the subclone part containing the mutation is considered as an additional subclone. Inside of a subclone, selection is considered to be negligible.
- Growth is considered high enough to make random genetic drift negligible [2, 3, 4]. This allows to consider a constant allele frequency between mutation occurrence and sampling and sequencing events.
- Finally, considering that ploidy changes might compensate themselves, and that large changes can be detected and analysed separately, we also make the assumption that ploidy $\pi$ is constant.

Considering the above assumptions, the observed allele frequency $f_i(t; \tau_r)$ may be expressed as follows:

$$f_i(t; \tau_r) = \frac{1}{N_i(t) \cdot \pi} \cdot \frac{N_i(\tau_r)}{\sum\limits_j N_j(\tau_r)} \tag{3}$$

$$\iff N_i(t) = \frac{1}{f_i(t; \tau_r) \cdot \pi} \cdot \frac{N_i(\tau_r)}{\sum\limits_j N_j(\tau_r)} \ , \tag{4}$$

where $j$ denotes a subclone, and maybe equal or different of $i$ in 1,2,...,$n_c$, and $\pi$ is assumed constant.

The factor term $N_i(\tau_r)/\sum_j N_j(\tau_r)$ is denoted $K_{i,r}$ in the following. This factor corresponds to the proportion of subclone $i$ among all subclones. $K_{i,r}$ is constant since it depends only on constant values.

Expressions (2.1) and (2.1) for the observed allele frequency and for number of cells are rewritten using $K_{i,r}$:

$$f_i(t; \tau_r) = \frac{K_{i,r}}{N_i(t) \cdot \pi} \tag{5}$$

$$\iff N_i(t) = \frac{K_{i,r}}{f_i(t; \tau_r) \cdot \pi} \ .$$

Finally, observed allele frequency $f_i(t; \tau_r)$ is comparable between subclones. As a result, it is possible to denote this frequency simply $f$, to represent the allele frequency observed at sampling time over all subclones, with $t$ taken into account via $K_{i,r}$, and with $\tau_r$, which is the same for all subclones, implied. For clarity we continue to use $f_i(t;\tau_r)$ when looking at a single subclone $i$, and we use $f$ when summing over all subclones (section 3.2).

#### 2.2 Number of cells and time

The new cells are generated via a certain number of divisions per time unit, equal to $\omega_i(t)$ and, since not all new cells survive, this number is modulated by the survival rate $\gamma_i(t)$. As a consequence, $Ni(t+dt)$ can be expressed as follows:

$$N_i(t+dt) = N_i(t) + \omega_i(t) \cdot \gamma_i(t) \cdot N_i(t) \cdot dt \ . \tag{6}$$

An increment of cells $dN_i$ appeared during time increment $dt$ is defined as follows:

$$dN_i = N_i(t+dt) - N_i(t) \ .$$

Consequently, $dNi$ can be written as a function of $dt$, using (2.2):

$$dN_i = \omega_i(t) \cdot \gamma_i(t) \cdot N_i(t) \cdot dt \ . \tag{7}$$

#### 2.3 Allele frequency as a function of time

From here, we use inverse allele frequency $1/f_i(t;\tau_r)$ and its increment $d(1/f_i(t;\tau_r))$, instead of $f_i(t;\tau_r)$ and $df_i(t;\tau_r)$, because it is equivalent and yields simpler formulations. From the above expression (2.1), $1/f_i(t;\tau_r)$ can be expressed as a function of $N_i(t)$:

$$(1/f_i)(t;\tau_r) = \frac{N_i(t) \cdot \pi}{K_{i,r}} \ .$$

Doing the same at $t+dt$ and substracting the expression at $t$ from the expression at $t+dt$ yields the increment:

$$\begin{aligned} d(1/f_i(t;\tau_r)) &= (1/f_i)(t+dt;\tau_r) - (1/f_i)(t;\tau_r) \\ &= \frac{N_i(t+dt) \cdot \pi}{K_{i,r}} - \frac{N_i(t) \cdot \pi}{K_{i,r}} \\ &= \frac{(N_i(t+dt) - N_i(t)) \cdot \pi}{K_{i,r}} \\ &= \frac{dN_i \cdot \pi}{K_{i,r}} \ . \end{aligned}$$

Then, $dN_i$ can be replaced by its expression (2.2), leading to the following:

$$d(1/f_i(t;\tau_r)) = \frac{\omega_i(t) \cdot \gamma_i(t) \cdot N_i(t) \cdot dt \cdot \pi}{K_{i,r}} \ . \tag{8}$$

These equations, as well as those for the number of cells dynamics, are summarized in table 1.

**Table 1:** Dynamics of $N_i(t)$ and $(1/f)(t)$.

| | Value at $t$ | Value at $t + dt$ | Increment during $dt$ |
|---|---|---|---|
| $N_i$ | $N_i(t)$ | $N_i(t + dt)$ $=$ $N_i(t) + \omega_i(t)\,\gamma_i(t)\,N_i(t)\,dt$ | $N_i(t + dt) - N_i(t) = dN_i$ $=$ $\omega_i(t)\,\gamma_i(t)\,N_i(t)\,dt$ |
| $1/f$ | $(1/f_i)(t;\tau_r)$ $=$ $\frac{N_i(t)\,\pi}{K_{i,r}}$ | $(1/f_i)(t + dt;\tau_r)$ $=$ $\frac{N_i(t+dt)\,\pi}{K_{i,r}}$ | $(1/f_i)(t + dt;\tau_r) - (1/f_i)(t;\tau_r) = d(1/f)(t)$ $=$ $\frac{(N_i(t+dt)-N_i(t))\,\pi}{K_{i,r}} = \frac{dN_i \cdot \pi}{K_{i,r}}$ $=$ $\frac{\pi\,\omega_i(t)\,\gamma_i(t)\,N_i(t)\,dt}{K_{i,r}}$ |

### 3 Expressing mutation accumulation as a function of allele frequency

#### 3.1 Mutation accumulation stratified by subclone

Using equation (2.3), one can reformulate mutation accumulation from (1.2) as a function of $d(1/f)$:

$$dM_i(t) = \mu_i(t) \cdot \pi \cdot \omega_i(t) \cdot \gamma_i(t) \cdot N_i(t) \cdot dt \cdot \frac{d(1/f_i)(t;\tau_r) \cdot K_{i,r}}{\omega_i(t) \cdot \gamma_i(t) \cdot N_i(t) \cdot dt \cdot \pi} \ . \quad (9)$$

Equation (3.1) simplifies into the following:

$$dM_i(t) = \mu_i(t) \cdot K_{i,r} \cdot d(1/f_i)(t;\tau_r) \ . \quad (10)$$

#### 3.2 Mutation accumulation as a function of time for all subclones

Considering all subclones, mutation accumulation is simply the sum over subclones 1, ... , $i$, ... , $n_c$ of the mutation increment in each subclone:

$$dM(t) = \sum_{i=1}^{n_c} dM_i(t) \ , \quad (11)$$

Using (3.1) and (3.2) yields mutation accumulation as a function of the inverse allele frequency (3.2). As mentioned in section *2.1*, $f$ is the observed allele frequency and is identical from one subclone to the other, while discrepancies between subclones are represented by constant $K_{i,r}$. Mutation accumulation over all subclones can then be written as follows:

$$dM(t) = \left( \sum_{i=1}^{n_c} \mu_i(t) \cdot K_{i,r} \right) \cdot d(1/f) \ . \quad (12)$$

From equation (3.2), mutation accumulation $dM$ is proportional to $d(1/f)$. This relation is modulated by a weighted sum over the mutation rates in subclones $i$, where the weights are the $K_{i,r}$ terms. Since the weights $K_{i,r}$ do not depend on time, this means that constant mutation rates lead to a linear relationship between $dM$ and $d(1/f)$, regardless of the presence or not of natural selection.

Conversely, it is not possible to say that linearity between $dM$ and $d(1/f)$ implies constant mutation rates, since these terms could in principle exactly compensate each other to form the same constant sum $\sum_i \mu_i(t) \cdot K_{i,r}$. However, this might be considered unlikely.

As a result, one can write the following equivalence, implication and consideration:

$$\left\{ \sum_i \mu_i(t) \cdot K_{i,r} \textit{ is constant} \right\} \iff \{M(1/f) \textit{ is linear}\},$$

$$\{\mu_i(t) \textit{ is constant } \forall i\} \Rightarrow \{M(1/f) \textit{ is linear}\}$$

and

$$\{M(1/f) \textit{ is linear}\} \textit{ possibly implies} : \{\mu_i(t) \textit{ is constant } \forall i\}.$$

As a consequence, equation (3.2) also means that the profile of $M$ as a function of $1/f$ corresponds to a sum of subclones at each $1/f$. As a consequence, any nonlinearity in this curve will reflect mutation rate dynamics happening in one (or several) subclone(s), and does not correspond to a single subclone.
