## Supplementary Tables for "Time-resolved, integrated analysis of clonally evolving genomes"

**This PDF file includes:**

Supplementary Tables 1-3

**<u>Supplementary Table 1.</u>** *P-values* for test of difference between segments for P. virginalis and glioblastoma samples.

| **Sample** | ***p-value*** **(for primary tumor for glioblastoma samples)** | ***p-value*** **for recurrent tumor (for glioblastoma samples)** |
|---|---|---|
| P.virginalis | $6.345498x10^{-2}$ | not applicable |
| GBM1 | $<2.2x10^{-16}$ | $<2.2x10^{-16}$ |
| GBM2 | $<2.2x10^{-16}$ | $<2.2x10^{-16}$ |
| GBM3 | $<2.2x10^{-16}$ | $<2.2x10^{-16}$ |
| GBM4 | $<2.2x10^{-16}$ | $<2.2x10^{-16}$ |
| GBM5 | $<2.2x10^{-16}$ | $<2.2x10^{-16}$ |
| GBM6 | $<2.2x10^{-16}$ | $<2.2x10^{-16}$ |
| GBM7 | $<2.2x10^{-16}$ | $<2.2x10^{-16}$ |
| GBM8 | $<2.2x10^{-16}$ | $<2.2x10^{-16}$ |
| GBM9 | $<2.2x10^{-16}$ | $<2.2x10^{-16}$ |
| GBM10 | $<2.2x10^{-16}$ | $<2.2x10^{-16}$ |
| GBM11 | $<2.2x10^{-16}$ | $<2.2x10^{-16}$ |
| GBM12 | $<2.2x10^{-16}$ | $<2.2x10^{-16}$ |
| GBM13 | $<2.2x10^{-16}$ | $<2.2x10^{-16}$ |
| GBM14 | $<2.2x10^{-16}$ | $<2.2x10^{-16}$ |
| GBM15 | $7.962633\ x10^{-15}$ | $<2.2x10^{-16}$ |
| GBM16 | $<2.2x10^{-16}$ | $<2.2x10^{-16}$ |
| GBM17 | $<2.2x10^{-16}$ | $7.891463\ x10^{-01}$ |
| GBM18 | $<2.2x10^{-16}$ | $<2.2x10^{-16}$ |
| GBM19 | $<2.2x10^{-16}$ | $<2.2x10^{-16}$ |
| GBM20 | $<2.2x10^{-16}$ | $<2.2x10^{-16}$ |
| GBM21 | $<2.2x10^{-16}$ | $<2.2x10^{-16}$ |
| GBM22 | $<2.2x10^{-16}$ | $1.883298\ x10^{-03}$ |
| GBM23 | $<2.2x10^{-16}$ | 1.000000 |
| GBM24 | $<2.2x10^{-16}$ | $4.886992\ x10^{-11}$ |
| GBM25 | $<2.2x10^{-16}$ | $2.928589\ x10^{-02}$ |
| GBM26 | $<2.2x10^{-16}$ | $<2.2x10^{-16}$ |
| GBM27 | $<2.2x10^{-16}$ | $<2.2x10^{-16}$ |
| GBM28 | $<2.2x10^{-16}$ | $<2.2x10^{-16}$ |
| GBM29 | $<2.2x10^{-16}$ | $<2.2x10^{-16}$ |
| GBM30 | 0.000000 | $<2.2x10^{-16}$ |
| GBM31 | $<2.2x10^{-16}$ | $<2.2x10^{-16}$ |
| GBM32 | $<2.2x10^{-16}$ | $<2.2x10^{-16}$ |
| GBM33 | $<2.2x10^{-16}$ | $<2.2x10^{-16}$ |
| GBM34 | $<2.2x10^{-16}$ | $<2.2x10^{-16}$ |
| GBM35 | $<2.2x10^{-16}$ | $<2.2x10^{-16}$ |
| GBM36 | $<2.2x10^{-16}$ | $<2.2x10^{-16}$ |
| GBM37 | $<2.2x10^{-16}$ | $<2.2x10^{-16}$ |
| GBM38 | $<2.2x10^{-16}$ | $<2.2x10^{-16}$ |
| GBM39 | $<2.2x10^{-16}$ | $<2.2x10^{-16}$ |
| GBM40 | $<2.2x10^{-16}$ | $<2.2x10^{-16}$ |
| GBM41 | $<2.2x10^{-16}$ | $<2.2x10^{-16}$ |
| GBM42 | $<2.2x10^{-16}$ | $5.637243\ x10^{-04}$ |

**Supplementary Table 2.** Characteristics of tumor cell survival ratio $\gamma_R/\gamma_P$ (n=20). Index R denotes the start of the recurrence and P the end of the primary. The lower and higher bounds on $\gamma_R/\gamma_P$ correspond to tumor emergence 2 and 7 years before diagnosis, respectively.

| | log $\gamma_R/\gamma_P$ | |
|---|---|---|
| **sample** | **lower bound** | **higher bound** |
| 1 | 1.9782320 | 2.5223001 |
| 4 | 3.3990271 | 3.9430951 |
| 6 | 1.3989074 | 1.9429809 |
| 7 | 1.5080083 | 2.0520764 |
| 8 | 1.2397434 | 1.7838077 |
| 10 | 1.4497716 | 1.9938336 |
| 11 | 3.2291480 | 3.7746907 |
| 14 | 1.4457411 | 1.9898273 |
| 15 | 1.2063772 | 1.7504348 |
| 18 | 1.4437458 | 1.9877957 |
| 21 | 1.6193366 | 2.1634227 |
| 23 | 2.2465702 | 2.7907532 |
| 24 | 1.3218368 | 1.8659186 |
| 27 | 2.3521282 | 2.8961963 |
| 28 | 1.8456369 | 2.3896593 |
| 30 | 1.0004823 | 1.5445460 |
| 36 | 1.2751182 | 1.8191862 |
| 39 | 0.0260095 | 0.5700782 |
| 40 | 0.7027364 | 1.2468078 |
| 42 | 0.5513292 | 1.0953980 |
| **mean** | 1.561994 | 2.106140 |
| **95% CI** | [1.250896 ; 1.947115] | [1.795669 ; 2.490084] |
| **median** | 1.444743 | 1.988812 |
| **95% CI** | [1.240748 ; 1.732487] | [1.784811 ; 2.276541] |

**<u>Supplementary Table 3.</u>** *Procambarus virginalis* Samples

| **Name** | **Date of birth (y)** | **Uncertainty (y)** | **Name in Gutekunst *et al.* 2018** |
|---|---|---|---|
| Animal 1 | 2012.12500 | 0.166667 | Petshop 1 laboratory strain |
| Animal 2 | 2014.87500 | 0.083333 | Petshop 2 laboratory strain |
| Animal 34 | 2017.87500 | 0.083333 | n.a. |
| Animal 35 | 2018.20833 | 0.083333 | n.a. |
| Hannover | 2016.95833 | 0.083333 | Hannover aquarium lineage |
| Heidelberg | 2010.00000 | 2.000000 | Heidelberg laboratory strain |
| Madagascar 1 | 2011.00000 | 1.000000 | MA1 |
| Madagascar 2 | 2015.25000 | 0.250000 | MA2 |
| Madagascar 3 | 2015.58333 | 0.166667 | MA3 |
| Madagascar 4 | 2015.58333 | 0.250000 | MA4 |
| Madagascar 5 | 2015.58333 | 0.166667 | MA5 |
| Moosweiher | 2011.00000 | 1.000000 | Moosweiher |
| Reilingen | 2015.00000 | 1.000000 | Reilingen |

n.a. : not applicable
